# Parameters for stable Notch-independent Her6 oscillations across a frequency spectrum in neural stem cell populations

**DOI:** 10.64898/2026.09.15.751824

**Authors:** Christian Sigloch, Adrian L. Hauber, Jan Rothenpieler, Samuel Böhm, Dominik Spitz, Roland Nitschke, Jens Timmer, Wolfgang Driever

## Abstract

Neural proliferation zones drive strong and persistent growth of the developing vertebrate brain. How Notch signaling and HES/Her transcription factors *in* vivo maintain stable neural stem cell (NSC) populations during brain growth is not well understood. Genetic evidence revealed Her6, a Notch-independent homolog of mammalian HES1, to be essential for maintenance of zebrafish NSCs. We recombined mNeonGreen into the *her6* locus and analyzed temporal Her6-mNG fluorescence profiles *in vivo*. Her6 is expressed dynamically in all NSCs, with a majority revealing stable oscillation, albeit from cell to cell across a wide frequency range. Her6 oscillates cell-autonomously, unaffected by inhibition of Notch, and is robustly reinitiated after experimental perturbations. Mathematical modelling predicts, and we indeed find *in vivo*, that Her6 oscillations prevail only when both transcript and protein degradation rate constants are equal. These intrinsic control parameters may have evolved for the Her6 oscillator to contribute to robust maintenance of NSCs.

## Introduction

Neural proliferation zones (NPZs) harbor long-term neural stem cells (NSCs) and drive massive brain growth during vertebrate development(Paridaen and Huttner, 2014), while in adult neural stem cell niches NSC proliferation is more restricted(Kriegstein and Alvarez-Buylla, 2009; Paridaen and Huttner, 2014; Urban and Guillemot, 2014; Gotz et al., 2016; Obernier and Alvarez-Buylla, 2019). In each environment, NSCs must achieve a balance between stem cell maintenance, proliferation, and exit into differentiation. Notch signaling is a key regulator of NSCs and neural progenitor cell (NPC) lineage progression(Chitnis et al., 1995; Shimojo et al., 2008; Lampada and Taylor, 2023). Notch activates expression of *HES/Her* genes(Ohtsuka et al., 1999), transcription factors that play a crucial role in neurogenesis by repressing differentiation(Imayoshi and Kageyama, 2014; Kageyama et al., 2015). In mice, *Hes1* and *Hes5* expression are both activated by Notch signaling(Ohtsuka et al., 1999), but *Hes1* also displays Notch-independent expression(de la Pompa et al., 1997; Kageyama et al., 2007). Notch-independent HES/Her activities execute lineage-based cell autonomous regulation of NSCs, while Notch-dependent HES/Her activities reflect local juxtacrine Notch signaling within the ventricular neuroepithelium or stem cell niche. Both HES1 and HES5 expression oscillate in NPC culture(Hirata et al., 2002; Imayoshi et al., 2013; Sueda et al., 2019). Most of our knowledge on dynamic HES/Her regulation in NSCs derives from *ex vivo* studies, whereas the differential contributions of *HES/Her* family members in NSCs and NPCs *in vivo* are less well understood.

Zebrafish are an excellent model to dissect relative contributions of Notch-dependent and - independent HES/Her activities. Zebrafish have two HES1 homologs, *her6* and *her9*, which are expressed in a Notch-independent manner, and several HES5 homologs (*her2, her4, her12* and *her15*) that are strictly Notch-dependent(Chapouton et al., 2011; Sigloch et al., 2023). Genetic evidence revealed that activity of *her6* and to a lesser degree *her9* is required for NSC maintenance in larvae(Sigloch et al., 2023). In contrast, Notch-dependent *her* genes appear to predominantly control NPC lineage progression, and combined genetic loss of their activity hardly affects NSC number(Sigloch et al., 2023). This correlates to the dynamic changes of *her* gene expression with neural lineage progression, as judged from scRNAseq-derived pseudotime profiles, which show that *her6* is predominantly expressed in NSCs, while most Notch-dependent *her* genes are expressed in NPCs, with the exception of *her4* that is expressed in both NSCs and NPCs(Rothenpieler et al., 2026). Her6 expression has been shown to oscillates in hindbrain(Soto et al., 2020) and telencephalon(Doostdar et al., 2024), with the observed noise hindering a more detailed characterization.

Neural proliferation is broad in the zebrafish embryonic neural tube, but during early larval development distinct neural proliferation zones (NPZs) form throughout the CNS, which harbor long-term NSCs and drive brain growth(Wullimann and Knipp, 2000; Tallafuss, 2003; Zupanc et al., 2005; Grandel et al., 2006; Ryu and Driever, 2006; Ganz and Brand, 2016). With regard to NSC regulation, much effort has focused on the mostly quiescent zebrafish adult brain(Adolf et al., 2006; Chapouton et al., 2007; Foley et al., 2024). In contrast, establishment and regulation of proliferation in NPZs are less understood, in particular given the differences in NSC proliferative activity compared to the general ventricular layer. For mammalian systems, a model has been put forward distinguishing boundary regions of low proliferation with sustained high HES expression levels, as compared to compartment regions with lower oscillating HES expression(Baek et al., 2006; Kageyama et al., 2008). The zona limitans intrathalamica (ZLI) is one of these boundaries, next to the isthmic organizer and the floor and roof plates, which together share high sustained *Hes* gene expression. However, differences in HES oscillation and function in early NPZs like the thalamic proliferation zone (TPZ), as well as similarities to zebrafish remain open questions. Distinct Notch signaling modes may differentially contribute to NSC maintenance, and potentially also to spatial patterning and size control of neural proliferation zones. For example, classical Notch lateral inhibition predominantly controls NSC and NPC lineage progression, whereas Notch lateral induction may also control NSC pool size and spatial organization of NSC domains(Boareto et al., 2015; Bocci et al., 2020; Ortica et al., 2026). While the mechanisms regulating Notch lateral inhibition versus induction are unknown, recent work suggests that Her6 may shift Notch signaling towards lateral induction(Rothenpieler et al., 2026), which would support a role of Her6 in maintaining coherent NPZs(Sigloch et al., 2023). Her6 thus appears to play a special role in NSC maintenance and NPZ spatial patterning.

Here, we use high resolution *in vivo* imaging of Her6-mNeonGreen expression from the endogenous *her6* locus to determine dynamic parameters of Her6 oscillations. We find Her6 expression to oscillate at distinct frequencies in different NSCs, independent of anatomical context. Oscillations are unaffected by inhibition of Notch signaling, and largely cell-autonomous. A mathematical model identifies critical parameters for robustness of the Her6 oscillator.

## Results

### *Her6* mRNA expression in ventricular NSCs

We first characterized *her6* expression in distinct contexts of general ventricular neuroepithelium, regional organizing centers, and NPZs. Sox2 immunoreactive nuclei reveal NSCs to line essentially the whole ventricular wall of the larval brain from 2 days post fertilization (dpf) on, where BrdU labelling indicates actively cycling NSCs (Figure 1A). We focused our analysis of *her6* expression on the diencephalic TPZ(Scholpp et al., 2009; Sigloch et al., 2023), including the ZLI organizing center and thalamic NPZ. Whole mount *in situ* detection of *her6* transcripts (Figures S1 and S2) revealed that *her6* in the greater thalamus is expressed at two distinct levels. High *her6* levels prevail in the ZLI, where *her6* is coexpressed with *shha,* and in a second transversal stripe in the posterior domain of the rostral thalamus (rTh(p); Figure S1). Low level expression of *her6* was detected in the anterior domain of the rTh and the caudal thalamus (cTh), where high *her4* expression prevails (Figure S1B). Thus, expression of *her6* was detected throughout the ventricular layer (Figures S1C-F), albeit in some regions expressing cells appear scattered and the HCR signal difficult to distinguish from background. Based on expression patterns of proneural genes, neurogenic lineage progression in the thalamus proceeds from the ventricular Sox2 and *her6* positive NSC layer into the deeper neuroepithelium, were we detected *neurog1* expression in NPCs in subventricular layers (Figure S2D), and in even deeper layers *neurod1* in early neurons (Figure S2E; see also scheme in Figure 1H and I).

**Figure 1.**
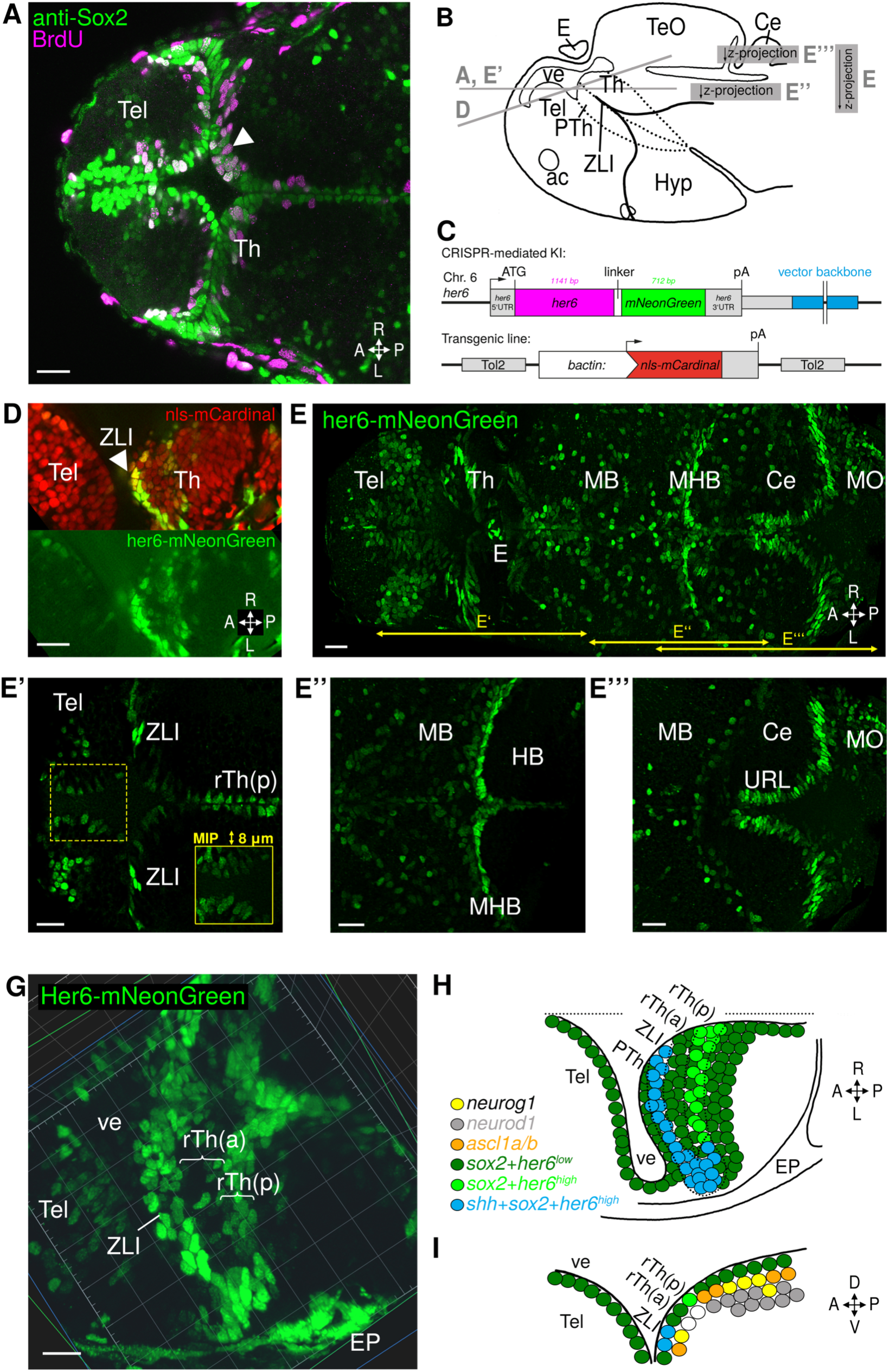
Her6 expression in NSCs of the ventricular wall and larval neural proliferation zones. **(A)** Anti-Sox2 immunostaining of NSCs and BrdU-labelling of cells in S-phase in the forebrain of a 56-hour old zebrafish larva. Arrowhead indicates BrdU-positive proliferating NSCs at the ventricle. **(B)** Anatomical model with section planes depicted in (A), (D) and (E). **(C)** Schematic of transgenes for in vivo labelling: Top: Knock-in of linker-mNeonGreen as carboxyterminal Her6-fusion into the endogenous *her6*-locus. Vector backbone sequences (blue) integrated 700 bp downstream of the 3’UTR and polyA site. Bottom: *beta-actin* promoter driven NLS-mCardinal transgene is expressed in all cell nuclei. **(D)** SPIM image of the forebrain showing Her6-mNG (green, *Tg(her6:her6-mNeonGreen)^m1366/m1366^*) combined with beta-actin:NLS-mCardinal (red, *Tg(beta-actin:NLS-mCardinal)^m1509^*(embryo07, time point 10, age 59.35 hpf). Lower panel: isolated Her6-mNG signal subjected to non-linear level adjustment (gamma = 2) to visualize low Her6-mNG expression in ventricular wall NSCs outside the ZLI NPZ. **(E)** Maximum intensity projection (MIP) of confocal stack of life 3 dpf *Tg(her6:her6-mNeonGreen)^m1366^*larva, position indicated in (B). (E’) Forebrain single confocal plane. Inset: 8 µm MIP of region marked by dotted line. (E’’) MIPs at the MHB and (E’’’) the cerebellar URL. **(G)** 3D reconstruction of Her6-mNG expression in the forebrain, centered on the ZLI (SPIM image stack, dorsal view, embryo07 at 59.35 hpf). **(H, I)** Schematic representation of the thalamic proliferation zones, (H) dorsal view, and (I) parasagittal section, orientation as indicated. Cells are color-coded according to expression as indicated. Sox2-positive NSCs green or blue. In subventricular layers, orange and yellow indicate progenitors, grey early neurons. White circles: cells expressing none of the listed neurogenesis markers. Abbreviations: A, anterior; ac, anterior commissure; Ce, cerebellum; D, dorsal; E, Epiphysis; EP, epidermis; HB, hindbrain; Hyp, hypothalamus; L, left; MB, midbrain; MHB, mid-hindbrain boundary; MO, medulla oblongata; Th, thalamus; P, posterior; PTh, prethalamus; R, right; rTh(a), rostral thalamus anterior domain; rTh(p), rostral thalamus posterior domain; Tel, telencephalon; TeO, optic tectum; URL, upper rhombic lip; V, ventral; ve, ventricle; ZLI, zona limitans intrathalamica. Scale bars: 20 μm.

### Distinct Her6 protein expression modes distinguish general ventricular NSC layer from neural organizers and proliferation zones

We developed an antibody against Her6 and demonstrated coexpression of Her6 and Sox2 in neural stem cells (Figure S3). Anti-Her6 immunofluorescence levels were highly variable in Sox2-positive NSC nuclei, and there were also Sox2-positive cells at the ventricular surface with very low or absent Her6 immunoreactivity (Figure S3B,C). Her6 appeared downregulated in cells leaving the ventricular surface, with signal in second tier cells typically lower, and very rarely detectable in deeper layers.

For documentation of Her6 dynamics *in vivo*, we engineered a fluorescent reporter strain by recombining the rapidly maturing mNeonGreen fluorescent protein coding sequence into the *her6* locus, generating a full-length Her6 fusion protein with a carboxy-terminal 11 amino acid linker and mNeonGreen [*Tg(her6:her6-mNeonGreen)^m1366^*, in short *her6^m1366Tg^*]. The *her6^m1366Tg^* encoded mRNA also contains the 312 bp wildtype *her6* 3′UTR (Figures 1C and S4A-D). Expression of *her6* and of *mNeonGreen* mRNAs in the TPZ of *her6^m1366Tg^*embryos were indistinguishable (Figure S4E). To validate Her6 activity of the Her6-mNeonGreen (Her6-mNG) protein, we generated *her6^m1358^; her9^m1368^* double mutant zebrafish larvae that are deficient in *Hes1* ortholog activity and develop a severe NSC deficient neural phenotype(Sigloch et al., 2023). A single allele of *her6^m1366Tg^*, similar to a single *her6* wildtype (WT) allele, completely rescued the *her6^m1358^; her9^m1368^*double-mutant phenotype, demonstrating full Her6 activity of Her6-mNG (Figure S4F). To enable better anatomical orientation, and to be able to track individual nuclei over time, we generated a *Tg(beta-actin:NLS-mCardinal ^m1509^* transgenic line that labels nuclei with a far-red fluorophore (Figure 1C,D).

We used light sheet volume imaging (SPIM) to determine Her6 expression in the TPZ *in vivo* (Figure 1D,G), and confocal volume imaging for reconstruction of the whole brain (Figure 1E-E’’’). As expected for the transcription factor Her6, the Her6-mNG signal was located to the cell nucleus. In the TPZ, Her6-mNG fluorescence was detected in bright transversal stripes correlating with ZLI and rTh(p), but at lower levels also in interspersed nuclei along the diencephalic ventricular walls (Figure 1D, lower panel). Reconstruction of the whole brain revealed that Her6-mNG is expressed at high levels in most NPZ and organizer regions of the early larvae, including the mid-hindbrain boundary and the upper rhombic lip (Figure 1E-E’’’). In addition, Her6-mNG was detected in a salt-and-pepper pattern throughout the ventricular NSC layer of telencephalon, diencephalon, mesencephalon, cerebellum and rhombencephalon. This indicates that endogenous Her6, similar to HES1, has multiple different expression modes: high-level variable expression in NPZs and organizer regions, and low-level variable expression in most other ventricular NSCs.

### Dynamic expression of Her6 *in vivo*

We performed time series *in vivo* SPIM volume imaging of homozygous *Tg(her6:her6-mNeonGreen)^m1366^* brains, typically starting at 56 hours post fertilization (hpf), when the TPZ has formed. Documentation was possible for 10-15 h, after which viability of the agarose-embedded larvae decreased. To track individual nuclei even at low levels of Her6-mNG expression, we crossed *Tg(beta-actin:NLS-mCardinal)^m1509^*into *her6^m1366Tg^*, and imaged double transgenic embryos at 9 min intervals (Figures 2, 3A and S5A-C; Methods). From the reconstructed volume time series data, we determined Her6-mNG fluorescence dynamics for individual cell nuclei. We exploited the high resolution of the primary image data to develop a measurement strategy aimed at reducing detection errors and noise. In brief, we placed three separate measurement spheres into each nucleus, each sphere consisting of 228 voxels, and determined average intensity values for each nucleus from the three spheres (Figure 2A first panel, Figure S5E). We performed quality control for each of the three measurement spheres for each nucleus and time point (see Methods), and corrected position of any misplaced sphere. This strategy resulted in measurement error far lower than the changes in Her6-mNG intensity over time (Figure S5D), and we think that therefore our time series data reliably reflect dynamics of Her6-mNG expression. Intensity profiles for Her6-mNG and NLS-mCardinal revealed cycling Her6-mNG fluorescence, while NLS-mCardinal fluorescence decayed over time (Figure 2C).

**Figure 2.**
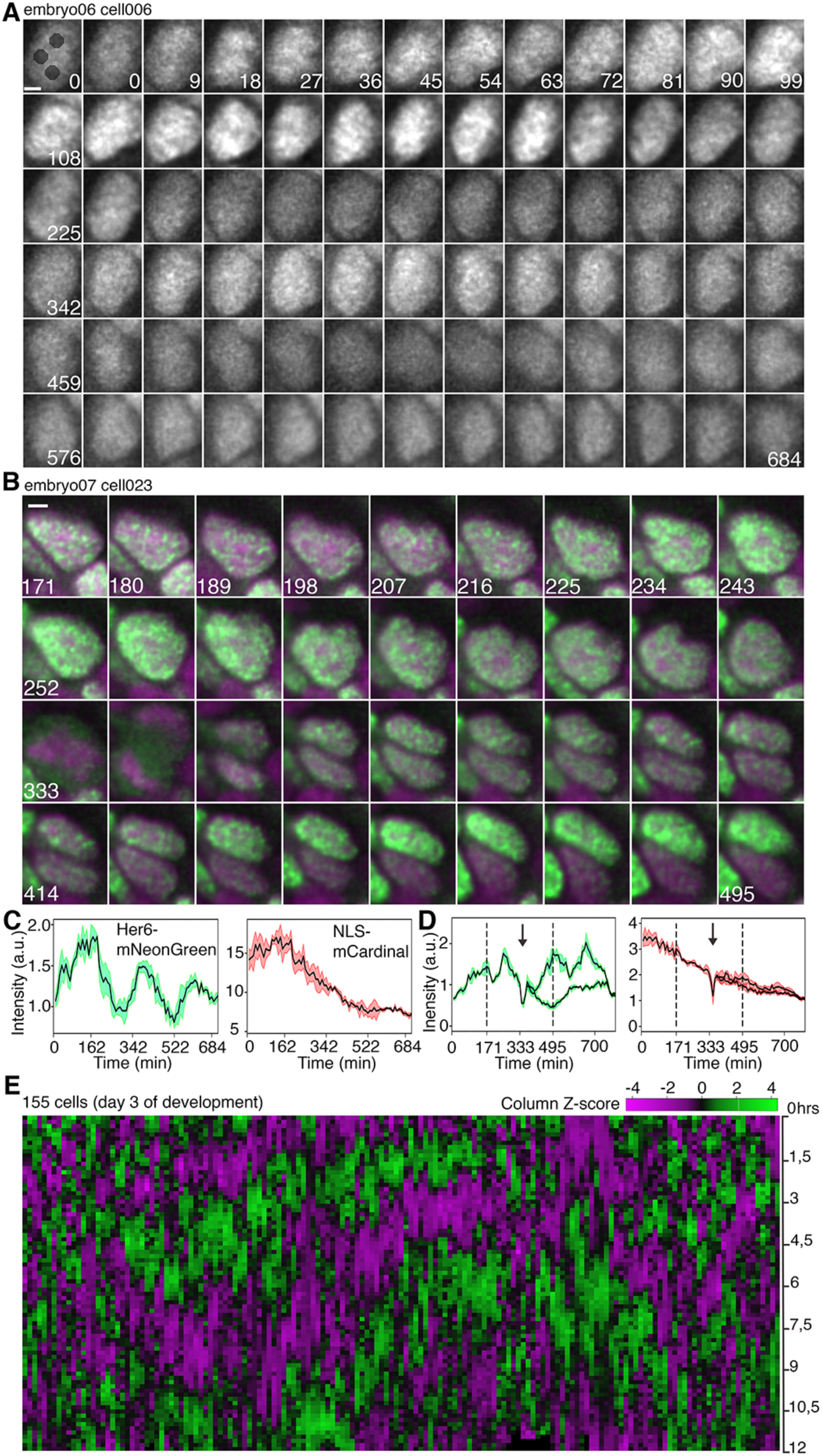
Dynamic expression of Her6-mNG *in vivo*. **(A)** *In vivo* SPIM imaging of a Her6-mNG–expressing cell in the ZLI starting at 56 hpf. Numbers indicate minutes after start of recording. Black spheres in the first panel indicate sphere positions for intensity measurements (for detail, see Figure S5E,F). Scale bar, 2 µm. **(B)** SPIM time series of a cell division in the ZLI starting at 60.85 hpf. Numbers indicate minutes after start of recording. Green, Her6-mNG expression; magenta, *beta-actin* driven NLS-mCardinal expression. Scale bar, 2 µm. **(C)** Quantification of the cell shown in **A**. Left, Her6-mNG mean intensity with standard deviation (SD) shaded. Right, NLS-mCardinal mean intensity, SD shaded. **(D)** Quantification during cell division shown in **B**. Same color code as in **C**. Dashed lines mark beginning and end of time series shown in **B**. Arrow indicates cell division. **(E)** Heatmap showing hierarchical clustering of detrended Her6-mNG intensity data of 155 tracked cells during day 3 of development with start of tracking set to zero irrespective of exact developmental time. For non-clustered data on developmental time scale see Supplementary Figure S5G. Each column represents one cell with data scaled by Z-score.

We tracked and measured a total of 356 cell nuclei from the dorsal diencephalon and telencephalon on the third day of development (46 to 71 hpf; 5 independent larvae), and selected 244 cells with long tracks (average 13 h) for further analysis (Figure S5G; Methods). All nuclei showed repeated cycles of increasing and decreasing Her6-mNG fluorescence (Figures 2A,E and S5B). ZLI nuclei remained above background also in the low expression phase, whereas telencephalic ventricular layer NSCs were difficult to distinguish from background during their low expression phase. We did not observe any nuclei with stable constant level Her6-mNG expression.

The data also enabled us to investigate Her6 expression after cell division in daughter cells. Among 14 cell divisions documented, in 9 Her6-mNG expression remained at similar levels in both daughter cells, whereas in 5 divisions one daughter cell started a new cycle of high Her6-mNG expression, while the other cell continued at lower levels (Figure 2B,D). Thus, after mitosis, daughter cells may independently initiate new cyclic expression of Her6-mNG. We further analyzed nuclei with overall decreasing Her6-mNG fluorescence, and observed in some nuclei that stayed at the ventricular surface prolonged decrease through several oscillation phases (Figure S6A,A’), while other nuclei with decreasing Her6-mNG signal moved away from the ventricular surface (Figure S6B,B’) into subventricular layers in which *sox2-*negative and *neurog1*-positive progenitors reside. We conclude that downregulation of Her6 may correlate with exit from the ventricular NSC layer into the subventricular NPC zone.

### Mathematical analyses of Her6 oscillations

To enable analysis of potential oscillations, long-term trends were removed(Phillips et al., 2017) (Figure S7A-C). For 155 cells with tracks longer than 12 h, we aligned the first 80 time points, applied hierarchical clustering to the time series Her6-mNG data, and generated a heatmap (Figure 2E). The heatmap reveals that all tracked nuclei have dynamic Her6-mNG expression, a majority cycling with apparent 3–5 h periods.

Next, we analyzed Her6-mNG expression in stringent mathematical terms (see Methods). To identify cells with low variability in frequencies of potential oscillations (stationary processes), the instantaneous phase was calculated from the analytical signal of the detrended time series (Figure S7A,B,D). The instantaneous phase presents the unique time-varying phase angle of a time series signal at any exact moment, and facilitates analysis of oscillations with frequency or amplitude changing over time (Figures S7D and S8A). Then, the method of surrogate data was applied to the instantaneous phase to decide whether the time series of a specific cell originates from a stationary or a non-stationary process (Figure S7A)(Hauber et al., 2022). To investigate whether oscillations were present in cells with stationary time series, we applied two independent tests. First, the power spectrum was estimated, and analyzed for statistically significant peaks (spectral analysis). Second, a model selection approach was applied by comparing the agreement between the data and two Gaussian Process (GP) models: a first-order stochastic process that by construction is not able to exhibit oscillations, versus a second-order process that can be oscillatory(Phillips et al., 2017). A likelihood ratio statistical test was applied to discriminate between the two GP models. Frequencies extracted from the power spectrum and from the GP models were similar (Figure S7E).

These techniques grouped the cells into four classes (Figure S7A and S8E,F). Class 1 (10% of 244 cells analyzed): cells with non-stationary time series typically express Her6-mNG dynamically, but dynamic properties change over time (includes short time series, single peaks, continuous decay of signal). Class 2 (31%): stationary but non-oscillating cells for which both methods (spectral analysis and GP regression) agree on non-oscillating behavior, i.e. the absence of a second-order oscillating process. These cells show repeated increase and decrease in Her6-mNG expression, but no constant period length (Figure 2E and Supplementary Data Sheet 1). Among the nuclei that did not oscillate (Figure S8F right half), some showed stochastic fluctuations (example: cell e07c045), while others appeared to document the decay phase of Her6-mNG expression that may coincide with exit from the Sox2-positive ventricular layer (example: cell e07c006; Figure S6B, B’). Class 3 (59%): stationary potentially oscillating cells, which include cells for which only one of the two methods states the presence of oscillations (“indeterminate” in that Her6-mNG oscillates, but not regularly enough for both methods to define a period). Class 4 (28%): oscillating cells for which both methods determine oscillatory behavior with an average period of 3.5 h (SD 1.9 h; Figures 3A, S8E). The oscillation period distribution did not show a sharp peak, but was rather broad with periods ranging between 2 and 7 h (Figures 3A, S8A). In summary, not a single cell expressed Her6-mNG at constant levels, but all were dynamic, with mathematical methods identifying oscillation periods for nearly two thirds of all cells.

**Figure 3.**
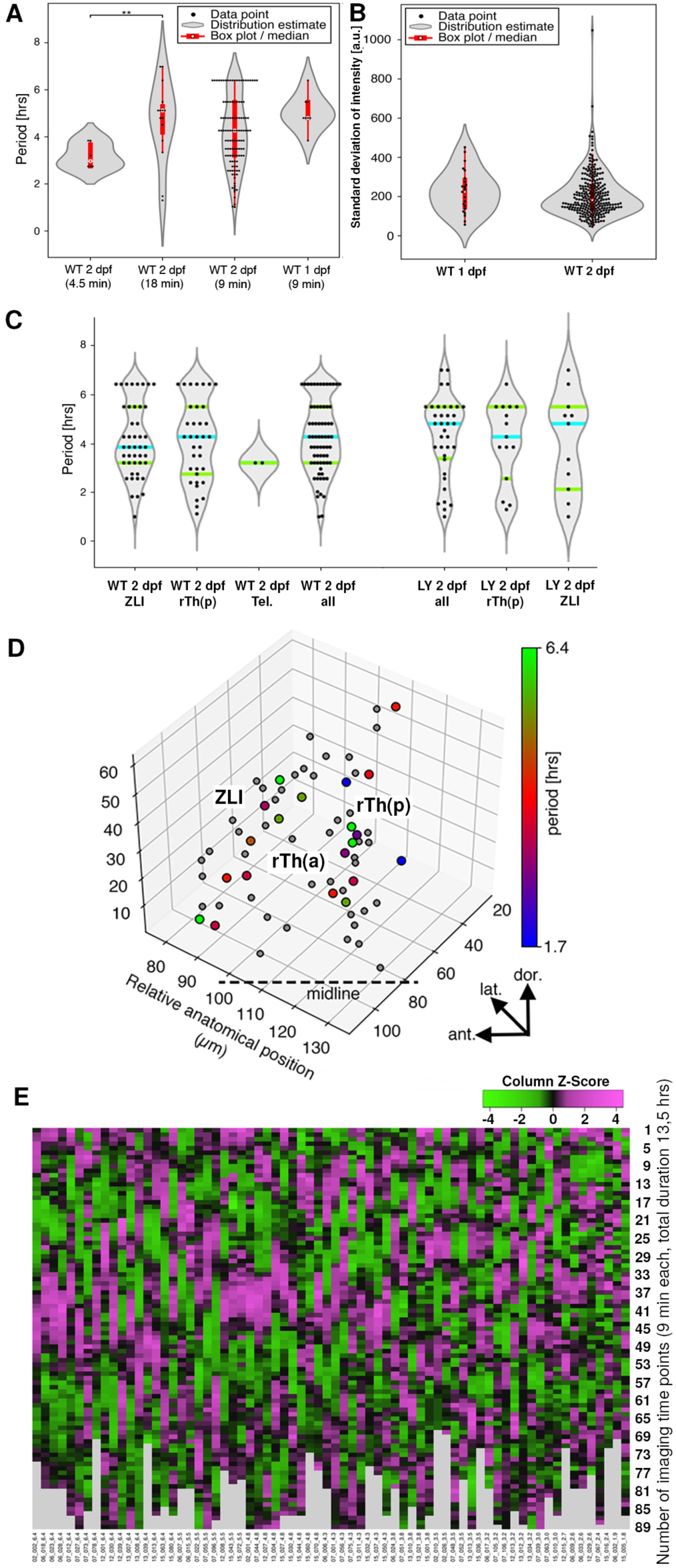
Characterization of Her6-mNG oscillations. **(A)** Estimated distributions of significant periods found in the spectra of individual cells, grouped by sampling interval times and age (** p=0.02). **(B)** Estimated distribution of standard deviation of fluorescence intensity tracks as proxy for amplitudes for individual cells during second (start 1 dpf) and third (2 dpf) day of development. **(C)** Estimated distributions of significant periods found in the spectra of individual cells, grouped by anatomical region for wildtype and LY treated Notch signaling inhibited 2 dpf embryos. **(D)** Position of cells in a 3D anatomical reference volume showing ZLI and thalamus of the right half of the brain. Cells are color-coded for Her6-mNG oscillation period. **(E)** Heatmap showing detrended Her6-mNG intensity data for all 2-3 dpf wildtype cells classified as oscillatory and sorted from left to right by decreasing period length. Cell numbers and oscillation periods are indicated below each column (NN_NNN_N.N gives embryo number – cell number – period length in h). Oscillation periods are based on major frequencies from the spectral analysis of oscillatory cells. The start of tracking was set to zero for each cell irrespective of exact developmental time. Data were scaled for each cell (column) and Z-scores are plotted using a color scheme magenta – high to green – low. Grey indicates time points after end of recording.

Given that oscillation frequencies vary significantly from cell to cell, we asked whether our mathematical classification defined cell populations with distinct biological features. We first compared mean Her6-mNG expression levels (fluorescence intensities; Figure S8B), and, as proxy for amplitudes, the standard deviation of the intensity (Figure S8C) for each cell, and found no differences for the four mathematical classes. Second, using data from one embryo (e07), we generated heat maps for cells separately for each of the four mathematical classes defined above (Figure S8F). It appears that, irrespective of the mathematical class, all cells had highly dynamic apparently cycling Her6-mNG expression profiles. Third, we asked whether cells of different mathematical classes cluster anatomically or move differently during the recording window. Following cell tracks over time, the positional changes were mostly caused by global anatomical movements, including progressing cephalic flexure. However, all cells followed parallel tracks independent of the mathematical class, indicating similar cell behavior. We also observed no anatomical clustering of cells by the mathematical class. We interpret these findings in that Her6 expression periods and likely also amplitudes are variable from cell to cell, but may not define cell populations with distinct biological features.

### Characteristics of Her6 oscillations during development

We evaluated whether Her6-mNG oscillation parameters change with maturation of NPZs during the second and third day of development (Figures 3A, S8A, E). The average periods in oscillating cells were longer in 1 dpf embryos (1 dpf 4.96 h vs. 2 dpf 3.46 h; p=0.06), indicating that the Her6 oscillator may be developmentally modulated. However, instantaneous frequencies of all cells plotted over time were largely constant from 24-71 hpf (Figure S8A), arguing against a prominent developmental modulation. We further compared mean intensities of Her6-mNG fluorescence in the 1 dpf and 2 dpf time series, and found a wide range of intensities in each time window, but no significant differences (Figure S8D).

We determined whether oscillation periods differ between anatomical regions of diencephalon and telencephalon (Figure 3C), and mapped color-coded oscillation periods for each oscillating cell into a 3D anatomical model (Figure 3D). Oscillation periods did not cluster anatomically. Oscillation periods of cells assigned to ZLI, rTh(p) or telencephalon did not significantly differ (p>0.7; Figure S8E). Thus, anatomical location is not a determinant of Her6 oscillation periods. Recent work reported that Her6 oscillations in the zebrafish hindbrain have a median period of approximately 1.3 - 1.5 h when 28-34 hpf embryos were examined(Soto et al., 2020). Because our light sheet imaging was focused on the forebrain area, we did not examine oscillations in the hindbrain. There may be differences between fore- and hindbrain, or our long observation periods, as compared to shorter detection windows, are better for detecting longer oscillation periods when the noise is high(Soto et al., 2020).

However, when we compared relative intensities of Her6-mNG fluorescence in data normalized to the level of NLS-mCardinal nuclear fluorescence, we observed a significant difference between expression levels in NPZ and organizer regions versus general ventricular NSC layer of the telencephalon. While comparison of ZLI organizer and rTh(p) NPZ did not detect differences (p=0,73), relative fluorescence intensity in both domains is about 4 times higher than in the general ventricular NSC layer of the telencephalon (p=1.0 E-09 and p=3.4 E-07). Therefore, in zebrafish organizer and NPZ Her6-mNG expression appears significantly higher than in general ventricular NSCs, consistent with our volume whole brain imaging (Figure 1E), and similar to data reported for Hes1 in boundaries and compartments for mammals(Baek et al., 2006).

### Notch-independent cell-autonomous Her6 oscillation

Based on cross regulation of *her* genes in zebrafish(Sigloch et al., 2023), Notch signaling may also influence oscillation parameters of Notch-independent Her6. We therefore tested whether *her6* expression dynamics may be affected through Notch signaling, either directly, or indirectly through the Notch target *her4.* The Notch inhibitor LY-411575 strongly suppressed *her4* expression in treated embryos (Figure 4A; Figure S9A,C,F,G), but had little effect on *her6* expression (Figure 4A; Figure S9B,D), confirming that *her6* is Notch-independent(Hans et al., 2004; Sigloch et al., 2023). When we analyzed Her6-mNG temporal dynamics upon LY-411575 treatment (Figure 4B), we found that Her6-mNG was oscillating in 40% of cells in treated embryos showed (Figure S8E), and that oscillation frequencies, mean intensities, instantaneous frequencies, and amplitudes did not differ significantly from untreated age-matched embryos (Figure 4D,E, Figure S8A, S10A-D). Therefore, Her6-mNG oscillations are not affected by inhibition of Notch signaling under conditions where expression of Notch-dependent *her4* is inhibited.

**Figure 4.**
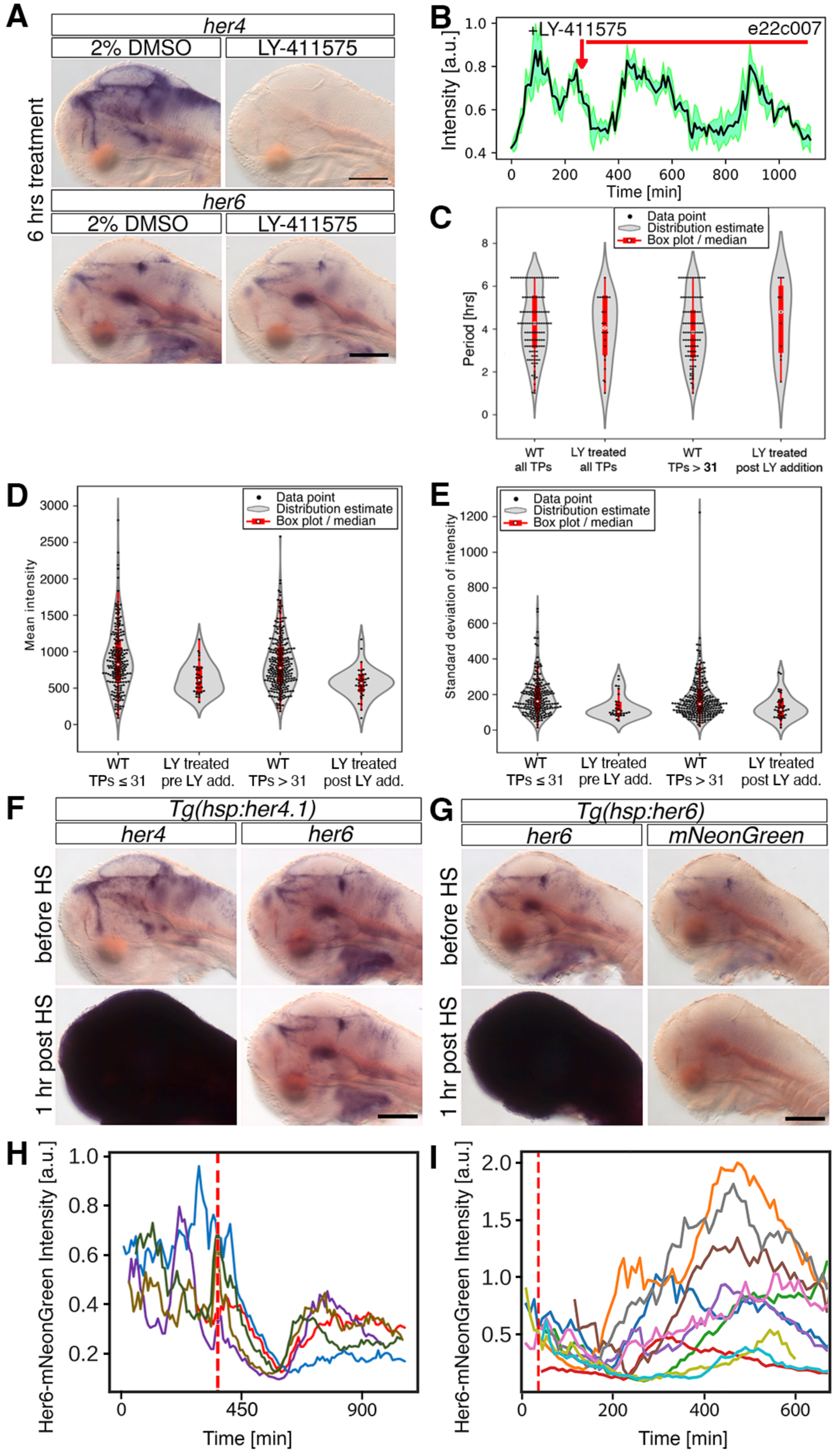
Her6 oscillation is robust and Notch-independent. **(A-E)** Notch inhibition. **(A)**, *In situ* hybridization for *her4* (top) and *her6* (bottom) in LY-411575 (50 µM) treated and control embryos (2% DMSO). Scale bars 200 µm. **(B)** Her6-mNG fluorescent single nucleus track, LY inhibitor addition as indicated. **(C)** Period length in cells of embryos treated with LY (LY addition time point em22 after imaging cycle 28, em23 after cycle 31); left: all time points (TP) show full dataset (WT control from Figure 3C); right: reduced dataset TPs only after LY addition, for WT control time points >31. **(D)** Comparison of mean fluorescence intensities of individual cells, groups as in (C). **(E)** Comparison of standard deviation of fluorescence intensity tracks of individual cells as proxy for amplitudes, groups as in (C). **(F-I)** Heat-shock overexpression of *her4* or *her6* **(F)** Heat-shock overexpression of *her4*. In situ hybridization of *her4* (left) and *her6* (right). Scale bars 200 µm. **(G)** Heat-shock overexpression of *her6*. In situ hybridization of *her6* (left) and *mNeonGreen* (right) in *Tg(her6:her6-mNeonGreen)^m1366^* heterozygous embryos. Scale bar 200 µm. (**H,I)** Her6-mNG fluorescent single nucleus tracks, Her6 heat-shock indicated by red dotted line. Continuous imaging of live zebrafish is limited to 12-15 h, and thus we focused on the time before and after heat-shock in **H** (5 cells tracked, color coded). In a second experiment, we characterized oscillations for 10 h after heat-shock (**I**, 9 cells).

Next, we tested whether overexpression of Her4.1 affects *her6* expression. We used the heat-shock–inducible *Tg(hsp70l*:*her4.1-FLAG) ^m1541^* transgenic line(Sigloch et al., 2023), and analyzed *her4* and *her6* expression after heat-shock at 15-min intervals for 4 h (Figures 4F, S11). *her4* expression was increased for more than 3 h after heat-shock, but *her6* expression was not significantly reduced in heat-shocked *Tg(hsp70l*:*her4.1) ^m1541^* embryos (Figures 4F, S11D,E). This demonstrates that overexpression of Notch-dependent Her4.1 does not affect *her6* expression.

A strong coupling of cells would cause local synchronization of changes in fluorescence intensity of Her6-mNG expressing cells. However, this was not observed when plotting intensity over time in anatomical space. In addition, we applied mathematical tests to detect potential coupling of Her6 oscillations between stably oscillating cells (Figure S10E-H, Supplementary Methods). Coherence analysis showed the correct size of the test in the simulation study, and that a 20 % cell non-autonomous contribution to the oscillation would have been detected in the experimental data (Figure S10G,H). Since the number of false positives detected was not higher than expected under the null hypothesis of cell autonomy, we conclude that there is no coupling of Her6 oscillations between cells. Given that we not only compared anatomical neighbors, but rather all cells displaying Her6 oscillation, our interpretation is that systematic cell-environmental effects, which should affect several cells in a similar manner, are unlikely, and that Her6 oscillation in the forebrain is therefore largely cell-autonomous.

### Auto-regulation and robust reinitiation of Her6 oscillation

To investigate auto-regulation of *her6*, we used transgenic *Tg(hsp70l:her6-FLAG)^m1492^* to overexpress Her6 in *her6^m1366Tg^* heterozygous embryos. This enabled detection of effects on *her6-mNG* mRNA levels as surrogate for endogenous *her6* expression in presence of high levels of heat-shock derived *her6* mRNA (Figure 4G; Figure S11A,B). Heat-shock-induced Her6-FLAG completely eliminated *her6-mNG* expression for approximately 2 h after heat-shock, but *her6* and *her6-mNG* mRNA expression recovered rapidly from approximately 2.5 h after heat-shock onwards (Figure S11B,C). These data confirm *her6* negative autoregulation(Sigloch et al., 2023). We also performed heat-shock overexpression of *her6* within the SPIM microscope observation chamber to monitor Her6-mNG expression *in vivo* (Figure 4H,I). Her6-mNG fluorescence recovered robustly with about 1 h longer delay after end of heat-shock. While the intensity tracks suggest that Her6-mNG initiated simultaneously in measured cells, the synchronicity got lost soon. Furthermore, the rapid recovery of Her6-mNG expression after heat-shock perturbation suggests that *her6* expression is controlled by strong lineage dependent transcriptional activation mechanisms in NSCs.

### Modelling Her6 dynamics reveals critical kinetic parameters to sustain oscillation

Next, we aimed at developing a mathematical model to identify critical parameters of Her6 oscillations. We tested whether the observed Her6-mNG dynamics can be explained by a cell-autonomous Her6 feedback model. An ordinary differential equation (ODE) system was established to represent a *her6* mRNA production term, *her6* mRNA degradation rate, Her6 protein synthesis and degradation rates, and Her6 protein negative feedback on *her6* expression (Figure 5A; Supplementary Methods Equation 1). The linear chain trick(MacDonald, 1976) was used to accommodate potential delays in the feedback loop, such as those arising from nuclear export and translation of *her6* mRNA. The parameters of the model were estimated by the method of maximum likelihood(Raue et al., 2013). Identifiability analysis and model reduction were performed using the profile likelihood(Raue et al., 2009; Maiwald et al., 2016; Wieland et al., 2021) (Figure S12A-C; Supplementary Methods). This ODE system captured the temporal dynamics of Her6-mNG observed experimentally (Figure 5B) with reliable parameter estimation (Figure S12D). The model validates that Her6-mNG dynamics are indeed predominantly determined by cell-autonomous negative feedback auto-regulation.

**Fig 5:**
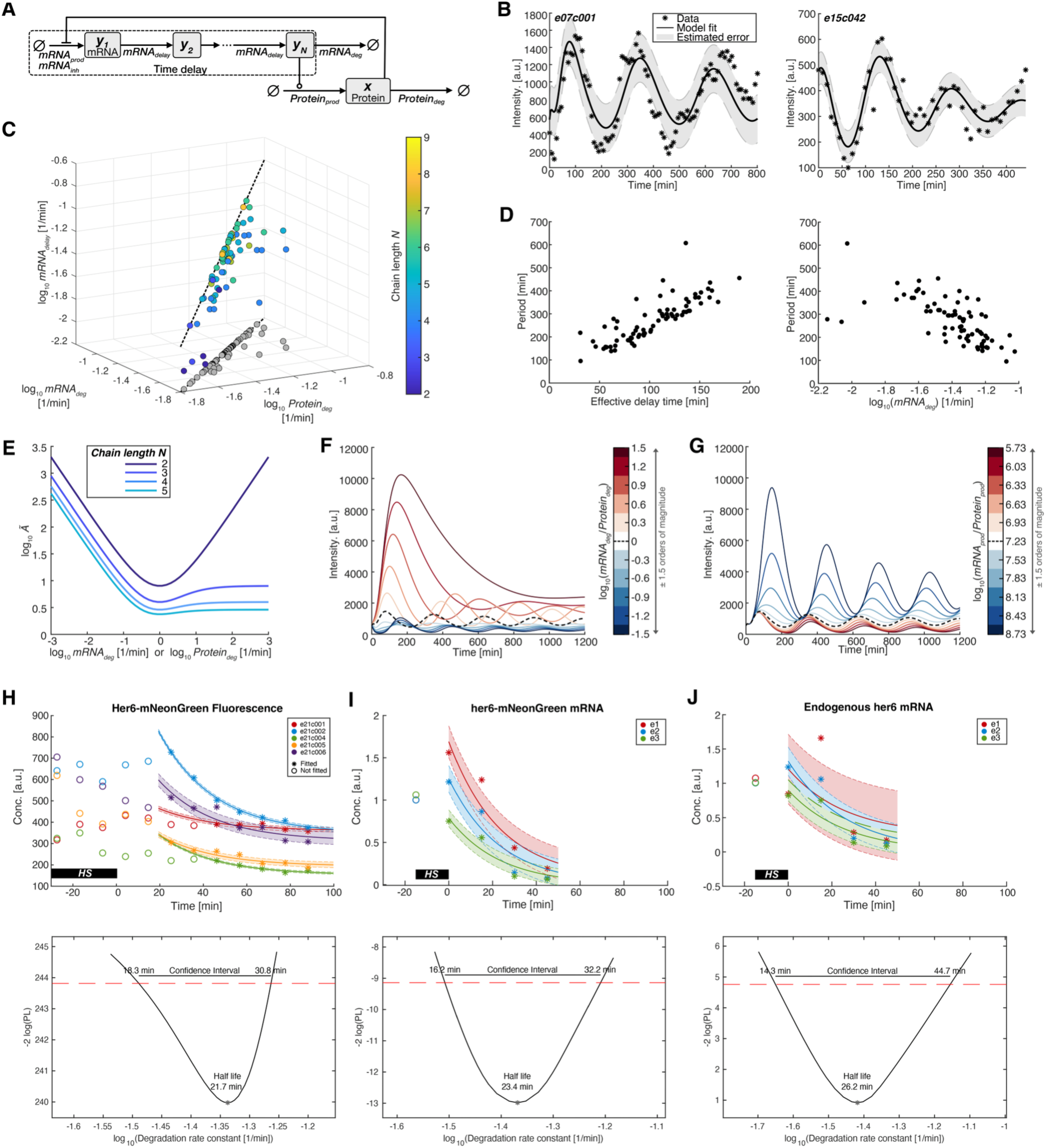
A dynamic model explains Her6 oscillations. **(A)** Pathway schematic for the mathematical model describing Her6 oscillations. Arrows, state transitions; circles, activation of a reaction; horizontal bars, inhibition of a reaction. **(B)** Data and fits of the model shown in (A) for two specific cells. **(C)** Representation of the fits for the individual cells in the space spanned by the relevant kinetic parameters. The chain length used for modelling is encoded in color. Grey dots represent ratios of *mRNA_deg_* and *Protein_deg_* only. The ratios of the parameters are close to one for all analyzed cells. The dashed black lines have unit slope and correspond to equal rate constants in the respective subspace. **(D)** Dependence of the oscillation period on the effective delay time and *mRNA_deg_*. Periods are estimated from the autospectrum of the model predictions of the protein time course for the estimated parameters of a specific time series. Each dot represents one oscillating cell. Left: Validates that effective delay time introduced by the model module highlighted by dashed box in (A) correlates with the period of oscillations exhibited by the model fits. Right: The period of oscillation is inversely correlated with the rate *mRNA_deg_*. **(E)** The critical value *Ã* (Methods 18.3) represents a measure for the restrictions in parameter space that hinder oscillations. It is minimized for *mRNA_deg_* = *mRNA_delay_* = *Protein_deg_*. Shown is the dependence of *Ã* on both *Protein_deg_* for fixed *mRNA_deg_* = *mRNA_delay_* = 1, and *mRNA_deg_* for fixed *Protein_deg_* = *mRNA_delay_* = 1. Colors indicate the different model chain lengths. The longer the chain, the less strict the condition for sustained oscillation becomes. **(F,G)** Dependence of the model simulations for the protein time course when varying (F) the ratio between mRNA and protein degradation rates, and (G) the ratios between mRNA and protein production rates. Simulations are performed with the best-fit parameter set of e07c001. The dashed black line shows the model prediction for (G) equal degradation rates or (F) optimal ratio of production rates. In both (F) and (G) the protein time courses were calculated for a range of 1.5 orders of magnitude for the ratios (color code). Variation in degradation ratios eliminates oscillations, while variation in production rates predominantly affects amplitudes. **(H-J) Top**: Data that approximately show (H) protein degradation (Her6-mNG fluorescence intensity data from 5 measured cells, em21), or (I,J) mRNA degradation (I) *Her6-mNG* mRNA and (J) endogenous *her6* mRNA, based on RT-PCR data from 4 embryos per data point, following *her6* heat-shock (HS) overexpression. Lines indicate the corresponding model fit and shaded areas the estimated error. Baselines, initial values, and noise levels were estimated individually for each dataset. **Bottom:** Profile likelihood of the parameters of the degradation model (Equation 2) protein (left panel) and for mRNA (right two panels) for the data shown in H-J top panels. The confidence interval of *mRNA_deg_* (best fit half-life 23.4 min for *her6-mNeoGreen* mRNA, and 26.3 min for endogenous *her6* mRNA) includes the best-fit value of *Protein_deg_* (21.7 min for Her6-mNeonGreen). Therefore, the result of the analysis of experimental data is in agreement with the outcome of equal degradation rates that resulted from mathematical modeling of oscillatory Her6 time series.

Given the observed variability in amplitude and frequency as compared to strictly periodic oscillations, we analyzed the parameters that contribute to Her6-mNG dynamics. As expected, the effective delay time in our model correlated with the oscillation periods observed for each oscillating cell, and the oscillation periods were inversely correlated with the mRNA degradation rate constant (Figure 5A,D). Surprisingly, model-based estimation of mRNA and protein degradation and mRNA delay rate constants from the actual time series data of all oscillating cells revealed that the ratios of these parameters are close to one for all analyzed cells (Figure 5C). A plot of model predictions for Her6 and *her6* half-lives for each cell confirms equal decay rate constants, while between different cells the half-lives vary, corresponding with the different oscillation frequencies observed (Figure S13). The critical value *Ã* (Methods 18.3) represents a measure for the restrictions in parameter space that hinder oscillations. It is minimized for *mRNA_deg_ = mRNA_delay_ = Protein_deg_* (Figure 5E, chain length =2)). The longer the delay chain, the less strict the condition on *mRNA_deg_* and *Protein_deg_* for sustained oscillation becomes (Figure 5E, chain length 3 to 5), which in biological terms suggests that if the *her6* mRNA would be subject, e.g., to a larger number of independent processing steps, each with distinct temporal requirements, the optimum of equal rate constants would get lost. Our model suggests that for Her6 it is mainly RNA and protein stability that are tuned to maintain oscillations, and that a coordinated combination of multiple independent mechanisms that act on their stability are essential to maintain oscillation. The strong effect of an imbalance of RNA and protein degradation rate constants in our model is illustrated by plotting expected Her6 dynamics at varying degradation rate constant ratios, strongly affecting both frequency and amplitude, with complete loss of oscillations at extreme ratios tested (Figure 5F). In contrast, when we tested the same range of variation of RNA and protein production rate constants, our model predicts only a minor effect on oscillation frequencies, while amplitudes are strongly modulated (Figure 5G).

To test the model predictions, we aimed at obtaining estimates of *her6* mRNA and Her6 protein degradation rate constants based on measurements in living larvae. *her6* mRNA measurements may be obtained from quantitative RT-PCR, however, this requires to artificially affect oscillations *in vivo* to eliminate the production and measure pure decay. We achieved this by exploiting the Her6 negative feedback loop: following *hsp70l:her6-FLAG* heat-shock, endogenous *her6* transcription is minimal or absent, and thus *her6* and *her6-mNG* mRNA (Figure S11C) and Her6 protein dynamics driven by degradation (Figure 4H). We were able to obtain the Her6-mNG decay data from em21 time series data, in which about 20 min after the heatshock strong Her6 negative feedback is observed (Figure S11C), and determined Her6-mNG degradation rate constants based on the mNG intensity profiles for the 20 to 100 min interval after heat-shock (Figure 5H). For mRNA rates, we used *Tg(her6:her6-mNeonGreen)^m1366^* heterozygous fish, which enabled us to simultaneously measure *her6-mNG* (Figure 5I) and endogenous *her6* mRNA(Figure 5J) decay rates from time series RT-qPCR analyses (Figure S14). The confidence interval of *mRNA_deg_* (best fit half-life 23.4 min for *her6-mNG* mRNA, and 26.3 min for endogenous *her6* mRNA) included the best-fit value of *Protein_deg_* (best fit value 21.7 min; Figure 5H-J). Therefore, the experimental data are in agreement with the conclusion of the mathematical model that Her6 oscillation is sustained by equal rate constants of mRNA and protein degradation.

### Model predictions on Her6 amplitude difference in NPZs versus ventricular NSC layer

Our estimates of relative Her6-mNG fluorescence in the ZLI and NPZ versus telencephalic ventricular NSC layer revealed about four-fold higher Her6 signal in these regions than in other ventricular NSCs. However, we did not observe a difference in mean oscillation periods. To understand potential differences in organizer and NPZ versus general ventricular NSCs, we investigated model predictions on amplitude in more detail. We note that our heat maps (Figures 2E, 3E, S5G, S8F) suppress information on the amplitude based on the use of z-scores for visualization. Further, due to progressive cephalic flexure during the second and third day of development, most recorded cells locate to deeper imaging planes as the time series progresses, which results in decreased fluorescence intensity (Figure 2C, Data Sheet S1).

However, there is also significant variation in instantaneous amplitudes observed (Supplemental Materials SM1 and standard deviation of intensity in Figure S8C). Given that a reduction in amplitude may be expected during cell state transitions to NPCs, and concomitant exit from oscillation, we investigated potential correlations of observed components of our model and the amplitude for individual cells. We used the standard deviation for each cell fluorescence intensity track as proxy for the amplitude and classified oscillating cells by period length (Figure S15A). We find a significant positive correlation between period and amplitude (p=0.0008; Figure S15B). Using our data of Notch signaling inhibited LY-treated embryos, we can also calculate a linear fit (albeit potentially due to fewer data points not significant: p=0.14; Figure S15C), suggesting that Notch signaling, next to not affecting period or coupling, may also not affect amplitude. We used our mathematical model to ask which components may have strong effects on the amplitude. While variation of *mRNA_delay_* has a major effect on period length, it has only a minor effect on amplitude (Figure S15D). In contrast, the strongest effects on amplitude are by variation of *mRNA_deg_* or *Protein_deg_* rate constants (Figure S15E,F, see also Figure 5F). However, varying degradation rate constants always comes at the cost of changes in oscillation frequency (Figure 5F). Therefore, our model suggests that differences in amplitude between NSCs in organizers and NPZs versus ventricular layer may predominantly be caused by differences in *her6* mRNA production (Figure 5G).

## Discussion

During zebrafish larval stages the brain grows tremendously, substantially fueled by proliferation of NSCs in NPZs like the thalamus and the upper rhombic lip, but also sustained by NSC proliferation in most other regions of the ventricular NSC layer(Wullimann and Knipp, 2000; Tallafuss, 2003; Ryu and Driever, 2006; Mueller and Wullimann, 2016). Across the NPZs, similar constraints may exist for the spatial organization of NSCs into coherent and stable zones of active NSCs. However, these NSCs at the same time must maintain a high potential for dynamic lineage progression into NPCs and early neurons. In zebrafish, the HES1 homolog Her6 plays a crucial role in NSC maintenance(Sigloch et al., 2023), but also in persistence of the ZLI organizer(Scholpp et al., 2009; Sigloch et al., 2023). Her6, but not the Notch signaling dependent Her factors, may also shift Notch signaling from a lateral inhibition to lateral induction mode(Rothenpieler et al., 2026), thereby potentially supporting larger coherent NSC zones. Here, we investigated the dynamics of Her6 oscillatory expression to begin to understand how Her6 encodes both NSC stability and plasticity.

We show that in TPZ and ZLI, Her6 expression oscillates between high peak and low values that in most cases are still above background. In contrast, in non-NPZ ventricular layer NSCs Her6 oscillates at overall lower levels of expression, and typically reaches background levels in the low phase. We did not observe any cells with persistent high non-oscillating Her6 expression. For TPZ and organizer regions, this is at variance with reports for Hes1, whose expression is reported to be maintained at persistent high levels in the mouse “boundary” region ZLI (Baek et al., 2006). In the mouse ZLI, high Hes1 expression has been linked to low levels of cell proliferation and NSC quiescence(Baek et al., 2006), while at 3 dpf in zebrafish, proliferation is still observed in the ZLI(Sigloch et al., 2023). We hypothesize that these observations may reflect three distinct Her6 dependent NSC states that differentially rely on the three main features of Her6: (1) the ability of high Her6 to promote quiescence by driving NSCs into cell cycle arrest(Rothenpieler et al., 2026) may dominate long-term NSCs in mature boundaries like the ZLI; (2) the ability of high Her6 to spatially pattern and enlarge NPZs by lateral induction(Rothenpieler et al., 2026) may drive establishment and maintenance of NPZs; and (3) lower Her6 oscillating levels may be sufficient to balance neural stem cell maintenance versus neurogenesis in non-NPZ ventricular NSCs. The dynamic equilibrium between such activities may relate to observations in mouse NSCs that Hes1 continues to oscillate in quiescent cells, although at elevated expression levels, and that HES1 oscillations persist when quiescence is induced in culture(Sueda et al., 2019; Marinopoulou et al., 2021).

For mice, Hes1 oscillation periods of 2.5 to 3 h have been reported, with slightly longer periods in NPC cultures derived at E9.5 compared to E14.5(Shimojo et al., 2008). Similarly, Hes5 oscillates with a mean period of 3.3 h(Manning et al., 2019). We observed a mean Her6 period of 3.5 h, with oscillations apparently sustained over a wider range of periods from 2 to 7 h. The similarities between Hes1 and its ortholog Her6 suggest conservation of mechanisms controlling oscillation parameters. Hes5 expression, predominantly controlled by Notch signaling, has been reported to oscillate in phase with Hes1 in murine NPC cultures(Imayoshi et al., 2013). However, we observed that *her4* mRNA is typically expressed at low levels in *her6* high-expressing cells, consistent with the strong repression of *her4* by Her6(Sigloch et al., 2023; Rothenpieler et al., 2026). Our finding that Her6 oscillations were not affected by inhibition of Notch argues against a prominent role of Notch signaling in Her6 oscillations, as neither period nor amplitude were affected. We note that these observations may only apply to *her6* in NSCs, but not necessarily to other tissues, as *her6* appears to be under different transcriptional control in the somite, where it is Notch-dependent(Pasini et al., 2004).

Several independent lines of evidence indicate that Her6 functions as a largely cell-autonomous oscillator. First, our coherence analysis indicates that Her6 oscillations are not coupled among cells by potential local, regional, or global cell-nonautonomous mechanisms. Thus, we could not observe cell coupling postulated previously from expression of destabilized Her6(Doostdar et al., 2024). We also did not detect any synchronized microclusters as reported for HES5 in the mouse ventral spinal cord(Biga et al., 2021). Second, Her6 mean expression levels as well as oscillation periods and amplitudes were unaffected by pharmacological inhibition of Notch signaling under conditions that strongly suppressed *her4* expression. Third, after mitosis, daughter cells could initiate distinct Her6 expression dynamics, consistent with independent reestablishment of the oscillator in individual cells. Lastly a delayed negative-autoregulatory model was sufficient to capture the measured Her6-mNG profiles in all oscillating cells, without need to imply communication between cells. We conclude that the Her6 oscillator is cell-autonomous, which together with its robustness against perturbations, and stable reinitiation of *her6* expression after experimental inhibition, are features that may contribute to the stability of the NSC lineage.

We developed a dynamic negative feedback model based on ODEs that faithfully described the observed Her6 oscillations. In this model, a *her6* delay chain (accounting for factors such as *her6* mRNA transit and translation), as well as production and stability of *her6* mRNA and Her6 protein, together determine the dynamic properties of the oscillator. Our analysis of model fits in parameter space revealed that oscillations are most prevalent when *mRNA_deg_ = mRNA_delay_ = Protein_deg_*. We validated this finding by demonstrating that *her6* mRNA and Her6 protein degradation rate constants are similar *in vivo*. Interestingly, theoretical models have previously suggested that self-repressed genes with similar mRNA and protein half-lives have the propensity to oscillate, and that the oscillatory behavior is optimized when the degradation rate constants are equal(Page and Perez-Carrasco, 2018).

An example for effects of variation of *mRNA_delay_* has been the somite oscillator, in which variation of *her* gene intron length controls the delay in the negative feedback loop and thus properties of the oscillator(Takashima et al., 2011; Harima et al., 2013). Intron length has also been manipulated for *Hes1* and shown to control oscillation parameters(Ochi et al., 2020). Distinct mechanisms may have evolved to calibrate oscillations through control of degradation rate constants. For example, miR-9 is a conserved regulator of HES mRNA stability(Li et al., 2006; Bonev et al., 2012), and has been shown to also regulate zebrafish *her6*(Soto et al., 2020). At the protein level, ubiquitination has been shown to regulate HES/HER protein stability throughout evolution(Barry et al., 2011; Kobayashi et al., 2015). Global control of protein stability in segmentation clock cells has been revealed to be responsible for the slower human compared to mouse clock oscillations(Matsuda et al., 2026), and the tempo of neural development(Nakanoh et al., 2026). So far unknown mechanisms may have evolved to stabilize the oscillator by ultimately equalizing mRNA and protein degradation rate constants.

It is intriguing to speculate that evolution may have avoided to make the oscillator excessively stable, as intrinsic sensitivity to changes in Her6 protein and mRNA stability also provides a possible pathway for exit from oscillatory states, and therefore for cellular plasticity. Still, it is difficult to see how the Her6 oscillator works reliably given the significant variation in frequencies, amplitudes, and the presence of noisy fluctuations. RNA and protein stability in general are affected by multiple signals and physiological parameters, and the constrains to keep these mechanisms responsive to other needs in NSCs may have resulted in evolution not prioritizing precision in oscillation, but responsiveness of mRNA and protein decay mechanisms to other signals. Indeed, several mechanisms have been demonstrated to link protein degradation and RNA stability, for example though the ubiquitin proteasome system(Cano et al., 2012; Thapa et al., 2020), by HSP90 and PMR60(Peng et al., 2008) in context of heat shock response(Laroia et al., 1999), and in cellular stress(Stohr et al., 2006) or cancer(Yu et al., 2026). Therefore, the variable periodicity may not represent a defect of the oscillator, but rather a consequence of its regulatory flexibility integrating multiple inputs, while negative feedback loops stabilize the dynamics against noise and perturbations(Kaern et al., 2005).

The mathematical model also predicts that changes in production terms hardly affect the oscillation periods, but effectively alter the amplitude. This may explain why we find higher Her6 expressing cells of the NPZ and ZLI to oscillate with very similar parameters as other ventricular NSCs that express at low levels. The differences in expression level may simply be explained by Shh in the ZLI providing an addition independent transcriptional activation input for *her6.* In mice, a role for Shh in activation of HES1 expression has been shown in several contexts, including cooperation of Shh and Notch signaling in control of HES1 in the neocortex (Dave et al., 2011), and in Shh activation of Notch-independent HES1 expression in retinal progenitors(Wall et al., 2009). We can only speculate that oscillation at different expression levels may encode distinct Her6 function: at high levels lateral induction may shape NPZs and cell cycle regulation quiescence in boundary regions(Rothenpieler et al., 2026), while at lower levels Her6 in ventricular neuroepithelium may maintain stem cell and control progression into NPCs.

Our mechanistic analysis of Her6 expression dynamics provides a framework to understand distinct modes of Her6 regulation in control of ventricular NSCs broadly, but also specifically in NPZs, by detailing distinct contributions of *her6* production and decay as well as Her6 decay to oscillator features that ultimately control stability as well as lineage dynamics in NPZs.

## Materials and Methods

### 01 Zebrafish maintenance

ABTL strain zebrafish were kept and bred at 28°C. Experiments were performed in accordance with the German Animal Welfare Guidelines. All experiments were approved by the ethics committee for animal experiments at the Regierungspräsidium Freiburg. Zebrafish embryos were incubated in E3 medium (5 mM NaCl, 0.17 mM KCl, 0.33 mM CaCl_2_, 0.33 mM MgSO_4,_ supplemented with 2 ppm Methylene blue). To prevent pigmentation, N-Phenylthiourea (PTU, Sigma Aldrich) was added to the E3 medium to a final concentration of 0.2 mM. The following published mutant lines were used: the CRISPR/Cas9 knockout alleles *her6^m1358^* and *her6^m1368^,* which each generate a frameshift that deletes most of the protein including the bHLH domain (see Figure 3 in (Sigloch et al., 2023)). The following published transgenic lines were used: *Tg(hsp70l:her4.1-FLAG)^m1541^*and *Tg(hsp70l:her6-FLAG)^m1492^* (Sigloch et al., 2023), see also Figure S11A).

### 02 Generation of transgenic lines

New transgenic lines were generated using the Tol2 kit vector system (Kwan et al., 2007). To label all cell nuclei with mCardinal we cloned *beta-actin:mCardinal-NLS* by fusing the NLS sequence (MAPKKKRKV from base pairs 5’-ATGGCTCCAAAGAAGAAGCGTAAGGTA-3’) to the C-terminus of mCardinal (Chu et al., 2014). A 5.3 kb beta-actin promoter (Chi-Bin Chien lab, gift from Ken Poss) was cloned upstream of mCardinal into a Tol2 vector containing Tol2 sites and a red heart marker (*cmlc2:tagRFP*), see Figure 1C lower construct. The Tol2 vector was sequenced and injected together with 40-60 µg/µl mRNA encoding Tol2 transposase into 1-cell stage zebrafish embryos. Transposase mRNA was transcribed using the SP6 mMessage mMachine kit (Thermo Fischer). The following alleles were used in this study: *Tg(beta-actin:mCardinal-NLS)^m1508^*and *^m1509^*.

### 03 Targeted generation of knockout and knock-in alleles

The CRISPR/Cas9 system was used for targeted manipulation of *her6*. Cas9 mRNA was transcribed from the pCS2+hspCas9 plasmid (Ansai and Kinoshita, 2014) using the SP6 mMessage mMachine kit (Thermo Fischer) and a polyA tail was added with the PolyA tailing kit (Thermo Fischer). For the *her6-mNeonGreen* knock-in, two sgRNAs each were designed with CRISPRscan (Moreno-Mateos et al., 2015) and ordered as oligonucleotides to generate a DNA template (Gagnon et al., 2014). Oligonucleotides were annealed cooling from 95 °C down to 25 °C in 1 hr. Annealed oligonucleotides were filled with T4 DNA Polymerase (New England Biolabs) for 20 min at 11 °C. sgRNAs were transcribed 3-4 h at 37°C using the MEGAscript T7 kit (Thermo Fischer). Sequences of specific oligonucleotides for the transcription of sgRNAs include: The T7 promoter (left) shown in lower case letters, the target specific region of the sgRNA in the middle in capital letters, and the annealing site for the constant oligonucleotide (right, lower case).

For the *Her6-mNeonGreen* knock-in, the sgRNAs transcribed from Oligo04 and Oligo23 were used combined. The Her6-mNeonGreen *her6:her6-linker-mNeogreen-3’UTR[her6]* knock-in donor plasmid pCS75c1 as shown in Figure S4B was assembled by Gibson cloning. The left and right homology arm were PCR amplified (Pfu Ultra II, Agilent) from DNA of ABTL embryos. The following primers were used to amplify the homology arms: p108F + p109R, p110F + p111R, p114F + p115R. p121F + p122R were used to amplify the pUC19 backbone. Primers p112F + p113R were used to amplify the *mNeonGreen* ORF (Allele Biotech) (Shaner et al., 2013) from a plasmid that contained an additional polyadenylation signal, which was removed in a subsequent cloning step. The resulting plasmid pCS61 contained the unwanted polyadenylation signal between the 3′UTR and the right homology arm. This polyA signal was removed by a PCR with p132F and p133R. The right homology arm was cloned again from genomic ABTL DNA to elongate it (p196R + p132F). This fragment was assembled by Gibson cloning with the fragment amplified by p195F and p133R from pCS61. To avoid cutting the template with the targeting sgRNA, the Oligo23 binding site was wobbled in the homology arm using p136F + p137R. pCS75c1 was confirmed by sequencing. To generate knock-in zebrafish, 1 nl of the following injection mix was injected into 1-cell stage embryos: 0.5 µl *her6-mNeonGreen* donor knock-in plasmid pCS75c1 (650 ng/µl), 3 µl Cas9 mRNA (approx. 600 to 1200 ng/µl), 1 µl sgRNA from Oligo04, 1 µl sgRNA from Oligo23. F1 knock-in embryos were identified by PCR and based on mNeonGreen expression in proliferation zones, and F2 lines established.

Genotyping of mutant alleles: DNA was isolated from whole live embryos or tail biopsies of fixed 96 hpf embryos in 60 µl TE (10 mM Tris and 1 mM EDTA) buffer by heat denaturation at 95 °C for 5 min and digestion by adding 3 µl Proteinase K (20 mg/ml, Sigma) for 4 h at 55 °C, followed by heat inactivation for 10 min at 95 °C. PCRs (annealing temperature 58 °C, 30 cycles) were performed using MyTaq (Bioline). For genotyping the *her6* knockout allele *m1358*, primers p5F + p10R were used. For genotyping the *her9* knockout allele *m1368*, p197F + p198R were used. For genotyping the *her6-mNeonGreen* knockin allele *m1366* primers p5F (genomic *her6,* not on plasmid) and p129R (on *mNeonGreen*) were used.

Sequencing: Genomic DNA for sequencing was amplified by Pfu Ultra II (Agilent) unless otherwise stated and sequenced at GATC (Eurofins Deutschland). For *Tg*(*her6:her6-mNeonGreen*)*m1366*, DNA from homozygous knock-in embryos was used and the locus was amplified with Takara LA Polymerase (Takara). For genotyping *Tg*(*her6:her6-mNeonGreen*)*m1366*, primers p5F (upstream of integration site) and p352 (downstream of integration site) were used.

### 04 Experimental manipulation of embryos

Heat-shock driven overexpression: Embryos heterozygous for the transgenes *Tg(hsp70l:her4.1-FLAG)m1541 or Tg(hsp70l:her6-FLAG)m1492* were used. For the analysis of mNeonGreen expression in *Tg(hsp70l:her6-FLAG)m1492*, embryos heterozygous for *Tg(her6:her6-mNeonGreen)m1366* were analyzed. For expression analysis by *in situ* hybridization, the heat-shock was started at 55.75 hpf by incubation in a water bath at 39.5 °C for 15 min. After heat-shock treatment, the embryos were incubated at 28 °C and fixed in 4% paraformaldehyde at the indicated times post heat-shock. Control embryos not carrying the *m1541* or *m1492* alleles were heat-shocked in parallel, and *in situ* hybridizations were performed in the same tube. Quantification of *her6* mRNA response in *Tg(hsp70l:her4.1-FLAG)m1541* or *her6-mNeonGreen* mRNA response in *Tg(hsp70l:her6-FLAG)m1492; Tg(her6:her6-mNeonGreen)m1366* (heterozygous) after heat-shock was performed in single image planes using FIJI (same procedure as described in Methods section 08).

For heat-shock during live imaging, the heating element of the Zeiss Lightsheet.Z1 microscope was set to 39 °C for 30 min (embryo21) or 15 min (embryo33) (in the imaging chamber of the SPIM, 39°C were reached 9 min after start of the heat-shock). Quantification of Her6-mNeonGreen protein response in *Tg(hsp70l:her4.1-FLAG)m1541* after heat-shock was performed using arivis Vision4D (see section on fluorescence measurements). The Her6-mNeonGreen intensities for 5 cells before and after heat-shock are shown in Figure 4H, and for a longer period after heat-shock for 9 cells in Figure 4I.

LY-411575 Notch inhibition: For bulk treatment of embryos, Notch signaling was blocked by the incubation of embryos in 50 µM LY-411575 (Sigma) in 2% DMSO (AppliChem). Control embryos were treated with 2% DMSO. The E3 based media contained 1 mM tricaine methanesulfonate (MS-222). During the treatment, embryos were placed in a drop of 0.6% low melting agarose (Sigma) to incubate them in similar conditions compared to embryos imaged with the SPIM. For Notch inhibition during SPIM imaging, embryos were recorded first under standard conditions to observe normal *her6* expression dynamics, and then 100 µl of a 10 mM stock solution LY-411575 (LY; dissolved in DMSO) were added drop-wise to the chamber containing 20 ml E3 medium with PTU and Tricaine (final concentration 50 µM). The medium was pipetted up and down with the syringe that is connected to the chamber until the added liquid was equally distributed in the imaging chamber. We also checked the integrity of embryos which were treated with LY and embedded in agarose. We did not detect morphological defects under conditions similar to those in the SPIM chamber (Figure S9G). Following completion of imaging, embryos treated with LY were analyzed by *her4 in situ* hybridization to control for efficient inhibition of a Notch dependent *her* gene (Figure S9F).

BrdU treatment [modified after (Shepard et al., 2004)]: Up to 15 embryos were transferred into a 2 ml tube containing 0.2 mM PTU in E3. Embryos were incubated on ice for 15 min. After 15 min, the medium was replaced with 10 mM BrdU (Sigma), 15% DMSO in E3 and the embryos were incubated for another hour on ice. Embryos were washed four times with E3 and transferred to a petri dish containing 0.2 mM PTU in E3. After 1 hr at 28 °C, embryos were fixed in 4% paraformaldehyde overnight at 4 °C.

### 05 Analyses of mRNA and protein expression *in situ*

1. Whole-mount *in situ* hybridizations: Generation of probes. For the *her4* probe, which recognizes *her4.1* (ENSDART00000079274.4), *her4.2* (ENSDART00000137573.2), *her4.3* (ENSDART00000104209.4), *her4.4* ENSDART00000079265.6) and *her4.5* (ENSDART00000104206.4) a template was generated by using primers p159F and p160R and cloning into a pCRII-TOPO dual promotor vector. The *her6* probe template was cloned using p147F + p148R (ENSDART00000023613.9) and the *her9* probe template was cloned with p149F + p150R (ENSDART00000078936.4). *sox2* (ENSDART00000104493.5), *neurod1* (ENSDART00000011837.6), *neurog1* (ENSDART00000078563.5), *shha* (ENSDART00000149395.3) and *irx1b* [ENSDART00000079114.6; (Scholpp et al., 2009)] plasmids for probe generation have been previously published (www.zfin.org) or validated by sequencing. *mNeonGreen* probe: The *mNeonGreen* sequence (Shaner et al., 2013) was cloned into a Tol2Kit T7 middle entry vector. The vector was linearized with Apa1 (NEB) and transcribed with T7 RNA Polymerase (Thermo Fischer). Antisense RNA probes were generated using the DIG, DNP or Fluorescein RNA labelling kits from Roche (Merck, Germany).
2. Chromogenic whole-mount in situ hybridizations: Embryos were fixed at the indicated stages in 4% paraformaldehyde (PFA) overnight at 4 °C and dehydrated with methanol (25%, 50%, 75%, 100% MeOH in PBST). After rehydration, embryos were washed three times in PBST and digested in 10 µg/ml Proteinase K (AppliChem, 15 min treatment per 24 h of development). Embryos were washed and again fixed in 4% PFA for 20 min at room temperature. After five washing steps in PBST, embryos were incubated in 50% formamide, 5 x SSC, 5 mg/ml torula yeast RNA type IV, 50 µg/ml heparin, 0.1% Tween-20 at 65 °C for 3 to 4 h. Afterwards the digoxigenin-labeled antisense probes were added and the embryos were incubated at 65 °C overnight. At 65 °C, embryos were washed as follows: twice with 50% formamide in 2 x SSCT for 20 min each, twice with 25% formamide in 2 x SSCT for 20 min each, twice in 2 x SSCT for 20 min each, three times in 0.2 x SSCT for 20 min each, followed by washing steps at room temperature: once in 0.1 x SSCT in 0.5 x PBST for 10 min, twice in PBST for 10 min each. Embryos were incubated in 2% heat-inactivated goat serum (Vector Laboratories), 4 mg/ml BSA (AppliChem) in PBST for 2 to 3 h at room temperature. Afterwards the alkaline phosphatase-coupled anti-digoxigenin Fab fragment antibody (Roche) was added 1:3000, followed by an overnight incubation at 4 °C. Embryos were washed six times in PBST at room temperature for 20 min each, and three times for 15 min each in NTMT (100 mM NaCl, 100 mM Tris-HCl at pH 9.5, 50 mM MgCl_2_, 0.1% Tween-20). To the NTMT, BCIP (5-bromo-4-chloro-3-indolyl-phosphate, AppliChem) was added to final concentration of 0.18 mg/ml, and NBT (4-nitroblue tetrazolium chloride, AppliChem) to final concentration of 0.45 mg/ml. Staining reactions proceeded in the dark and were stopped by several washing steps in PBST containing 1 mM EDTA. Embryos were stored at 4 °C in 1 mM EDTA, 80% glycerol in PBST for later imaging using Leica MZ APO or Zeiss Axioskop2, Axiovision SE64, Rel.4.9.1 with AxioCam ICc1.
3. Fluorescent whole-mount in situ hybridizations: Embryos were treated as described above up to probe addition. The digoxigenin or dinitrophenol labeled antisense probes were added and the embryos were incubated at 65 °C overnight. Washes at 65 °C as above. After one washing step in TNT (100 mM Tris-HCl at pH 7.5, 150 mM NaCl, 0.5% Tween-20), embryos were blocked in TNT containing 1% Boehringer Block (Roche) for 2 to 3 h at room temperature. The horseradish peroxidase-coupled anti-digoxigenin Fab fragment antibody (Roche) was added 1:400 in blocking solution and embryos were incubated overnight at 4 °C. Embryos were washed eight times 15 min each in TNT, rinsed in 100 mM borate buffer pH 8.0 and incubated in staining mix: 2% dextrane sulfate supplemented 1% tyramide stock solution Alexa Fluor 488 (Thermo Fischer) containing 0.0015% H_2_O_2_, 112.5 µg/ml 4-iodphenole in 100 mM borate buffer pH 8.0 for 1 hr at room temperature. After three washing steps in TNT, embryos were incubated in 0.3% H_2_O_2_ in TNT for 30 min to inactivate peroxidase of first probe, and washed five times in TNT. Embryos were blocked in 1% blocking reagent (Roche) in 100 mM maleic acid, 150 mM NaCl at pH 7.5 for 2 to 3 h at room temperature. To this, the horseradish peroxidase-coupled anti-dinitrophenol Fab fragment antibody (Perkin-Elmer) was added 1:200 for overnight incubation at 4 °C. Again, embryos were washed eight times 15 min in TNT, rinsed in 100 mM borate buffer and incubated for 1 hr at room temperature in the staining mix, now containing 1% tyramide stock solution Alexa Fluor 555 (Thermo Fischer). Embryos were washed three times in TNT, another three times in PBST and, if nuclei needed to be stained, incubated overnight at room temperature in 1 µM TO-PRO-3 (Thermo Fischer) in PBST for nuclear staining. After three washing steps in PBST for 15 min each, embryos were stored at 4°C in 80% glycerol/PBS for later imaging.
4. *in situ* gene expression analysis using HCR probes: The HCR probes were obtained from Molecular Instruments (Los Angeles, USA) or designed for this study. Larvae were fixed in 4% paraformaldehyde (PFA) in PBS over night at 4 °C. For the HCR procedure, the “HCR-RNA-FISH (v3.0) Rev. 10” protocol for whole-mount zebrafish embryos and larvae from Molecular Instruments was used, with the following modifications: After rehydration, larvae were treated with proteinase K (10 µg/ml, AppliChem) in PBST (PBS with 0.1% Tween 20, AppliChem; pH 7.4) for 30 min. For the second overnight incubation with hairpins, 10 μg/ml DAPI (#D9542, Sigma-Aldrich) was added to the amplifier solution. After the last step of the protocol, the embryos were washed with 50% 5x SSCT (750 mM NaCl; 75 mM C_6_H_5_Na_3_O_7_, Carl Roth; 0.1% Tween 20; pH 7) / 50% PBST mix, followed by two 10 min PBS washes. Larvae were stored at 4°C in 80% glycerol/PBS for later imaging.
5. Generation of anti-Her6 antibody: The polyclonal anti-Her6 antibody was generated by immunization of rabbits with the zebrafish Her6 antigen peptide ADIMEKNSSSPVAATPASMNTTPD conjugated to KLH carrier by Davids Biotechnology (Regensburg, Germany) and affinity purification using the same peptide. The anti-Her6 antibody was validated based on correlation of stain pattern with published mRNA expression, and ability to recognize the Her6 protein expressed in *Tg(hsp70l:her6-FLAG)m1492* embryos upon heat-shock (Figure S3D).
6. Procedure for whole-mount immunofluorescence: Embryos were fixed overnight in 4% PFA at 4°C and directly processed for staining. Embryos were washed five times in PBST. For the Sox2 plus BrdU antibody staining shown in Figure 1A, an additional demasking step was performed: 5 min in 150mM TrisHCl pH 9.0 at room temperature followed by 15 min in TrisHCl pH 9.0 at 75°C; followed by two washes with H_2_O and a 1 hr incubation in 1 N HCl at room temperature, and finally two H_2_O and three PBST washing steps [modified from (Shepard et al., 2004)]. Embryos then were blocked for 30-60 min at room temperature in PBTD (1% BSA, 1% DMSO, 0.5% Triton X-100 in 1 x PBS pH 7.4) supplemented with 2% goat serum. The primary antibodies were diluted in PBTD + 2% goat serum and incubated 4 h at room temperature or at 4 °C overnight. All primary antibodies were diluted 1:400, except for the Her6 antibody in the *hsp:her6* overexpression experiment (Figure S3D) which was used 1:100. The primary antibodies were washed out six times 20 min in PBTD and the secondary antibodies were diluted 1:1000 in PBTD + 2% goat serum, and incubated 4 h at room temperature or at 4 °C overnight (anti-mouse 488 A11001 from Thermo Fisher, anti-rabbit 555 A21430 from Thermo Fisher). Embryos were washed six times 20 min each in PBTD and then transferred to 80% glycerol in PBS.

Antibodies used:

Anti-SOX2 antibody [20G5], Abcam, ab171380, LOT GR3253929-5

BrdU Polyclonal Antibody, Thermo Fisher Scientific, Catalog Number PA5-32256, LOT UA2696578B

Anti-Her6 from rabbit. The polyclonal antibody was generated for this study by immunization with a zebrafish her6 antigen peptide (see above).

Anti-Digoxigenin-AP, Fab fragments polyclonal, Merck, LOT 16646822 REF 11093274910

Anti-Digoxigenin-POD, Fab fragments polyclonal REF 11207733910 LOT 11650300

Anti-DNP HRP Conjugate, Perkin Elmer, FP1129 LOT 2590435

### 06 Microscopy and data collection

#### 06.1 Confocal microscopy

##### Confocal imaging

An inverted microscope with LSM-880 airyscan (Zeiss) was used with a 40x LD LCI Plan Apochromat 40x/1.2 autocorr (Zeiss) objective for imaging of all fluorescent *in situ* hybridizations and antibody stainings. HCR-samples were imaged on an upright Zeiss LSM-880 fast-airyscan setup using a LD LCI Plan-Apochromat 25x/0.8 Imm Korr DIC (UV) VIS-IR objective. Images shown in Supplementary Figure S1C were acquired as two tiled z-stacks and subsequently stitched. Live imaging of *Tg(her6:her6-mNeonGreen)^m1366^* shown in Figure 1E was performed on the same upright LSM-880 fast-airyscan setup using the W Plan-Apochromat 20x/1.0 Corr DIC M27 75mm objective and acquired in two tiles. For recordings with the inverted Zeiss LSM-880, the ZEN 2.3 SP1 FP3 (black) version 14.0.22.201 was used. All embryos were mounted in 1% standard agarose (Bioron) in 80% glycerol/H_2_O. All stacks were recorded with the Airyscan detector in the optimal mode and processed with the ZEN black software using the airyscan algorithm (v. 2.3 SP1 Zeiss). All pictures shown are single z-planes from stacks except for the in situ hybridization images of *her6* together with *sox2* (Figure 2B) and of *her6* together with *her6-mNeonGreen* (Figure S4E), for which a maximum intensity projection of 5 z-planes corresponding to 5 µm thickness were generated using ZEN black. Sagittal reslices in Figure S1C are average intensity projections over XX pixels corresponding to YY µm thickness centered around the midline ventricle.

#### 06.2 Selective plane illumination microscopy (SPIM)

##### SPIM live imaging sample preparation

Double transgenic zebrafish embryos homozygous for *Tg*(*her6:her6-mNeonGreen*)*m1366* and heterozygous for *Tg(beta-actin:NLS-mCardinal)m1508* were incubated in E3 plus 0.2 mM PTU until mounting. Embryos were anaesthetized with 1 mM tricaine, dechorionated by hand, and mounted with 0.4-1.0% low melting agarose (Sigma) in straight vertical (e01 and e02) or 90° bent glass capillaries (all other embryos) of 0.55 mm or 1.15 mm inner diameter (Zeiss, black respectively green coded capillaries). 90° bent capillaries enabled dorsal views onto the thalamus, and were generated by heating the glass capillaries and bending their tip. The low melting agarose was allowed to dry for about 10 min, and then the agarose in the end of the capillary was expelled from the capillary and cut off right before the embryo. The SPIM chamber was filled with E3 medium supplemented with 1mM MS-222. Embryos were mounted in the SPIM chamber in an orientation to avoid light sheet passing through the retina before imaging the thalamic proliferation zones. Temperature during imaging with the Lightsheet.Z1 (Zeiss) was held constant at 28.5 °C using the Z1 incubation chamber temperature control unit (TempModule LSFM).

##### SPIM Image Acquisition

We decided to use Selective Plane Illumination Microscopy (SPIM) for its higher imaging speed, and because it employs light of lower intensity than conventional LSM, suggesting less light induced damage to the imaged fluorophores and cells. Imaging was performed with the Lightsheet.Z1 microscope (Zeiss). mNeonGreen was imaged using the 488 nm (30 mW) laser at 2.5% intensity for all embryos. mCardinal was imaged using the low-level excitation with the same 488 nm laser line, except for e02 in which the 561 nm (20 mW) laser at 1.5% was used. These settings were optimized to minimize fluorophore bleaching. Both channels were recorded simultaneously by two cameras with an illumination time of 200 ms per z-plane. A W-Plan-Apochromat 20x/1.0Corr objective (Zeiss) was used together with the 10x/0.2 illumination objective (Zeiss 400900-9000-710). Two PCO.edge 5.5 with sCMOS sensor (5.5 MP, 2560 x 2160) cameras were used for all SPIM recordings. A LBF 405/488/561 (Zeiss filter number 404900-9100-000) notch filter was used together with a beam splitter ssLP 560 followed by a BP 505-545 bandpass filter to record green emission. The red emission was recorded simultaneously using a LP 585 longpass filter. This causes a minor bleed through of the green signal into the red detection channel (about 3% bleed through of mNeonGreen emission compared to 88% detection of mCardinal; see: https://www.fpbase.org/spectra/ output collection efficiency for mNeonGreen, mCardinal and Omega 585ALP filter). The red NLS-mCardinal signal was predominantly used for tracking of cell nuclei, which is not affected by green signal bleed through. A single sided, Gaussian light sheet (2.13 µm thickness) with pivot scan was used. Imaging intervals between each stack of a time series were 4.5, 9 or 18 min.

#### 06.3 Determination of SPIM time series recording intervals

To evaluate potential physical effects of the measurement procedure on measured oscillation periods, we implemented a control. Excitation light may damage protein fluorophores and initiate their proteolytic decay, so when using fluorescent fusions to assess protein dynamics and stability, it is important to determine whether imaging affects the protein of interest. We therefore examined Her6-mNG dynamics by imaging at shorter (4.5-min instead of 9-min) or longer (18-min) time intervals. Oscillation periods were only significantly different when 18-min intervals (3.7 h SD 2.6) and 4.5-min intervals (3.11 h SD 0.5) were compared (p=0.02; Figure 3A, Figure S8E). We conclude that fluorescent imaging may indeed reduce Her6-mNG stability, and the more rapid decay may have resulted in shorter oscillation periods. However, given the small difference between 18- and 9-min intervals (p=0.11), we deem that our observed periods closely approximate those of endogenous Her6 oscillations, and that light damage did not significantly affect Her6-mNG expression dynamics under our conditions.

#### 07.1 SPIM data processing, object tracking and quantification

##### SPIM Data Image Preprocessing

Recorded czi files from ZEN 2014 SP1 (v9.2.6.54) were deconvolved with Huygens Professional (v18.1 SVI). An automated absolute background estimation (area radius set to 0.7 micron) was used (no relative background was used for the deconvolution). For the parameter set up, excitation wavelengths were set to 488 nm and the emission wavelength was set to 521 nm for the green channel and to 569 nm for the red channel. A SPIM optimized deconvolution with a theoretical point spread function was performed (settings: Maximum iterations, 40; signal to noise ratio, 20; quality threshold, 0.1; iteration mode, “optimized”; brick layout, “auto”). Output data was saved as ics2 files and loaded into arivis Vision4D (v2.12.6, v3.0.1, v3.01, v3.1.2 and v3.1.3) as a time series.

##### Fluorescence measurements

In arivis Vision4D three 3D spheres were placed into each cell nucleus for each time point. Spheres were placed in the center of the nucleus and should overlap as little as possible. The red channel was used to place spheres when the green signal was too weak. Each sphere has a diameter of 1.5 µm that comprises 5 z-planes and contains 228 voxels (Figure S5E). The average intensity of these 228 voxels within each sphere (grey measurement dots in Figure S5D) was determined. The average intensity values from each of the three spheres were used to calculate an average intensity value for this nucleus (black line in Figure S5D).

##### Quality control of sphere measurements

Positioning of all spheres was validated a second time by another person, and corrected if necessary. In addition, the standard deviation (sd) of intensity data from each of the 228 voxel within a single sphere varies depending on the position of the sphere. sd is low when the intensities of the pixels are similar, for example when it is placed correctly in the middle of the cell (Figure S5E spheres labelled S1, S2 and S3). sd is high when spheres are misplaced (Figure S5E spheres labelled S4 and S5). The normalized standard deviation (standard deviation / mean intensity x 100) was calculated, and misplaced spheres have been identified by the normalized standard deviation exceeding those of the misplaced sphere S4 (Figure S5F, red “x” shows data for all sample spheres including S4 and S5). Standard deviations for all spheres from embryos em07, em12, em13 and em15 are shown in Figure S3F. Spheres at the border of the stack of em07 were removed from the analysis, because black voxels from outside of the stack would be included. Single spheres with a normalized standard deviation > sd of S4 were identified and checked again whether they were placed correctly: if they had been misplaced, a new sphere was generated, if they were properly placed, the data were considered to be validated.

##### Data export

For each sphere, name tags (embryo e, cell number c, and sphere identifier a/b/c: eNNcNNNa), time, mean voxel intensities of red and green channels, and the mean/sd of voxel intensities were exported directly from arivis Vision4D as excel files. Cell divisions are indicated in the tag. For documenting dividing cells, both daughter cells were tracked (Figure 2B, D). For mathematical identification of oscillations, only the longest continuous tracks without a mitosis were considered, because cell divisions influence signal intensities in the nucleus. All tracks shorter than 400 min were excluded from the analysis. A script was used to organize the data and calculate for each time point the average intensity (average of 3 spheres and standard deviation) for both channels in each individual nucleus.

#### 07.2 Comparison of relative fluorescence intensities by normalization to NLS-mCardinal fluorescence

Raw fluorescence intensity values for em07 were exported from arivis Vision4D software. For each tracked cell, three nuclear spheres (a,b,c) were quantified per time point. For each time point and cell, the mean green fluorescence intensity (Her6-mNeonGreen) was calculated as the average of sphere a, b, c. The mean red fluorescence intensity (mCardinal-NLS) was calculated analogously. For normalization of the relative expression values independent of imaging depth and variations in acquisition, we argued that nuclear mCardinal-NLS, which should be expressed at similar levels in each cell nucleus, may be used for normalization. The minor bleed through of mNeonGreen into the red channel (less than 5% of signal, see Methods 06) should not significantly affect the ratio. Her6-mNG fluorescence intensities were normalized to the red nuclear reference signal by dividing the green mean intensity by the red mean intensity for each cell. The ratio should eliminate many of the variables intrinsic to SPIM time series analysis, and provide relative Her6 protein level per cell that may be compared between all cells. To compare basal expression levels independent of long-term drift, only the first 30 time points were analyzed. Given the acquisition interval of 9 minutes, this corresponds to 270 minutes of imaging. For each cell, the mean normalized Her6-mNG fluorescence intensity across these 30 time points was calculated. All cells were assigned to the anatomical regions Telencephalon, ZLI and rTh(p). Only cells with a non-ambiguous assignment were used in the analysis. Statistical significance was assessed using two-tailed Student′s t-tests for independent samples.

### 08 Quantification of *her6* expression in inhibitor and heat-shock experiments

#### 08.1 Quantification of staining intensities of *in situ*-stained embryos

Four embryos of each condition were imaged in transmitted light recording Z-stacks in lateral views (Zeiss Axioplan2 with 10x /0.3 Plan-NEOFLUAR 440330, Axiovision SE64, Rel.4.9.1 with AxioCam ICc 1). The medial z plane was used for the analysis. Images were opened in FIJI (Figure S9E’), converted into 8-bit greyscale (Figure S9E’’) and inverted (Figure S9E’’’). For the *her6* and *her4* expression analysis, a single polygon was drawn around the thalamic and prethalamic expression domain of the respective gene and saved as template using the ‘ROI manager’ analysis tool (Figure S9E’’’’). For each image, this ROI was moved to the thalamic and prethalamic expression domain. With the measurement tool of the FIJI ROI manager, the mean staining intensity of this area was determined and exported to Microsoft Excel. The mean and its standard error were calculated based on four analyzed embryos for each condition (Figure S9C,D). For the her4 expression analysis, we used LY-411575 treated embryos that do not express her4 to estimate baseline intensities of head tissue in this assay. The baseline (Figure S9C) was calculated by averaging the intensity of the whole picture for *her4* stained embryos, which had been treated with LY-411575 for 6 h (Fig. S9A, 6 h LY treated panel). This procedure was done for four individual embryos and the average was plotted as the dashed line in Figure S9C.

#### 08.2 Generation of time series for *her6* mRNA decay rate measurements by qPCR after heat shock-induced Her6 overexpression

*Tg(hsp70l:her6-FLAG)^m1492^* heterozygous fish (carrying *clmc2:EGFP* fluorescent heart marker) were crossed with homozygous *Tg(her6:her6-mNeonGreen)^m1366^.* Embryos from this cross were sorted for *Tg(hsp70l:her6-FLAG)* based on their fluorescent heart marker; embryos without fluorescent heart marker expression were kept as a control.

##### Experimental design

RNA samples were obtained in triplicates from *Tg(hsp70l:her6-FLAG)* experimental and transgene-negative control embryo pools 15 min before heat shock started, at the end of the heat shock, and then every 15 min until 135 min after heats hock. Non-heat shocked control embryos (3 x 5 embryos each for the *Tg(hsp:her6-FLAG)* - 15 min experimental sample and for the *Tg(hsp:her6-FLAG)* – 15 min negative control sample) were grown in E3 medium and placed 15 min before 72 hpf into 150µl RNAlater (Ambion, AM7024) for RNA extraction. All other *Tg(hsp:her6-FLAG)* experimental embryos and control embryos without this transgene were heat shocked at 72 hpf. The heat shock was performed for 15 min in 39.5°C pre-warmed E3 medium in 50 ml tubes placed in a temperature-controlled water bath. Right after heat shock, 3 x 5 embryos each carrying *Tg(hsp:her6-FLAG)* “Tg_HS+0min” and *Tg(hsp:her6-FLAG)* negative embryos “WT_HS+0min” were placed into RNAlater. Successively, every 15 min until 135 min past heat shock, 3 Tg and 3 WT samples each containing 5 embryos each were placed into RNA later.

##### RNA extraction

Samples containing 5 embryos each in RNAlater were stored at −20°C. Total RNA was extracted from 3-5 embryos recovered form RNAlater (differences in embryo numbers were corrected by using similar amounts of mRNA for each cDNA synthesis reaction, and normalizing qPCR using *rpl5b* reference gene expression). For RNA isolation, the RNeasy Mini Kit (Qiagen, 74106) and QIAshredder columns (Qiagen, 79656) were used. The samples were homogenized in 1% β-mercaptoethanol in 600 μl RLT buffer (Qiagen, 74106) by application of physical force. Physical force was applied by pipetting the embryos several times through a 0.6 mm diameter canule using a syringe (Sterican canule Luer-lok 0,60×30 mm size 14 blue from B. BRAUN). The suspension was transferred to a QIAshredder column and centrifuged for 1 min at 10,000 g. Then 700 μl 70% ethanol was added to the flow through and transferred to RNeasy Mini spin columns. After centrifugation (15 s, 10,000 g), 350 μl RW1 buffer (Qiagen, 74106) was added to the membrane and centrifuged. Digestion with DNase I was performed by adding 80 μl of the DNase I incubation mix (consisting of 10 μl DNase I stock solution with a concentration of 2.73 Kunitz units/μl and 70 μl RDD buffer; Qiagen, 74106) to the RNeasy spin column membrane for 15 min to remove DNA from the membrane. After two more washing steps according to the manufacturer’s protocol, the column was dried by centrifugation for an additional 2 min. Total RNA was eluted by adding 30 μl H_2_O to the filter membrane. Tubes were incubated for 1 min at RT and centrifuged for 1 min at 10,000 g. The RNA concentrations were determined using a NanoDrop photometer.

We used 100 to 250 ng total RNA each as template for reverse transcription into cDNA. cDNA was generated with the SuperScript III RT Kit (Invitrogen) according to the manufacturer’s protocol using oligo(dT)12-18 (0.5 μg/μl) primers. After heat-inactivation of the reaction (70°C for 15min), the RNA was removed from the cDNA by addition of 1 µl *E.Coli* RNAseH and incubation at 37°C for 20 min.

The qPCR was performed with the SYBR Green Supermix (Sso Advanced Universal, Bio-Rad). Each 10 μl qPCR mix contained 5 μl 2× SYBR Green Supermix, 3µl H_2_O, 1 μl primer mix (5 μM final concentration of each primer) and 1 μl cDNA. The PCR plate was sealed and centrifuged for 10 min at 1,000 g. qPCRs were performed using the LightCycler (Roche). For all qPCRs the following qPCR program was used: step 1, 95°C for 30s; step 2, 95°C for 15s; step 3, 60°C for 50sec, step 2 and 3 repeated for 40 cycles; step 4, 95°C 60s; step 6, 40°C 60s; step 7 from 65°C to 95°C with 0.07°C per second. The data analysis was performed as in Taylor et al. (2019), and *rpl5b* expression was used as reference gene.

For all samples, the number of biological replicates is three. For each biological replicate, the number of technical replicates is three, unless stated otherwise. Statistical analyses were carried out using Microsoft Excel 2016.

### 09 Anatomical location of tracked cells

For all analyzed cells of embryos em06, em07, em12, em13 and em15 (day 3 of imaging), the location in the anatomical regions rostral thalamus(p), ZLI or telencephalon was determined by analyzing their position manually in arivis Vision4D. Cells that could not be unambiguously assigned to one of these anatomical regions were marked as not assigned. All cells in embryos em02, em19, em20, em22, em23, em27 were tracked in the ZLI or rTh(p). Cells in em01 (day 2 of imaging) were not assigned to thalamic region because morphological landmarks are missing during the second day of development, and assigned to the diencephalon.

### 10 Heatmaps and clustering, statistical tests of oscillation frequencies

Heatmaps were generated with Heatmapper expression module (Babicki et al., 2016) at http://heatmapper.ca. Heatmaps show detrended Her6-mNeonGreen intensity data. For the clustered heatmap (Figure 2E), only the first 80 time points of tracks longer than 12 h were considered (155 cells total from embryos e02, e06, e07, e13, e15). The starting points of these tracks were aligned to 0, ignoring the actual developmental time of recording. Cells from all embryos documenting day 2 or day 3 of development with 9 min imaging intervals were subjected to clustering using the Pearsońs Distance measurement method with centroid linkage (Scale type column, z-score = standard deviations of the mean of one cell). For the real-time heatmap (Figure S5G), all cells imaged during day 3, and separately day 2 of development, were aligned according to their developmental age, and no clustering was performed. For the heatmap of em07 cells were organized by mathematical classification (Figure S8G), the first 90 time points of the time series (3480 to 4281 min post fertilization) are shown (no clustering applied, cells shown in numerical order).

Comparison of oscillation frequencies and periods was performed using Wilcoxon-Ranksum test (α = 0.01).

### 11 Gaussian Process Regression

Gaussian processes (GP) are stochastic processes for which every finite subset of random variables is distributed as a multivariate Gaussian distribution. A GP is fully specified by its mean and covariance function *m(t)* and *k(t, t’)*, respectively. When subtracting the mean from the data, *m(t)* can assumed to be zero. The covariance function encodes the information in which way and to what extent two data points are correlated. To ensure Gaussianity, the data is transformed by replacing each data point of the measured ones by a realization of a normally distributed random variable with the same rank (Schreiber and Schmitz, 2000). This process is called Gaussianization.

Experimental data *y(t), t = 1, 2,…, N_d_* can be linked to the underlying dynamics *f(t)* by *y(t) = f(t) + ε(t), ε(t) ∼ N(0,σ^2^)*, where the measurement errors *ε(t)* are normally distributed with a variance of *σ^2^*. Instead of specifying the functional form of the underlying dynamics, for non-parametric regression it is assumed to be drawn from a GP:

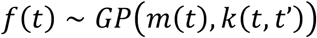

The mean and covariance functions include a number of parameters (hyperparameters) that must be chosen such that the probability of the data y given the GP model is maximal. This can be achieved by optimizing the logarithmized marginal likelihood function of the data ***y*** at times ***t***

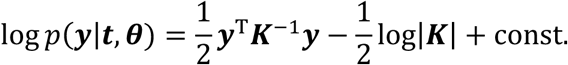

with respect to the hyperparameters ***θ***, where ***K*** represents the (*N_d_* × *N_d_*) covariance matrix, which is made up of the covariance function evaluated at the respective measurement time points. The marginal likelihood function for non-parametric GP regression encourages the compatibility between model and data in its first term and by construction penalizes model complexity in the second.

#### 11.1 Removing long-term trends superimposed to biological signals

Due to experimental or biological effects such as bleaching, cell growth or shrinkage, or the underlying gene expression dynamics, aperiodic long-term trends may be superimposed to the data, which can dominate the signal of interest. To be able to analyze the underlying signal of interest, the data has to be detrended (Phillips et al., 2017). GP regression poses a good alternative to parametric regression since the functional form of the trend is not known a priori. We use the infinitely differentiable squared-exponential covariance function

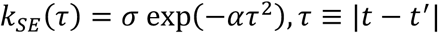

which leads to smooth time courses. The hyperparameters σ and α are found by optimizing the log-likelihood function using a multi-start approach. To ensure the GP does not capture the biologically relevant oscillations, the parameter space is constrained such that the characteristic time scale 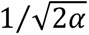 cannot decrease below approx. 2300 min, which is far above the time it takes a neural cell to progress from mitosis to differentiation in zebrafish larvae. Fitting of the GP is performed using the GPML toolbox for MATLAB (Rasmussen and Nickisch, 2010).

#### 11.2 Oscillatory classification

System size expansion (Van Kampen, 1992) together with the linear noise approximation (Elf and Ehrenberg, 2003) can be used to identify GPs, specifically the Ornstein-Uhlenbeck process, as an approximation for single cell gene expression dynamics (Phillips et al., 2017). The covariance function can be expressed in terms of the eigenvalues {*λi*} of the Jacobian of the underlying reaction rate functions:

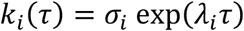

Depending on the imaginary part of the eigenvalue, this function can lead to either aperiodic fluctuations if the eigenvalue is purely real, or oscillating dynamics if it exhibits an imaginary part unequal zero. We distinguish these two cases by requiring a real decay constant α and expressing the oscillating behavior explicitly via a cosine term:

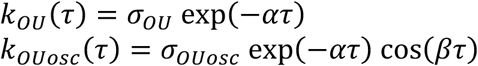

These auto-covariance functions pose two different GP models whose compatibility with the data can be compared by means of a likelihood ratio test. We use a multi-start approach for optimization with the standard deviation of the time series as an initial guess for the noise hyperparameters *σOU* and *σOUosc*, and sample initial guesses for the remaining hyperparameters from the region 1/250 min^−1^ < *α* < 1 min^−1^, 0min < 2π/*β* < 400min. We optimize in log-space which enforces positivity of parameters, and constrain the period of oscillations, i.e. 2π/*β*, to values 50 min < 2π/*β* < 600 min, which appears a reasonable range based on initial visual inspection of primary data (see Figure 2E).

To evaluate optimizer convergence, we analyze the negative log-likelihood of each optimization run ascendingly sorted which results in the so-called likelihood waterfall plot (Loos et al., 2018). We observe that most optimization runs converge to one of very few optima in parameter space, indicating a good optimization performance and reliable results. For both GP models, we pick the model fits that lead to minimal negative log-likelihood values among all multi-start runs and compare them using the test statistic

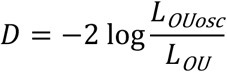

Because *k_OU_* follows from *k_OUosc_* when *β* is equal to zero and therefore is located on the boundary of parameter space, Wilk’s theorem is not directly applicable here. Instead, *D* follows a 50:50 mixture of a 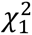 and a 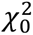 distribution (Self and Liang, 1987). For every time series, we fit both models and calculate the corresponding numerical value of the test statistic *D*. We derive a threshold for *D* from the significance level *α* = 0.01 below which the GP model with the *k_OU_* covariance function represents a valid simplification of the GP model with *k_OUosc_*. The time series for which this simplification is not allowed are then considered oscillatory and vice versa.

### 12 Hilbert Transformation

A signal *z(t)* is called analytic if its Fourier transform *Z*(*ω*) does not possess any negative-frequency components, and can therefore be expressed as

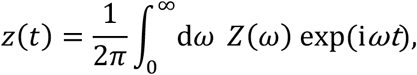

where *Z*(*ω*) ∈ ℂ defines amplitude and phase at every frequency. To obtain an analytical version of a real-valued signal *u(t),* one has to add a 90°-phase-shifted version of the signal, *v(t),* as an imaginary part, which can be obtained by means of the Hilbert transform:

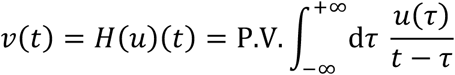

Here, P.V. is indicating that the integral has to be evaluated as a Cauchy principal value. The analytical signal is then given by *z(t) = u(t)* + i*v(t)* ∈ ℂ and can also be represented in polar form via time-dependent amplitude *A(t)* and phase *Φ(t):*

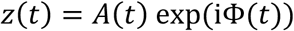

*A(t)* and *Φ(t)* can now be regarded as instantaneous amplitude and instantaneous phase. Since frequency is defined as the derivative of the phase, we can also define the instantaneous frequency as

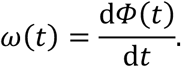

This analysis is meaningful if *A(t)* and *ω*(*t*) change slowly with time.

Consider a sinusoidal oscillating function in the time domain *y*_1_(*t*) = *Ã* sin(2*πt*/*T*) with constant amplitude *Ã* and period *T* (Figure S8C). Modulating the amplitude of *y_1_* with a linear function *A*(*t*) = *Ã* − *kt*, i.e. introducing a time-dependence to the amplitude, leads to a linearly damped oscillation *y*_2_(*t*) = (*Ã* − *kt*) sin(2 *πt*⁄*T*). To characterize the change in the instantaneous amplitude, one can calculate the slope of *A(t)*, which is given *Δ A*⁄*Δ t* = −*k* in this example. Similarly, if we instead let the period of oscillation vary with time, e.g. by *T*(*t*) = 1⁄(ℎ_1_ + ℎ_2_*t*), we obtain the function *y*_3_(*t*) = *Ã* sin(2*π*(ℎ_1_ + ℎ_2_*t*)) (Figure S8D). Since in our example the time-dependence of the period is more naturally captured in the frequency *f*(*t*) = 1/*T*(*t*) = ℎ_1_ + ℎ_2_*t*, we calculate the slope *Δ f*⁄*Δ t* = ℎ_2_ to quantify how strong the variation is.

### 13 Surrogate Data

Surrogate data mimicking the properties of stationary, linear Gaussian processes were suggested (Theiler et al., 1992) to substantiate the results of analysis methods of non-linear dynamics as fractal dimensions and Lyapunov exponents (Kurths and Herzel, 1987). A fundamental property of stationary, linear, Gaussian processes is that the Fourier transform for distinct frequencies of all orders are uncorrelated. Non-linear processes are characterized by their higher order correlation structure, e.g. the bi-spectrum in lowest order (Greb and Rusbridge, 1988). Surrogate data displaying only the second order, i.e. linear, properties in the time domain of the underlying process are generated by the following procedure:

i. Fourier transformation the data of every cell
ii. Randomization of the phases of the Fourier transform in order to destroy any kind of higher order correlations
iii. Back-transforming into the time domain using the inverse Fourier transform

This procedure ensures that the autospectra of the surrogate data are identical to the autospectra of the original data. In the field of non-linear dynamics, the application of surrogate data was criticized (Timmer, 2000) since first, mathematically strictly speaking, linear, stationary, Gaussian processes might be rare in nature and, second, the composite null hypothesis renders it difficult to draw conclusions about a specific alternative hypothesis.

In principle, for these applications, the same fundamental critics as in (Timmer, 2000) applies. To deal with Gaussianity, we applied Gaussianization to the data (Section 11). A hallmark of non-linearity are higher harmonics in the spectrum. Investigating the spectra of the experimental data shows that higher harmonics are rarely present.

We apply the surrogate data approach to exclude non-stationary time series from the discrimination analysis based on Gaussian processes and spectral analysis between first order aperiodic and second order oscillatory processes (Section 15). Additionally, we use it for determining finite size critical values for a test for significant coherence different from zero (Section 16.3), as well as synchronization between the time series of different cells in order to test for possible coupling between the cells (Section 17).

### 14 Autoregressive Models

Autoregressive (AR) processes are a class of stochastic processes for which the value of a time series *x(t)* depends linearly on its values at previous times *x(t−1), x(t−2),…, x(t - p)*, according to the order *p* of the process:

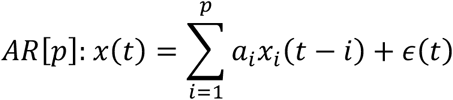

Noise is assumed to be normally distributed with mean zero, i.e. *ε(t) ∼ N(0,σ^2^).* AR processes can be interpreted as the superposition of relaxators and damped oscillators (Honerkamp, 1993). The AR[1] process corresponds to a simple relaxator as it can be thought of as the discretization of a first-order linear differential equation with a driving noise term, i.e. an Ornstein-Uhlenbeck process. On the other hand, the AR[2] process is able to produce oscillating time series for a specific choice of parameters {*ai*}.

#### 14.1 Second-order autoregressive models can lead to oscillations

An AR[2] process can be rewritten as first order difference equations by introducing a new variable ***z***(t) = (*x(t), x(t − 1)*)^T^:

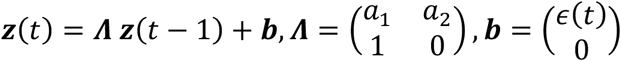

The general solution of this inhomogeneous matrix difference equation is given by the superposition of a particular solution of the inhomogeneous equation and the general solution of the homogeneous equation. Assuming ***Λ*** is diagonalizable we can write ***Λ*** = ***PDP***^−1^, where ***D*** is the diagonal matrix of eigenvalues *{λi}* of ***Λ***. Applying the transformation ***z***(t) → ***P u***(t) yields

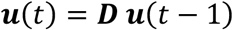

which has the solution ***u****i(t) = **c**i λ^t^_i_*. Because the elements of ***z****(t)* are linear combinations of the ***u****i(t)*, we can write the general solution of the homogeneous equation as

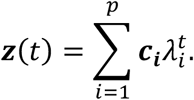

If the eigenvalues {*λi*} are complex, this leads to oscillating dynamics. This is the case for *a_1_^2^* < 4*a_2_* for the AR[2] process. The AR parameters can be related to period T and relaxation time τ of a damped oscillator (Timmer et al., 1998) via

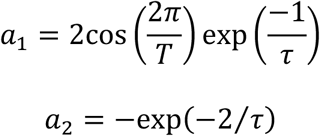

The power spectra (Section 16) of AR processes are given by (Box and Jenkins, 1976):

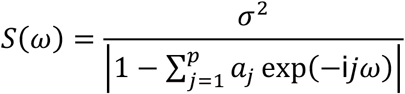

In contrast to the AR[1] process, an AR[2] process can have a distinct peak at *ω* > 0 in its spectrum corresponding to the oscillations occurring under conditions described above. More than one peak in the spectrum can occur for higher order processes that include more complicated dynamics.

### 15 Testing stationarity of time series

The applied Gaussian processes and spectral analyses methods to investigate oscillatory behavior and interactions between cells require stationarity of the underlying processes. Mathematically, this means that all joint distributions of the underlying process must be time-invariant. Essentially, it means that the underlying dynamics do not change with time, which might lead to uncontrollable results for the methods applied which assume stationarity.

Non-stationarity might come in different flavors: For oscillatory processes, the amplitude or the frequency might change over time. While changes in the amplitude do not affect the results of the methods applied in this work overly strongly, time-dependent frequencies might do so. Following Xiao et al. 2007, we use a surrogate data based approach to identify non-stationary time series: We generated 3000 surrogate data sets for every time series and calculated the instantaneous phase (Section 12). We regarded an experimental time series as a realization of a non-stationary process if its phase development deviates on a 1% level from the phase developments of the surrogate data.

To investigate the performance of this method under the null hypothesis, i.e. a stationary process, 1000 realizations of an AR[2] process with *T* = 270 min and *τ* = 63 min were simulated. Applying the same analysis with a significance level *α* = 1% to the simulated data reveals that the portion of false positives produced by this method is 12% and therefore larger than expected. We therefore conclude that the proposed method does not represent a formal statistical test but nevertheless provides a statistical conservative reproducible prescript.

### 16 Spectral analysis

#### 16.1 Auto-spectrum

The data are further examined by means of spectral analysis. The auto-spectrum *S(ω)* of a time series *x(t)* is defined as the expectation value of the squared modulus of its Fourier transform *X(ω)*, the periodogram:

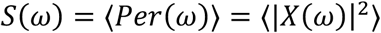

Since the periodogram is 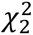 distributed and its standard deviation does not decrease when adding more data (Brockwell and Davis, 1987), it not a consistent estimator for the true auto-spectrum. Due to spectral leakage, the periodogram is a biased estimator. Since the true auto-spectrum of a mixing process is smooth, the periodogram can be convoluted with a normalized smoothening window function *w(j)* to acquire a suitable estimator:

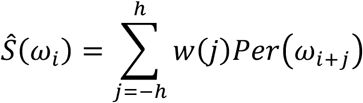

To calculate the auto-spectrum, we apply biologically justified cuts to the detrended data. The data are normalized to a standard deviation of one and a mean of zero, then padded with zeros to a length of 2^8^ = 256, and undergo the fast Fourier transform (FFT) algorithm implemented in the MATLAB release R2019a. Due to the symmetry *X*(−*ω*) = *X*^∗^(*ω*), only the positive frequency contributions have to be considered. The spectral power at zero-frequency is left out in further calculations because it vanished since the mean was subtracted from every time series beforehand. Including this value would therefore create an artificial peak in the auto-spectrum near zero frequency. The Bartlett window is chosen for smoothing and its width is determined individually for each frequency (Timmer et al., 1996), see Table Methods 16.1 (below). The confidence intervals to a significance threshold of *α* around *S(ωi)* are given by (Brockwell and Davis, 1987)

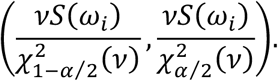

The degrees of freedom can be calculated from the window function by 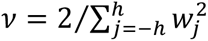 (Timmer et al., 1996).

**Table Methods 16.1:**
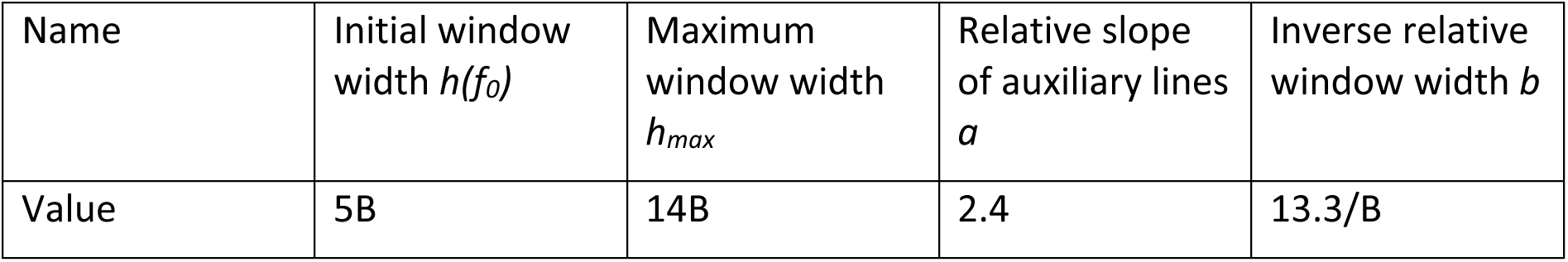
Parameter values for the adaptive smoothing algorithm presented in Timmer et al. 1996. B refers to the frequency bin size which depends on the sampling frequency f_s_ and the length of the (padded) time series B as B = f_s_/N.

The smoothed auto-spectrum is searched for peaks whose prominence exceed the 99% confidence interval. The prominence is a measure for how much a peak stands out from the overall frequency spectrum. It is defined as the height above the lowest value of the spectrum in a search range left and right of the peak of interest. The search range is limited by nearest values of the spectrum larger than the peak of interest, or the end of the spectrum if there is no such value. If a peak is present in the biologically relevant frequency range 1/435 min^-1^ < *ω*/2*π* < 1/60 min^−1^, the time series is considered oscillatory. Peaks that fall below the lower bound mentioned above are regarded as belonging to *ω* = 0 and therefore indicate a non-oscillating first-order process.

#### 16.2 Cross-spectrum

The complex-valued cross-spectrum of two time series *x(t*) and *y(t)* is defined analogously to the auto-spectrum via the Fourier transform

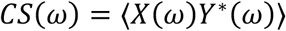

and is estimated via the cross-periodogram *X(ω)Y ^∗^(ω)*. The width of the smoothing window for the cross-spectrum is determined by the maximum of the window functions for the two corresponding auto-spectra at the respective frequency bin. Other than that, the calculation is identical to that of the auto-spectrum.

An important quantity concerning the relation between two processes is provided by the coherence, which measures their linear predictability on a scale between zero and one:

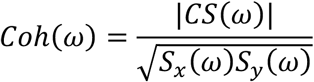

Adjacent cells might be coupled via the Delta/notch signalling pathway. Therefore, between such cells, we expect HER6 concentration time series to be linearly predictable, i.e. coherent. To infer coupling, we calculate the auto-spectra of the time series of all cells in one embryo and search for significant peaks as described above. If a pair of cells is found which has peaks with a frequency distance smaller than 5·10^−4^ min^−1^, we investigate whether those peaks originate from an interaction between the two cells by calculating the coherence between the corresponding time series (Figure S8G). The asymptotic critical value for non-zero coherence is given by (Timmer et al., 1998)

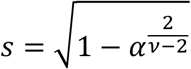

with significance level *α* and *ν* degrees of freedom. This results holds asymptotically for stationary so-called mixing processes. The vivid meaning of mixing is that mixing processes lose the information about their initial conditions. Harmonic oscillators with sine and cosine as solutions and limit cycles on their attractor are strictly periodic and not mixing and therefore never lose the information about their initial conditions. Coherence analysis which investigates linear predictability is not applicable for this kind of processes since it will always result in a coherence of one for processes with the same frequency. All other stationary processes, no matter whether they are deterministic or stochastic, exhibit a typical time scale for losing the information about their initial conditions, which is the called mixing time. Typically, this is the characteristic time scale of the exponentially decaying auto-covariance function introduced in Section 11.2. From a data analysis point of view, the asymptotics is reached when the data comprise at least around 20 times the data points of their mixing time. This condition is not satisfied by the present experimental data because often the mixing time is in the order of the length of the time series. This holds especially for those time series that could be modelled by an oscillatory deterministic differential equation (Section 18).

Applying the asymptotic results under this violation of the condition of sufficient data will lead to false positive results since one would be too close to the non-mixing situation. Thus, we applied the surrogate data approach to infer critical values for the test for coherence for this finite size condition. Therefore, we generate 1000 surrogate data and calculate the 99% percentile of the coherence.

#### 16.3 Power-of-the-test simulations

The coherence analysis between the time series of different cells within one embryo did not lead to more positive results than the expected false positive rate for the chosen significance level ɑ, which suggests cell-autonomous behavior. Of course, the rule „Absence of evidence is no evidence for absence” applies. To investigate how strong an interaction between cells must have been in order to be able to detect it by coherence analysis, a *power-of-the-test* simulation study was performed. Therefore, data for two AR[2] processes (Section 14) *x(t)* and *y(t)* with the same parameters *a_1_,a_2_* but independent realizations of driving noise *ε_1_(t),ε_2_(t)* were simulated. Importantly, the first process was fed into the second process as an additional driving term:

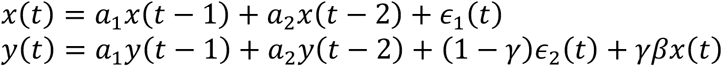

Thus, the interaction between the two processes is governed by the value of *γ.* For *γ* = 0, both cells are independent, while for γ = 1, the dynamics of the second cell is completely determined by the first one. *β* is chosen such that the variance of *x(t)* is equal to one in order to make it comparable to *ε_2_(t)*.

To analyze the power of the test, *γ* is varied between 0 and 1, and for each *γ*, 500 pairs of time series of average length and number of oscillations of the experimental data were simulated with the parameters *T* = 270 min, *τ* = 63 min (Section 16.1) for 30 data points with a sampling time of 9 min. The critical value for non-zero coherence was calculated from the quantile of 500 surrogate data using a significance level of *α* = 5%, and the fraction of cases in which the null hypothesis of zero coherence was rejected is calculated (Figure S8H).

The results show three properties of the test: (i) The significance level is correct, i.e. for *γ* = 0, 7% of the null hypotheses are rejected, which complies with the significance level *α* = 5%. Therefore, it is a valid test. (ii) A cell non-autonomous behavior would have been detected if its contribution would have been about *γ* = 0.2, and (iii) that complete non-autonomous behavior *γ* = 1 would have been detected only in about 65% of the cases. We hypothesized that the latter observation is a finite size effect. To test this hypothesis, time series of length 500 were simulated with the same parameters and the same analysis was applied. The results substantiate our hypothesis, nearly 100% of the complete non-autonomous cases are detected.

### 17 Synchronization analysis

Synchronization analysis poses an alternative to coherence analysis for inferring coupling. Synchronization between two oscillators is defined as the difference between their corresponding phases not exceeding a certain threshold, i.e.

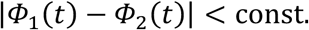

Since for noisy data, the instantaneous phase is also fluctuating, the definition of synchronization becomes more complicated and is possible in a statistical sense only. For example, rapid jumps in the phase can occur because of strong noise (Tass et al., 1998). Therefore, we suppose synchronization if the distribution of the cyclic relative phase |*Φ*_1_(*t*) − *Φ*_2_(*t*)|⁄2 *π* mod 1 shows a distinct peak (Figure S8E). For non-synchronized data on the other hand, we would expect a uniform distribution.

To measure uniformity, we use a synchronization index based on the Shannon entropy. We define 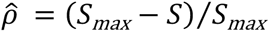 with 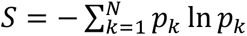, *S_max_* = ln *N* and *N* being the number of bins (Tass et al., 1998). To derive a significance threshold for non-uniform cyclic phase distributions, we apply the surrogate data approach: We generate 1000 surrogate data sets for each time series and calculate the synchronization between them. The 99% percentile then provides the threshold for the synchronization index *p̂* between the two time series.

As for the coherence analysis, synchronization analysis has to be tested for the given situation (Section 16.3). 500 datasets were simulated with the parameters *T* = 270 min, *τ* = 63 min (Section 14.1). The results show that for no coupling, *γ* = 0, in 35% of the cases, the test produces false positive (Figure S8F). Therefore, this is not a valid test. This result is substantiated by the fact that the synchronization analysis resulted in a similar number of significant cases even when cells of different embryos were analysed. To investigate whether this is a finite size effect, the analysis was repeated with tenfold longer time series. The result argues for this, because the number of false positives is reduced to 16%.

### 18 Parameter estimation in dynamical systems

The deterministic dynamics of biochemical reaction networks can be modelled by a system of ordinary differential equations (ODEs)

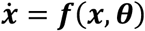

where ***f*** describes functional relationships between the concentrations of the molecular species ***x*** with the dynamical parameters and initial conditions comprised in the parameter vector ***θ***, and ***ẋ*** denotes the time-derivative of ***x***, i.e. ***ẋ*** ≡ d***x***/d*t*. For example, for three species A, B, and C, with their concentrations denoted by the symbols A, B, and C respectively, that interact via the simple conversion law shown below, the change in C can be specified as

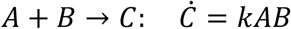

for the species vector ***x*** = (*A,B,C*)^T^. Other frequently used functional relationships include Michaelis-Menten and Hill kinetics. Based on these, arbitrary reaction networks can be translated into differential equations. Since the molecular species involved in such a reaction network can usually be observed only partially or not on the absolute scale, and experimental noise ε enters the measurements, it is necessary to introduce an observation function ***g*** that links observations ***y*** ∈ ℝ^n^ to the underlying dynamics of the model states ***x***:

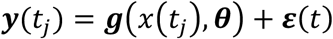

In addition to the parameters mentioned above, ***θ*** may also include observation-specific parameters such as scaling and offset. Experimental errors are usually assumed to be normally distributed, which allows defining the scaled negative log-likelihood

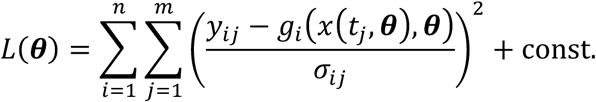

In this equation, *y_ij_* represents the measured data of the *i*-th observable at time point *t_l_*, while *g_i_*(***x***(*tj*), ***θ***) denotes the corresponding model prediction for the parameter vector ***θ***. To estimate the parameters, we use the maximum likelihood method, i.e. *L*(***θ***) is minimized to acquire an estimator for the true parameter vector:

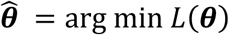

#### 18.1 Parameter profile likelihood

Assessing confidence intervals (CIs) for the estimated parameters 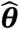 is a critical instance of parameter estimation. A widely used approach is obtaining CIs from the Fisher information matrix (Press et al., 2007), which relies on an approximation of the log-likelihood through a quadratic function. When this approximation does not hold, CIs calculated from the Fisher information matrix are not appropriate (Joshi et al., 2006). Instead, profile likelihood-based CIs are used frequently. The profile likelihood PL of the parameter of interest *θi* is calculated by scanning the corresponding parameter axis and re-optimizing the likelihood with respect to the remaining parameters at each step:

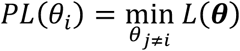

Thereby, an optimal path through parameter space is found which provides global information of the scaled negative log-likelihood instead of the local information posed by the Fisher information matrix (Raue et al., 2009). CIs can be derived from a threshold value that is defined by the fact that the test statistic

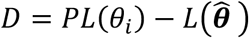

is 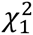 distributed, as it is a likelihood ratio test between models that differ in their degrees of freedom by one.

Depending on the shape of the profile likelihood, different scenarios concerning parameter identifiability, i.e. confinedness of CIs, can be distinguished (Raue et al., 2009):

- The profile likelihood of an identifiable parameter exceeds the threshold in both directions of the parameter axis resulting in a finite CI.
- A structurally non-identifiable parameter results from over-parametrization and leads to a constant profile likelihood with respect to the parameter of interest. It is caused by symmetries in the system that can be detected using Lie group theory (Merkt et al., 2015). To remove a structural non-identifiability, the corresponding parameters are set to fixed values which does constrain the solution space.
- So-called practical non-identifiabilities cover the case of a profile likelihood that is neither constant nor confined in both directions of the parameter axis. Increasing the amount of data or reducing model complexity by reparametrizing the system with a smaller set of parameters can eliminate a practical non-identifiability. We use the strategies of model reduction to resolve the practical non-identifiabilities (Maiwald et al., 2016).

#### 18.2 Model establishment

The biological processes of transcription and translation lead to a time delay that, together with a negative feedback, enables oscillatory dynamics. The feedback is supposed to be described by the Hill kinetics term

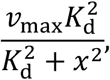

where *x* denotes the concentration of HER6 protein. We reparametrize this term by *K_d_^2^* → *mRNA_inh_* and *v_max_* → *mRNA_prod_/K_d_^2^,* which leads to the following system of ODEs

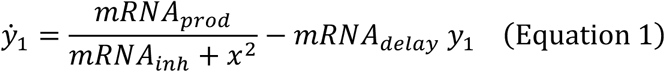

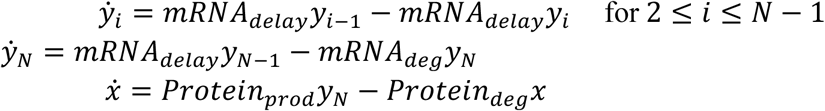

The production of HER6 is regulated by the abundance of mRNA via the production rate constant *Protein_prod_*. HER6 is degraded with a rate constant *Protein_deg_* and at the same time inhibits the transcription of nuclear mRNA. The biological processes of transcription, translation, and intercellular transport or diffusion each contribute to a time delay that, together with the negative feedback, enables oscillatory dynamics. The time is modelled by the linear chain trick (MacDonald, 1976) using the additional states *y_1_,y_2_,…,y_N_*. This corresponds to a effective delay time of *τ_delay_* = *N*/*mRNA_delay_*. Considering that the conversions within the linear chain take place with the rate constant *mRNA_delay_*, this adds up to a total of six dynamical parameters in the model. Since we estimate the noise parameter as well as the initial conditions for the *N*+1 internal model states from data simultaneously to the dynamical parameters, the model has *N*+8 parameters in total.

We use an iterative procedure to determine the optimal number of linear chain states (Hauber et al., 2019): When fitting a model including an auxiliary rate constant parameter in the linear chain that includes exactly one too many states, the auxiliary parameter becomes non-identifiable. Therefore, by scanning through the number of states, one can find the optimal number of states. This procedure was applied to all datasets individually to estimate the optimal chain length (Figure S12C). For 76 out of 104 datasets, the method suggests an optimal chain length between two and nine with larger chain lengths not being investigated. The remaining datasets result in practical non-identifiabilities that might be a consequence of even larger true chain lengths and were not investigated any further.

To investigate possible model reductions, we use the data set e07c001 (Figure 5B, left panel). We employ a multi-start approach for optimizing the scaled negative log-likelihood L(θ), because the likelihood might have multiple local optima. Initial guesses for the parameter vector are sampled from reasonable regions of parameter space. Optimization is performed numerically using lsqnonlin implemented in MATLAB R2019a. The ODEs are integrated with CVODES. Parameters are log-transformed which enforces positivity. All modelling and parameter estimation tasks are performed within the freely available Data2Dynamics toolbox for MATLAB (Raue et al., 2015; Steiert et al., 2019).

The profile likelihood reveals a practical non-identifiability of the offset parameter and a structural non-identifiability of the scaling parameter (Figure S10A). We therefore remove the offset and scaling from the observation function, i.e. setting *offset* = 0 and *scaling* = 1. Additionally, we observe a practical non-identifiability of *mRNA_inh_*, indicating that the dissociation constant is compatible with zero. Therefore, the value of this parameter is fixed to zero which reduces the Hill kinetics to a simple power law term.

After re-optimizing, we find that only the reaction flux can be unambiguously inferred from the data, but not the individual quantities determining the flux. This is because we measured protein but no mRNA concentrations. A large flux can be interpreted either as the result of a large protein production rate constant or a large overall mRNA concentration scale. Since the latter is determined by *mRNA_prod_*, the pair of parameters *mRNA_prod_* and *Protein_prod_* is linearly dependent in log-space and structurally non-identifiable (Figure S10B). To remove this over-parametrization of the model, we fix *Protein_prod_* and hence also *mRNA_prod_*. While the latter remains a non-fixed, fitted parameter, it has no biological meaning anymore since its value is determined by the arbitrary choice of *Protein_prod_*. All of the dynamical parameters of the resulting reduced model are identifiable (Figure S10D).

The reduced oscillation model with the fixed parameters *scaling*, *offset*, *mRNA*_p*rod*_ and *mRNA_inh_* was fitted to all 76 datasets that are considered oscillatory in terms of the classification procedure (Figure 3) and could be assigned an optimal chain length as described above. We find that the reduced oscillation model poses a valid simplification for all of the investigated datasets in terms of a likelihood ratio test.

#### 18.3 Stability analysis

The most important dynamical parameters exhibit uniformness within each cell. *Protein_deg_* and *mRNA_deg_* are close to being identical, while *mRNA_delay_* shows a weaker link to the degradation rate constants (Figure 5C). The overall scatter of all three parameters is roughly one order of magnitude which resembles the usual biological variability.

The fact that equal degradation rate constants facilitate oscillations was already reported in literature (Page and Perez-Carrasco, 2018) and can be reasoned by linear stability analysis of the fixed points of the system (Equation 1, section 18.2). If we denote ***z*** = (*y*_1_, *y*_2_,…, *y_N_*, *x*)*^T^* and the right-hand side of Equation 1 as ***f***(***z***), the fixed point ***z***^∗^ is defined as ***f***(***z***) = 0. The Jacobian evaluated at ***z***^∗^ is

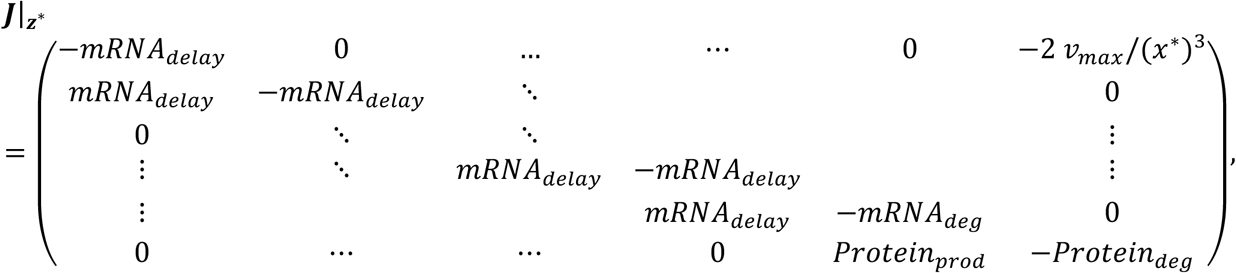

and the characteristic equation is given by

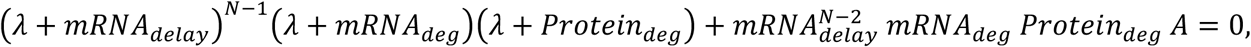

where *A* ≡ 2*v_max_ Protein_prod_* / *mRNA_deg_ Protein_deg_* (*x*^∗^)^3^ > 0. All rate constants can be assumed to be nonzero and positive because else there would be no dynamics in the system. Therefore, one can derive the inequality

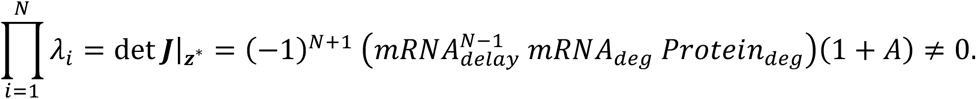

Because none of the eigenvalues can be zero, a change in stability of the system can only occur via a Hopf bifurcation. For *A = 0*, the eigenvalues are equal to minus the respective degradation rate constant and the fix point is stable. A change in stability occurs at the bifurcation where *A = Ã* and a pair of eigenvalues can be written as ±i*α*. Continuing from here, it can be shown that the critical value *Ã*, and therefore the restrictions on the whole system for exhibiting oscillations, is minimized when the degradation and delay rate constants coincide (Page and Perez-Carrasco, 2018). For the system (Equation 1), this corresponds to *mRNA_delay_ = mRNA_deg_ = Protein_deg_* (Figure 5E). For longer chains, the condition becomes more relaxed and larger values of the degradation rate constant have only a small effect on *Ã*. However, its minimum remains at equal degradation and delay rate constants. The delay rate constants show a larger spread because they must be tuned to match the time delay present in the data.

For the sake of simplicity, we show the uniformity of degradation rate constants for *N = 2*. Solving the characteristic equation at the bifurcation leads to

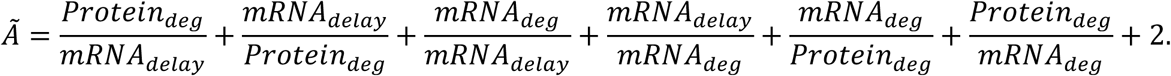

*Ã* is minimized for equal protein and mRNA degradation and delay rate constants. The results of parameter estimation show that uniformness of degradation and delay rate constants holds for the majority of analyzed time series (Figure 5C).

We find that while oscillations still occur even if the degradation rate constants differ by up to roughly one magnitude, the most sustained oscillations – which are realized in the oscillatory data – are produced for equal degradation rate constants (Figure 5F). However, because the assumption of the protein or mRNA degradation being an elementary reaction introduces a level of abstraction to the model, it is important to acknowledge that this result is to be interpreted within the scope of the model structure described above only.

To further back our statement, time series data for protein (Her6-mNG; embryo 21; embryo 33 was not used for decay rate calculation because the brief time after heat shock did not allow for determination of time window with pure Her6-mNG decay) and mRNA (*her6-mNeonGreen* mRNA and endogenous *her6* mRNA) were extracted from RT-PCR data from 4 embryos after her6 heat shock overexpression. The data were fitted with the following decay model

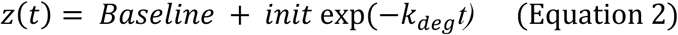

(Fig 5 H-J), where *k_deg_* denotes either *mRNA_deg_* or *Protein_deg_*. Data from different embryos were allowed to have individual baselines as well as individual noise parameters for the protein measurements. Assessment of asymmetric confidence intervals based on the profile likelihood shows that there is no statistical significance to reject the null hypothesis of equal degradation rate constants (Figure 5H-J bottom).

### 19 Software

- ZEN (v9.2.6.54) was used to collect imaging raw data and image processing
- Huygens Professional (v18.1) was used for deconvolution of imaging raw data
- Zeiss AxioVision SE64, Rel.4.9.1 was used for imaging of alkaline phosphatase-based in situ hybridizations
- Python 3.7 was used for data pre-processing (such as sorting, averaging, calculating standard deviation) together with the software library’s pandas and numpy for python. For data visualization the library matplotlib for python was used.
- MATLAB R2019b was used for data analysis together with the open-source toolboxes Data2Dynamics (github.com/Data2Dynamics) for dynamical modeling and parameter estimation, as well as tsFramework (Software S1) for time series analysis.
- Microsoft Excel 2016
- FIJI (imagej.net/Fiji)
- GPML v4.2 (www.gaussianprocess.org/gpml/code/matlab/doc/index.html) and GPosc (https://github.com/ManchesterBioinference/GPosc)
- (Clustered) Heatmaps were generated using Heatmapper http://heatmapper.ca (Babicki et al., 2016)
- Arivis Vision4D (v2.12.6, v3.0.1, v3.01, v3.1.2 and v3.1.3) was used for measurements and data export

## Resource availability

Further information and requests for resources and reagents should be directed to Wolfgang Driever.

## Ethics Oversight

All animal experiments were approved by state authorities (Regierungspraesidium Freiburg permits 35-9185/G-16/123, 35-9185.81/G-19/19 and 35-9185.81/G-19/54).

## Acknowledgements

We thank Sylke Lange for technical assistance, Angela Naumann and the Life Imaging Center (LIC) for support in SPIM and confocal imaging and image analysis, Wolf Heusermann and Oliver Biehlmaier at the Imaging Core Facility at Biozentrum Basel for advice on SPIM imaging, and Sabine Götter for excellent fish care.

## Funding

This work was funded by grants from the Deutsche Forschungsgemeinschaft (DFG, German Research Foundation) under Germanýs Excellence Strategy – EXC-2189 CIBSS– Project ID: 390939984 (Gefördert durch die Deutsche Forschungsgemeinschaft (DFG) im Rahmen der Exzellenzstrategie des Bundes und der Länder – EXC-2189 – Projektnummer 390939984), the Excellence Initiative of the German Federal and State Governments (BIOSS - EXC 294), and CRC 850. The authors acknowledge support by the Deutsche Krebshilfe (Grant no. 70112355), the state of Baden-Württemberg through bwHPC, and the German Research Foundation (DFG) through grant INST 35/1134-1 FUGG.

## Author contributions

W.D. conceived the study. C.S., J.R., S.B. and D.S. designed and performed experiments. C.S. performed cloning and generated transgenic lines. C.S., A.H. and S.B. analyzed data. J.T. and A.H. performed analyses of oscillations and coupling, and developed the ODE model. R.N. contributed to confocal and SPIM microscopy and image analysis pipeline. W.D., J.T., C.S. A.H. and J. R. wrote the manuscript.

## Declaration of interests

The authors declare no competing interests

## Supplementary Information

Supplementary Figures S1-S15 are appended to this PDF. Additional supplementary data will be made available upon peer review journal publication.

## Notes

### Competing Interest Statement

The authors have declared no competing interest.

### Summary of Updates

We have edited Abstract, Introduction, Results and Discussion for clarity. DOI links have been added to references.

