## Supplementary Figures S1 - S15 for "Parameters for stable Notch-independent Her6 oscillations across a frequency spectrum in neural stem cell populations"

GERMANY

Jens Timmer

Institute of Physics

University of Freiburg

Hermann - Herder Str. 3

D-79104 Freiburg

GERMANY

ORCID IDs: Sigloch (0000-0003-3226-7132), Hauber (0000-0002-3053-5124), Böhm (0009-0000-8004-7818), Nitschke (0000-0002-9397-8475), Timmer (0000-0003-4517-1383), Rothenpieler (0000-0001-8892-8230), Driever (0000-0002-9551-9141)

**Figure S1: *her6* expression in the zona limitans intrathalamica (ZLI), neural proliferation**

**zones, and general ventricular layer neural stem cells**

**(A, B)** Hybridization chain reaction (HCR) staining for *her6*, *her4* and *shh* in the 3 dpf TPZ shown in dorsal (A) and resliced frontal (B) views. ZLI is marked by *shha* expression (A''', B'''). High *her6* expression is restricted to ZLI and rTh(p) and is associated with absent or low *her4* expression. In contrast, *her4* expression in ventricular regions coincides with low *her6* expression. **(C)** Midsagittal section of a 3 dpf zebrafish brain stained by HCR for *her6*, *her4* and *shha*. In the diencephalon, *her4* expression correlates with lower *her6* expression levels. **(D)** Ventricular telencephalic cells, as part of the compartment, co-express *her6* and *her4* at different levels and intermingle with each other. In the thalamus, cells are organized in alternating stripes of *her6*-high and *her6*-low/*her4* expression. **(E)** Anatomical model indicating the section planes for (A) and (B). **(F)** Anatomical scheme of the thalamus highlighting the differential expression domains of NI *her6* and ND *her4*. Scale bars: 20 μm if not stated otherwise.

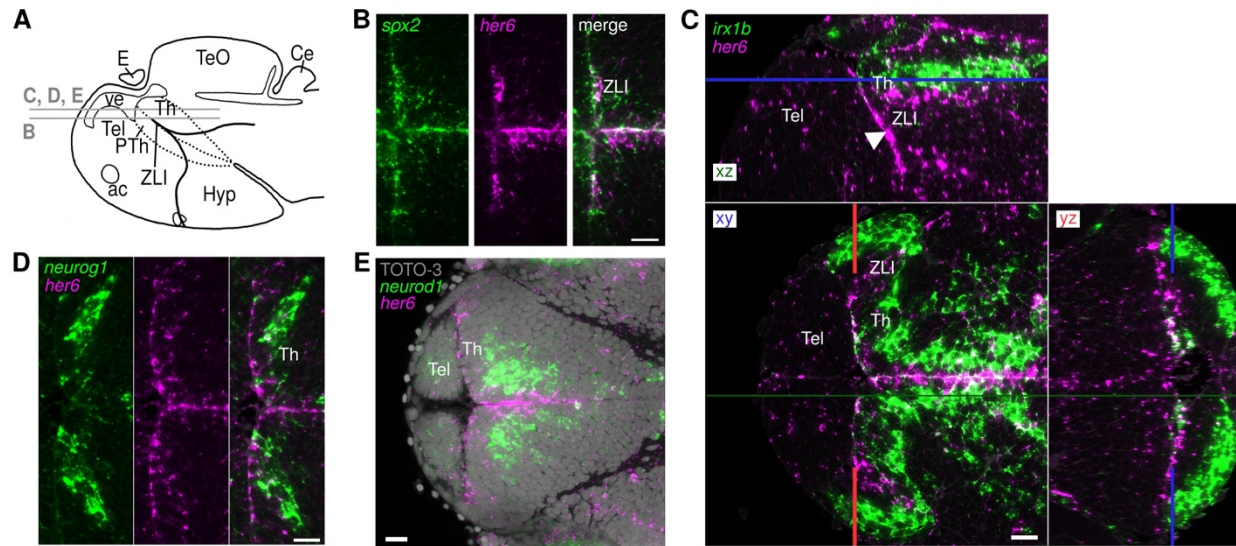

**Figure S2: *her6* expression in relation to neurogenesis and thalamic anatomical markers**

(A) Schematic representation of the forebrain, indicating imaging planes of (B) – (E). (B - E) Double-fluorescent in situ hybridization of (B) *her6* and *sox2* in 56 hpf wildtype embryos, with strong colocalization along the ventricle, albeit with differences in expression strength across diencephalic domains. (C) *her6* and *irx1b* in 56 hpf embryos. xy, image plane through the thalamus. xz, lateral view parasagittal plane. yz, frontal view. *irx1b* is expressed in the thalamus but not in the ZLI (arrowhead). (D) *her6* and *neurog1* in the 56 hpf thalamus, expressing *neurog1* mostly in the subventricular cell layers. (E) *her6* and *neurod1* in the 56 hpf thalamus with an additional nuclear TOTO-3 stain, revealing the distance of *neurod1* expression to the ventricular surface NSCs by 1-2 cell layers.

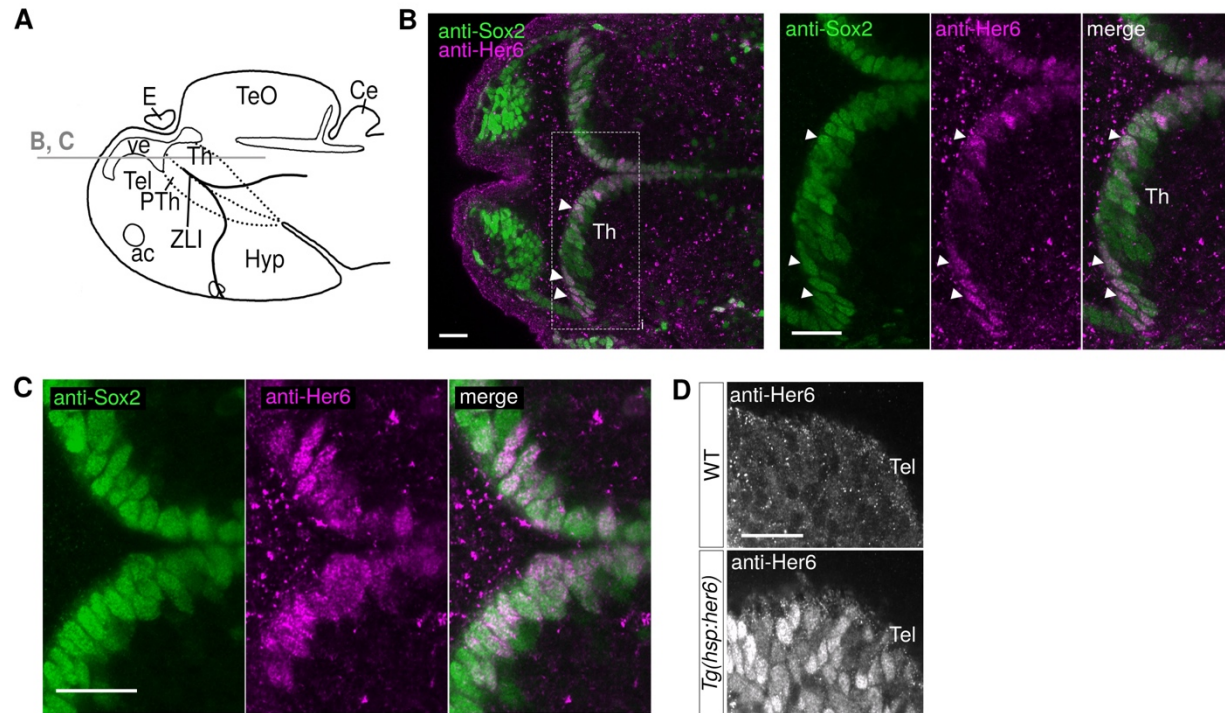

**Figure S3: Endogenous Her6 protein is detected at varying expression levels in nuclei of Sox2+ ventricular neural stem cells**

(A) Schematic representation of the forebrain, indicating imaging planes of (B) and (C). (B) Immunofluorescence staining of the stem cell marker Sox2 and NI Her6, revealing broad expression along the ventricular surface in the TPZ. White arrowheads show Her6<sup>high</sup> and Sox2 co-expressing cells of the ventricular wall. The dashed rectangle is enlarged in the right panel. (C) Protein levels of Sox2 and Her6 seem to be unlinked, with Her6 being much more dynamic either through oscillatory expression or total expression levels which differ between the diencephalic anatomical domains. (D) Immunofluorescent validation of the Her6-antibody as well as the overexpression capabilities of the *Tg(hsp70l:her6-FLAG)<sup>m1492</sup>* line in the 56 hpf telencephalon one hour after heat shock.

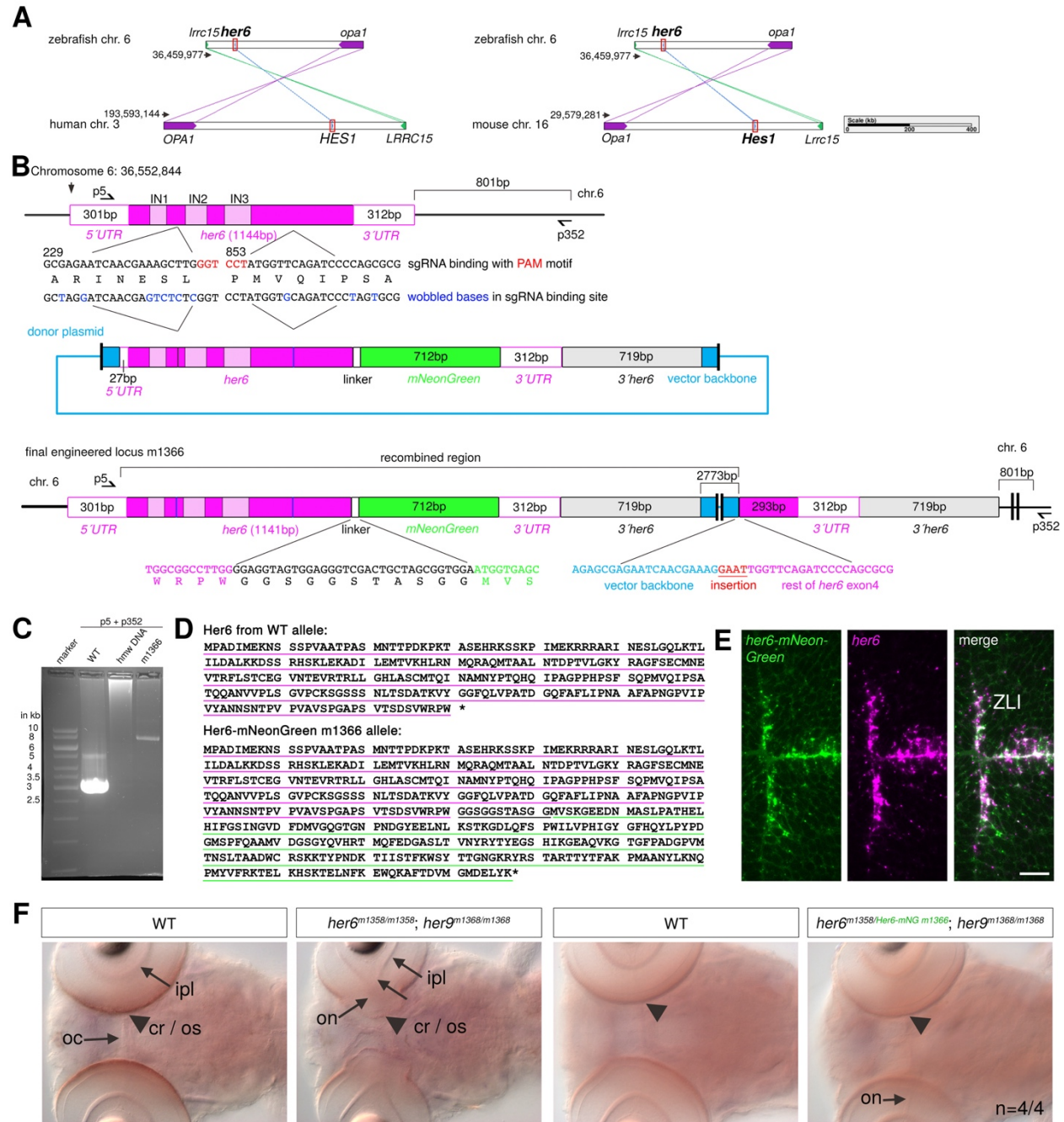

**Figure S4: Schematic of Her6-mNeongreen knock-in and validation of protein functionality**

(A) Zebrafish *her6* is located within the same synteny block as human *HES1* (left) and mouse *Hes1* (right). Base pair numbers indicate the locations on the chromosomes.

(B) Schematic representation of the *her6-mNeonGreen m1366* knock-in locus after integration of *mNeonGreen* at the C terminus of Her6 by homologous recombination. Schematic depiction of the *her6* locus, donor plasmid, and final engineered locus. The positions and sequences of the sgRNA target sites are shown with the PAM sequences in red. The donor plasmid *her6:her6-*

*linker-mNeogreen-3'UTR[her6]* contains both sgRNA target sites, but with several nucleotides exchanged (blue). The amino acid sequence remains unchanged (capital letters). A 33 bp linker was fused directly in front of the stop codon of *her6*. The *mNeonGreen* sequence is followed by the sequence of the endogenous 3'UTR and 719 nucleotides downstream genomic sequence of the *her6* locus. The final engineered locus contains the rest of the vector backbone followed by the rest of exon4 and the 3'UTR. Half arrows indicate primer binding sites for p5 and p352. The resulting PCR is shown in C. IN, introns; bp, base pairs.

(C) PCR that amplifies the whole locus. Both primers only bind on the chromosome but not on the Donor Plasmid. Expected size for the PCR product from WT control DNA: 3097 bp. Expected size for the *her6-mNeonGreen* allele m1366: 7955 bp. hmw DNA, unspecific high molecular weight DNA.

(D) Amino acid sequence of the Her6-mNeonGreen fusion protein in the m1366 allele.

(E) Double fluorescent in situ hybridization of *mNeonGreen* and *her6* in a 56 hpf homozygous *Tg(her6:her6-mNeonGreen)<sup>m1366/m1366</sup>* embryo. Dorsal view confocal optical section of the telencephalon and ZLI. Scale bar: 20 μm.

(F) Validation of biological activity of Her6-mNeonGreen fusion protein in *Tg(her6:her6-mNeonGreen)<sup>m1366</sup>*. Dorsal views of 4 dpf fixed embryos, DIC optical section. Wild type (WT) control embryos are shown in first and third panels for mutant fish in the second and fourth panel respectively. *her6<sup>m1358/m1358</sup>*; *her9<sup>m1368/m1368</sup>* double mutant with malformations at the central retina and optic stalk (arrowhead). ipl, inner plexiform layer; oc, optic commissure; on, optic nerve. The phenotype has high penetrance and is observed in all double mutant embryos. A single *Tg(her6:her6-mNeonGreen)<sup>m1366</sup>* allele rescues the malformations of the *her6/her9* double mutant phenotype at the central retina and optic stalk.

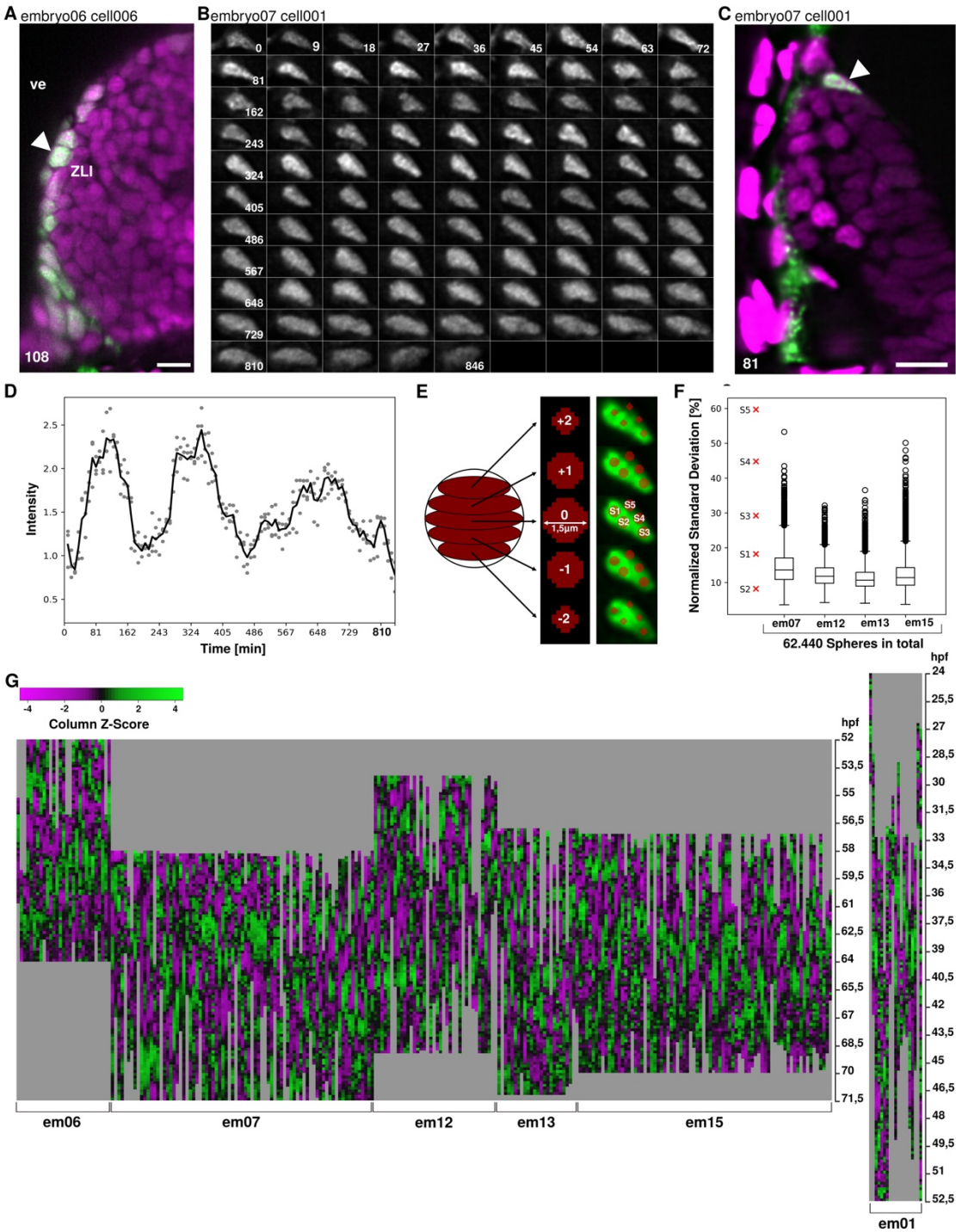

**Figure S5: Her6-mNeonGreen measurements in time series data**

(A) Location of the cell in the ZLI shown in main Figure 2A at the ZLI ventricular wall (green, arrowhead). SPIM image, dorsal view, NLS-mCardinal in purple. Scale bar, 10 μm.

(B) Time series SPIM imaging of a different Her6-mNeongreen expressing cell starting at 56

hpf. Numbers indicate minutes after start of recording. Scale bar, 2 $\mu\text{m}$.

(C) Location of the oscillating cell shown in **B**, SPIM image, scale bar 10 $\mu\text{m}$.

(D) Quantification of the Her6-mNeonGreen intensity of the cell shown in **B**. Graph shows individual measurements of all three spheres (small circles) for each time point, and mean intensity (black line).

(E) Measurement strategy. Depiction of individual layers from a sphere as individual circles (middle panel). The right panel shows five consecutive SPIM image planes of the cell shown in **B**. S1-S3 are well positioned spheres, while S4 and S5 are spheres that extend out of the nucleus and are considered poorly placed.

(F) Quality control of all sphere measurements visualized for four embryos. Each sphere consists of 228 quantified voxels. From these 228 data points, normalized standard deviation (sd) was calculated. Left lane (red x) shows normalized sd values for well positioned spheres S1, S2 and S3, as well as for misplaced spheres S4 and S5 (see **E**). For each embryo, normalized sds for each sphere were plotted. Boxes indicate the 68% interval and bars indicate the 96% interval. Spheres with a normalized sd larger than the one observed for S4 were considered potentially misplaced, and for these cells the sphere was positioned and measured again.

(G) Heatmap representation of Her6-mNeonGreen fluorescence for all measured cells aligned by real developmental time. Each vertical column represents one cell. Grey shaded areas indicate cells were not tracked. Left panel 2 dpf embryos as indicated at bottom, real developmental time indicated at right. Right panel same for 1 dpf embryo. Color code shows z-scores on data scaled on each cell as indicated. Data were not clustered.

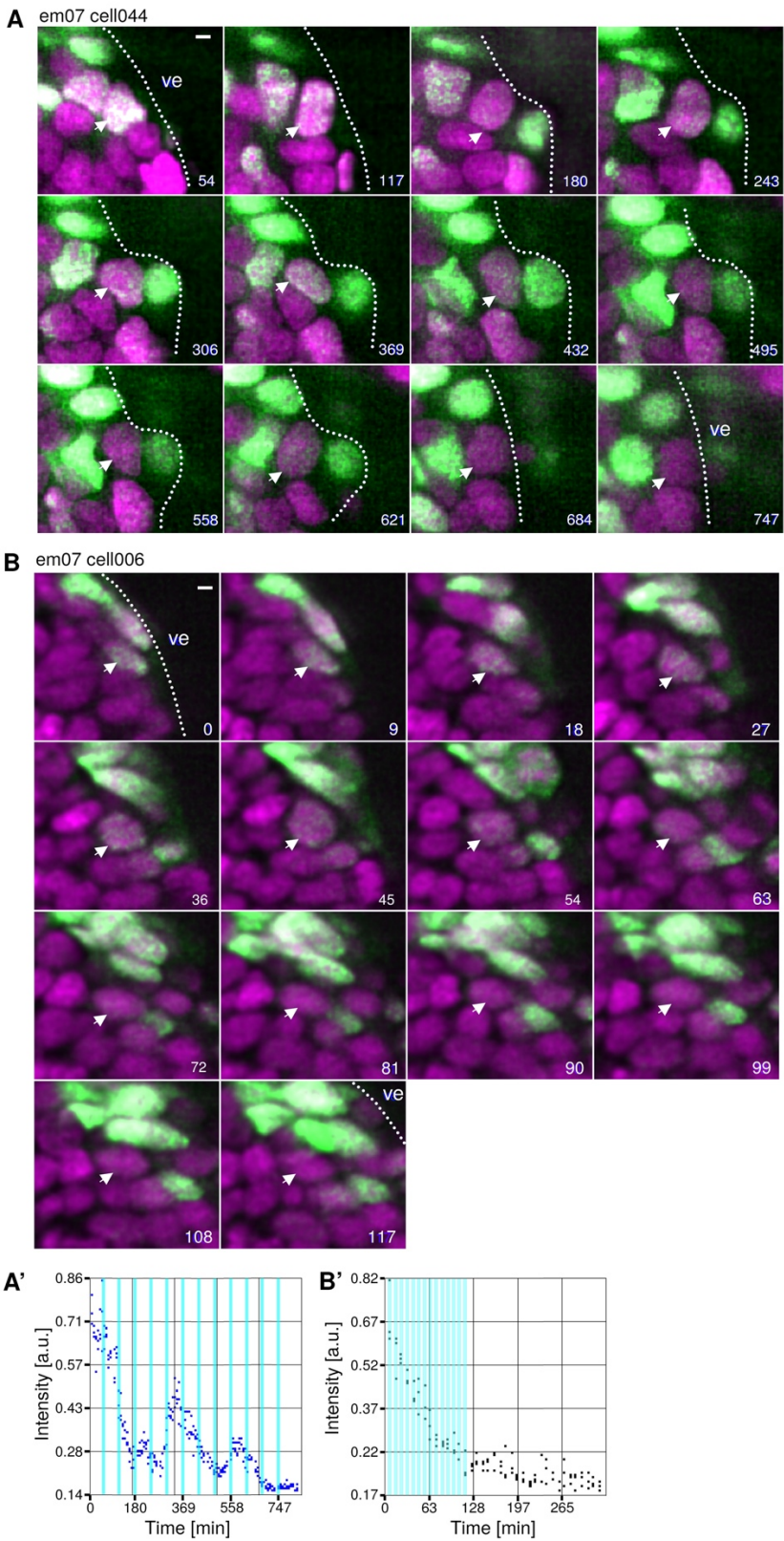

**Figure S6: Cell behavior during phasing out of Her6 expression.**

**(A)** Oscillation and Her6-mNeonGreen expression cease while cell remains at the ventricle. Cell 044 from embryo 07 remains at the thalamic ventricular surface and turns off Her6-mNeongreen expression. The SPIM image series starts at T=7 (58 hpf + 54 min). Numbers in bottom right corners represent time (min) since start of time series. Ve, ventricle; dotted line, ventricle surface.

**(B)** Downregulation of Her6-mNeonGreen concomitant with migration away from the ventricle. Cell 006 from embryo 07 migrating away from the thalamic ventricular surface turns off Her6-mNeongreen expression (arrow). The SPIM image series starts at T=1 (58 hpf). Numbers in bottom right corners represent time (min) since start of time series. Scalebars for A and B, 2 μm; ve, ventricle; dotted line, ventricle surface.

**(A', B')** Quantification of Her6-mNeongreen fluorescence intensity of cells in **A** and **B**. Cyan bars indicate the frames shown in **A**, **B**. In each nucleus, three measuring spheres were placed per time point. Each dot represents the measured intensity of one sphere (see Figure 2A and Figure S5E).

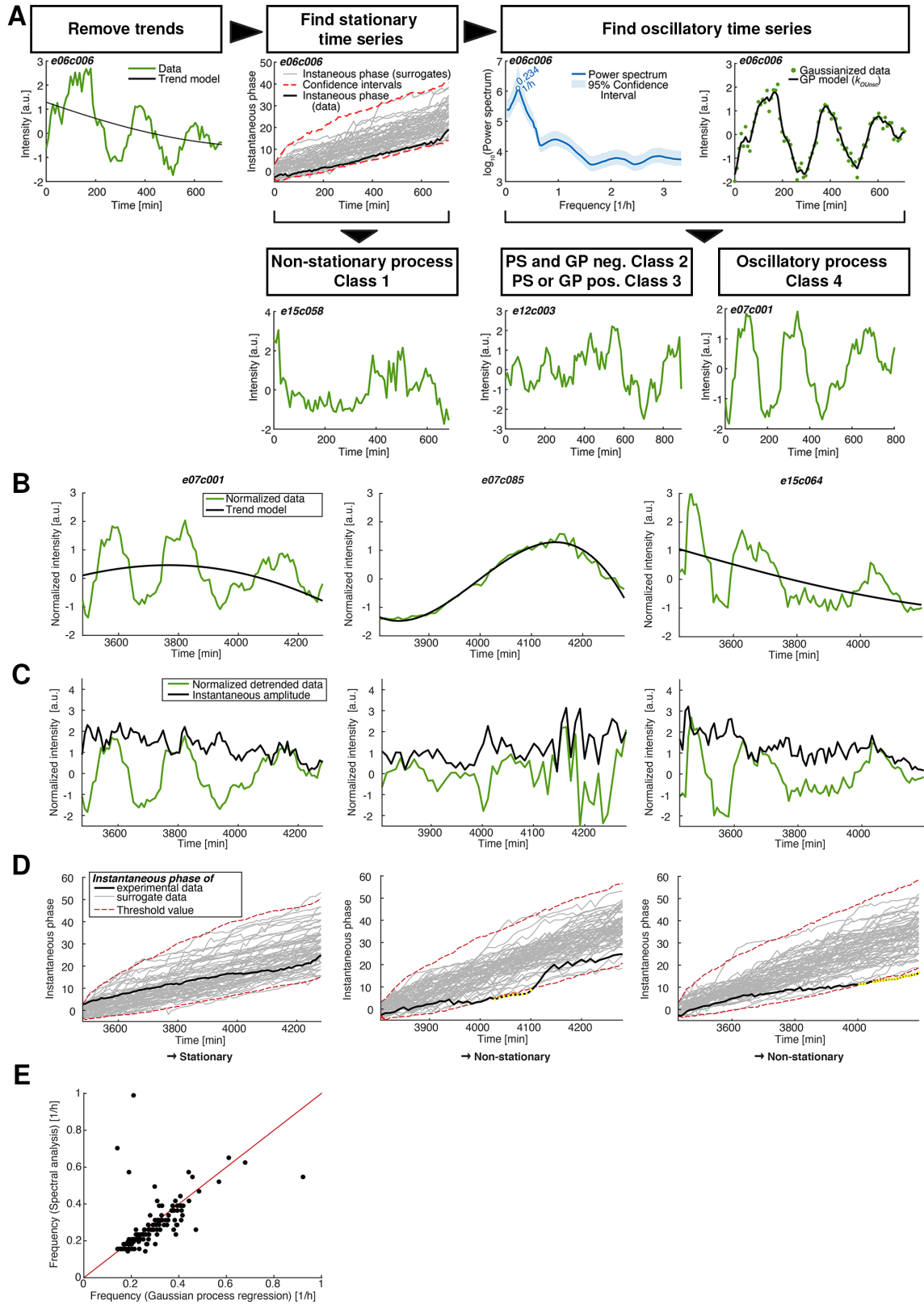

Figure S7: Analysis of Her6-mNeonGreen expression time series data

**(A)** Schematic representation of data classification pipeline. Normalized detrended data (Methods 11.1) is filtered for stationary processes (Methods 15). The power spectrum of the remaining time series is calculated (Methods 16.1) and a model selection approach using Gaussian processes is employed (Methods 11.2) to determine whether the underlying process is of first order (not able to oscillate) or second order (potentially oscillatory).

**(B)** Normalized data (green) with squared exponential trend model (black) calculated by Gaussian process regression. To ensure the trend does not depict dynamics of interest, the parameter governing the characteristic length scale of the process is restricted by a lower bound during optimization. Three cells (cell numbers indicated at top of each panel) with distinct characteristics of temporal profiles have been selected, which can be considered as typical for a large portion of cells analysed.

**(C)** The normalized detrended data (green), i.e. normalized data subtracted by the trend model in **(B)**, and the instantaneous amplitude (black).

**(D)** In addition to the instantaneous amplitude, the instantaneous phase (black) can be calculated from the detrended normalized data in **(C)**. A surrogate data approach is employed to discriminate between stationary and non-stationary time series: For each time point, 0.5% and 99.5% percentiles of the instantaneous phases of 3000 surrogate data (grey, only 60 are shown for better visibility) are used to derive threshold values (red) for the instantaneous phase of the experimental data. If the latter crosses one of the thresholds, the time series is regarded non-stationary.

**(E)** Comparison of the dominating frequencies in the data calculated from Gaussian process regression ($x$-axis) and spectral analysis ($y$-axis). The scattering around a line with unit slope (red) indicates consistent results.

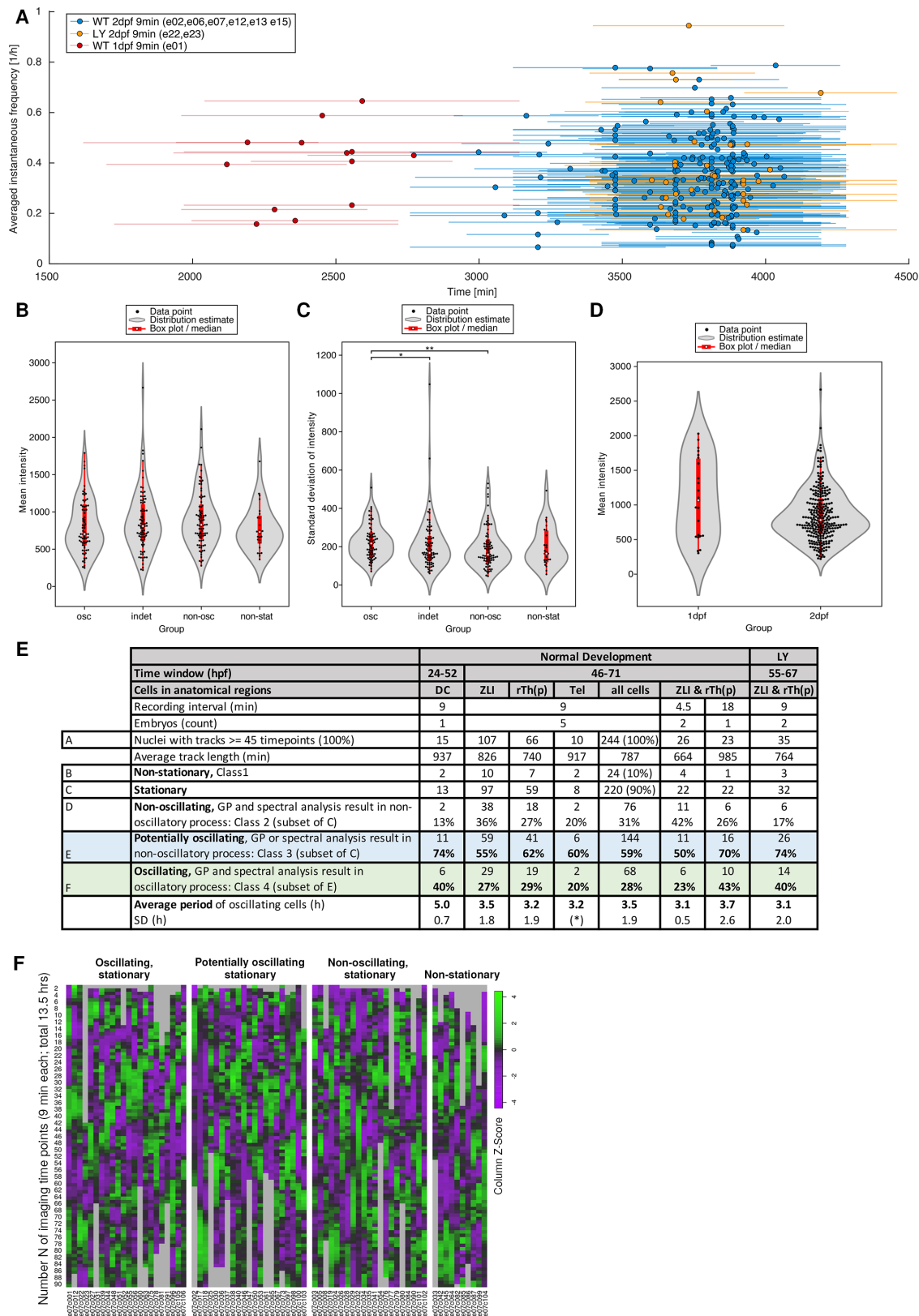

**Figure S8: Her6 oscillation frequencies in wildtype and experimental embryos**

**(A)** Averaged instantaneous frequency per cell (filled dots) versus the respective measurement time window (fine line) in minutes post fertilization (1 through 3 dpf). Data are derived from embryo numbers as indicated in box.

**(B)** Estimated distribution of mean fluorescence intensity for individual cells classified as oscillating (osc, class 4), indeterminate (indet, class 3 = potentially oscillating), non-oscillating (non-osc, class 2), and non-stationary (non-stat, class 1) during third (2 dpf) day of development according to Supplementary Figure 7A.

**(C)** Estimated distribution of standard deviation of fluorescence intensity tracks as proxy for amplitudes for individual cells classified as oscillating (osc, class 4), indeterminate (indet, class 3 = potentially oscillating), non-oscillating (non-osc, class 2), and non-stationary (non-stat, class 1) during third (2 dpf) day of development according to Supplementary Figure 7A. \* $p < 0.05$, \*\* $p < 0.01$.

**(D)** Estimated distribution of mean fluorescence intensity for individual cells during the second (1 dpf, left) and third (2 dpf, right) day of development.

**(E)** Summary of tracking data and analysis of all cells (note: “all cells” also contains 61 cells that could not be assigned to one of the three anatomical regions). Classification of cells according to Supplementary Figure S7A is indicated in lines B to F. Light blue shading indicates potentially oscillating cells in line E, while light green shading indicates oscillating cell based on both Gauss Process and spectral analysis. LY indicates Notch inhibitor treated embryos. SD, standard deviation. (\*) both cells had same period.

**(F)** Heatmap showing detrended Her6-mNeonGreen intensity data for all embryo e07 cells with tracks long enough for mathematical analysis ($N=74$; day 3 of development; see Supplemental Data Sheet 1). Cell numbers are indicated below each column. Imaging starts at timepoint $N=1$ equivalent to 3480 min and ends at $N=90$ equivalent to 4281 min post fertilization. The cells are grouped based on the classification shown in panel E and in Supplementary Figure S7A: from left: class 4 (oscillatory, stationary), class 3 (potentially oscillating [indeterminate], stationary), class 2 (non-oscillatory, stationary), and class 1 (non-stationary). Data were scaled for each cell (column) and Z-scores are plotted using a color scheme green – high to magenta – low. Grey indicates time points at which cells were not recorded.

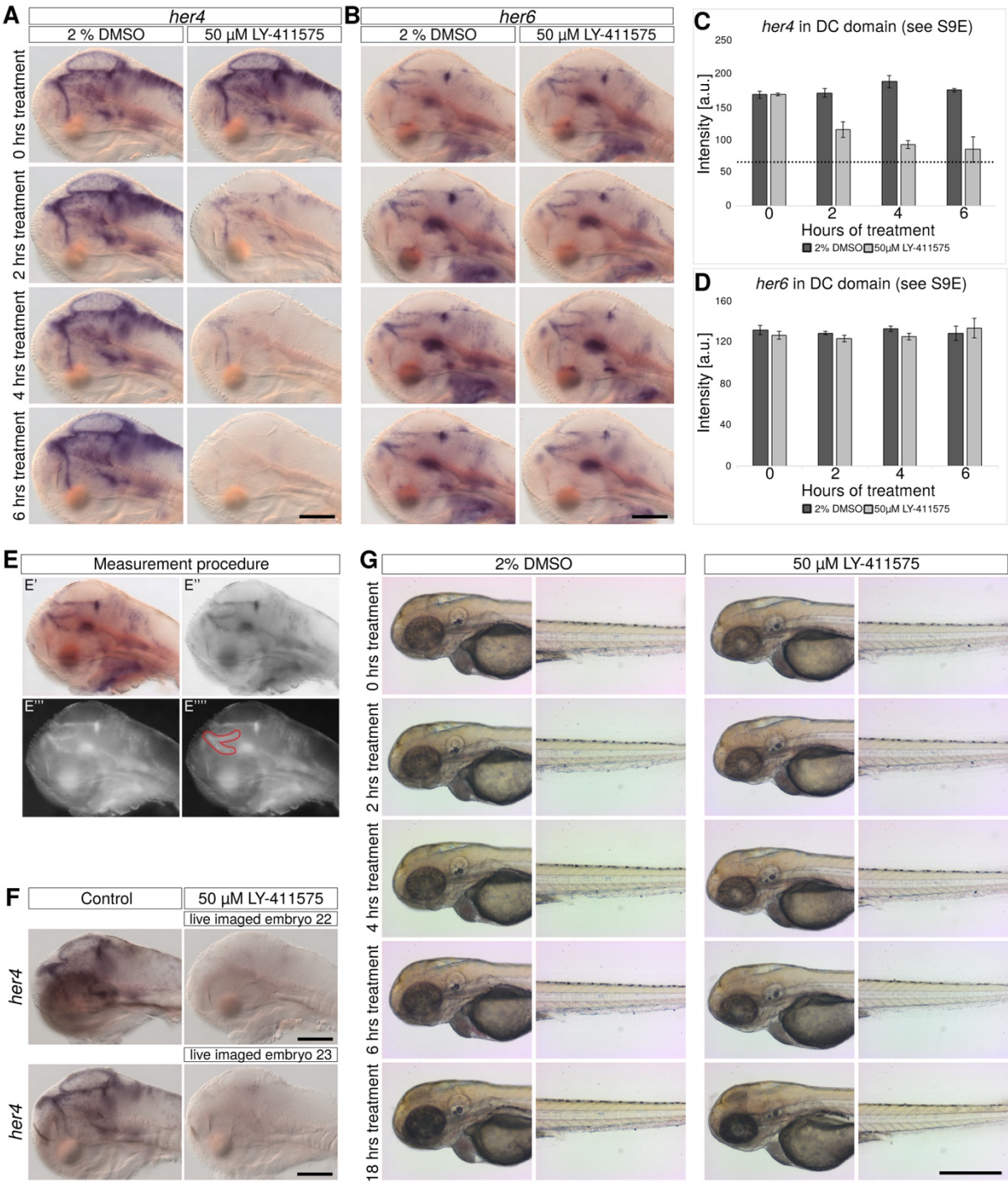

**Figure S9: LY-411575 Notch inhibition downregulates *her4* but not *her6* expression**

(A,B) In situ hybridization for *her4* (A) and *her6* (B) expression in LY-411575 (50 μM in 2% DMSO) treated embryos and in control (2% DMSO) embryos. Embryos were fixed before (0 h) and 2, 4, and 6 h after onset of treatment. Scale bars, 200 μm.

(C,D) Quantification of *her4* and *her6* in situ hybridization signals from experiments shown in A

and **B**, respectively. For each data point, stain intensities of four different embryos were measured in an area of the diencephalon (DC) indicated in **E**. The dashed line in **C** indicates the mean image intensity (full area of the picture) of the 6 h treated LY-411575 image from **A** to indicate background intensity values in the absence of specific expression.

**(E)** Method of intensity measurement shown in **C** and **D**, area of measurement indicated in red (see Methods).

**(F)** Validation of Notch inhibition in experimental embryos recorded in the SPIM. *her4* in situ hybridization of previously live imaged embryos 22 and 23. Embryos were recovered from the SPIM after completion of imaging, fixed and stained for Notch-dependent *her4* expression to validate successful LY-411575 treatment. Control embryos were kept in 2% DMSO and were also embedded in low melting agarose. Scale bar, 200 μm.

**(G)** Survival control of agarose embedded LY-411575 treated embryos. No imaging was performed with these embryos. Morphology appears normal until 6 h of treatment, but after 18 h of treatment turbid regions in the brain appear. Scale bar, 200 μm.

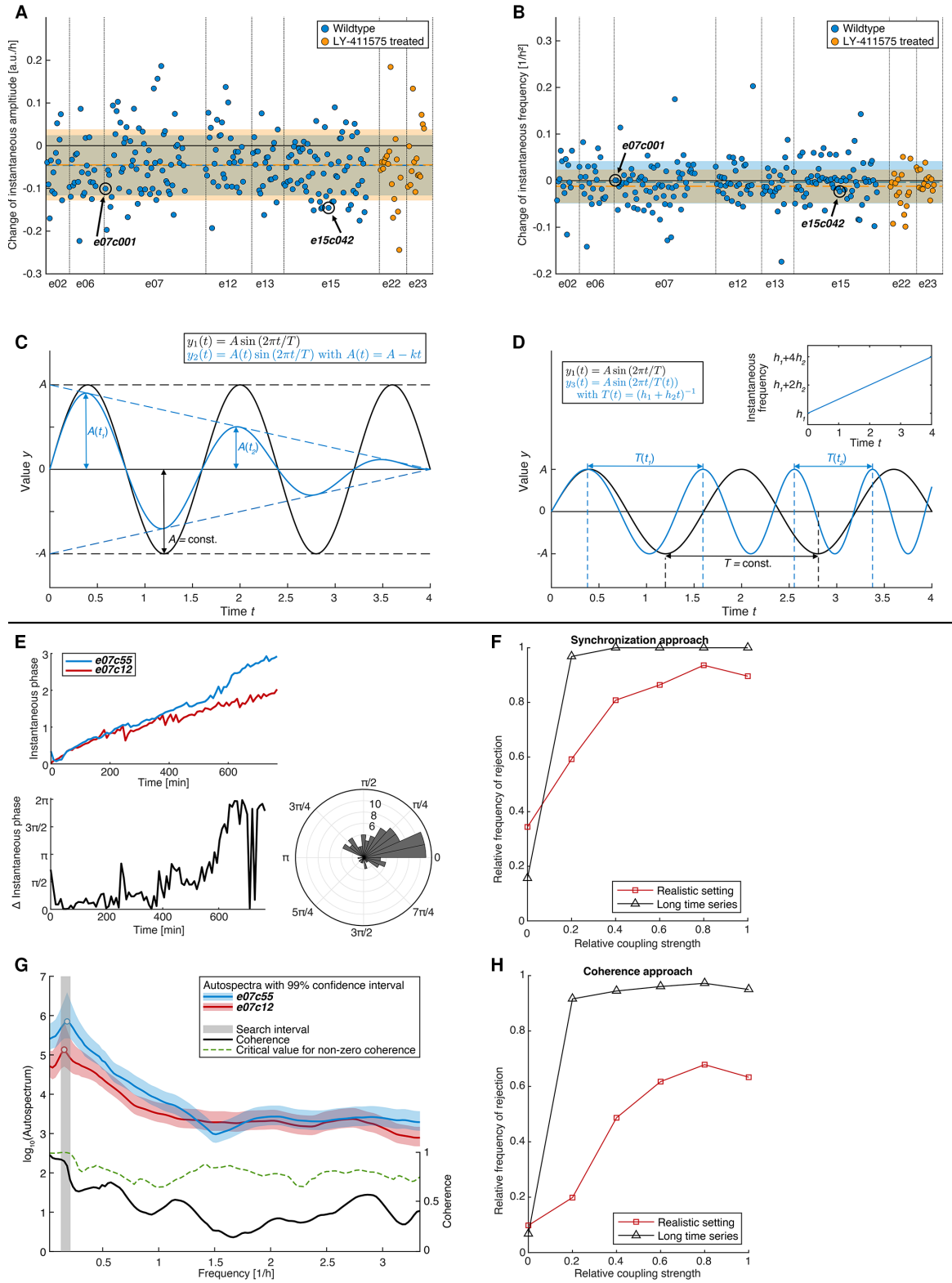

**Figure S10: Analysis of cell autonomy: (1) Instantaneous frequency and amplitude of Her6 expression are not affected by Notch inhibition, and (2) investigation of the validity and the power of the tests for coupling based on coherence and synchronization analysis**

(A) Distribution of changes of instantaneous amplitude. Each dot denotes one cell. Cells corresponding to LY-treated embryos are marked in orange. Mean and standard deviation of the two datasets are indicated by a dashed line and a shaded area respectively. The instantaneous amplitude and phase of cell e07c001 are shown in Figure S7B and C.

(B) Distribution of changes of instantaneous frequency. For details see (A).

(C) Illustration of concept of instantaneous amplitude used in (A). Plot of two functions $y_1(t)$ and $y_2(t)$, the first of them having a constant amplitude (black) and the second a linearly decreasing instantaneous amplitude (blue).

(D) Illustration of concept of instantaneous frequency used in (B). Plot of two functions $y_1(t)$ and $y_3(t)$. $y_3(t)$ (blue) has a linearly increasing instantaneous frequency (upper right panel). Since the period is the inverse of the frequency, the latter becomes shorter when going forward in time.

(E-H) Mathematical tests to detect potential coupling of Her6 oscillations between cells. A power-of-the-test simulation study that scanned from a cell-autonomous to a cell-non-autonomous situation revealed that synchronization analysis (E,F) cannot be applied under the given finite size conditions because it produced too many false positive under the null hypothesis in the experimental data. Coherence analysis showed the correct size of the test in the simulation study (G,H).

(E) Visualization of the synchronization approach for analysing coupling. The instantaneous phase of two cells are calculated (top panel) and their difference is calculated (bottom left panel). Phase differences are binned and visualized as a polar histogram (bottom right panel). For perfectly synchronized time series, the histogram would exhibit counts only for one bin, while for completely unsynchronized cells, the histogram would follow a uniform distribution. A statistical test for uniform distribution allows to infer coupling.

(F) Evaluation of the power-of-the-test of the synchronization approach for analysing coupling. Realistic datasets were simulated, for which the coupling between the two time series can be controlled, and subjected to synchronization analysis. Rejections of the null hypothesis of uncoupled time series were counted as a function of the coupling strength. This method was applied for simulated datasets of a length 80 time points that corresponds to the experimental data (red), as well as with datasets of length 1000 (black). The synchronization approach does not reproduce to correct significance level of 5% under the null hypothesis of no coupling, neither for the realistic nor the long time series setting. Therefore, it is not suitable to test for coupling because it produces too many false positives.

(G) Visualization of the coherence approach for predicting couplings. The autospectra (blue, red) of the time series of two cells are searched for significant peaks (white dots), which represent the dominating frequency in the time series. If there are significant peaks within a biologically reasonable frequency interval in the spectra of both time series, the coherence (black) - a measure for linear predictability - is calculated. Critical values for non-zero coherence are calculated from surrogate data (green dashed). If the coherence exceeds its critical value in a range (gray) around the frequencies of the peaks in the spectra, the cells corresponding to the respective time series are considered coupled.

**(H)** Evaluation of the power-of-the-test (see **F**) of the coherence approach. The coherence
approach leads to only 7% rejections under the null hypothesis and complies with the
significance level and poses a valid test.
293

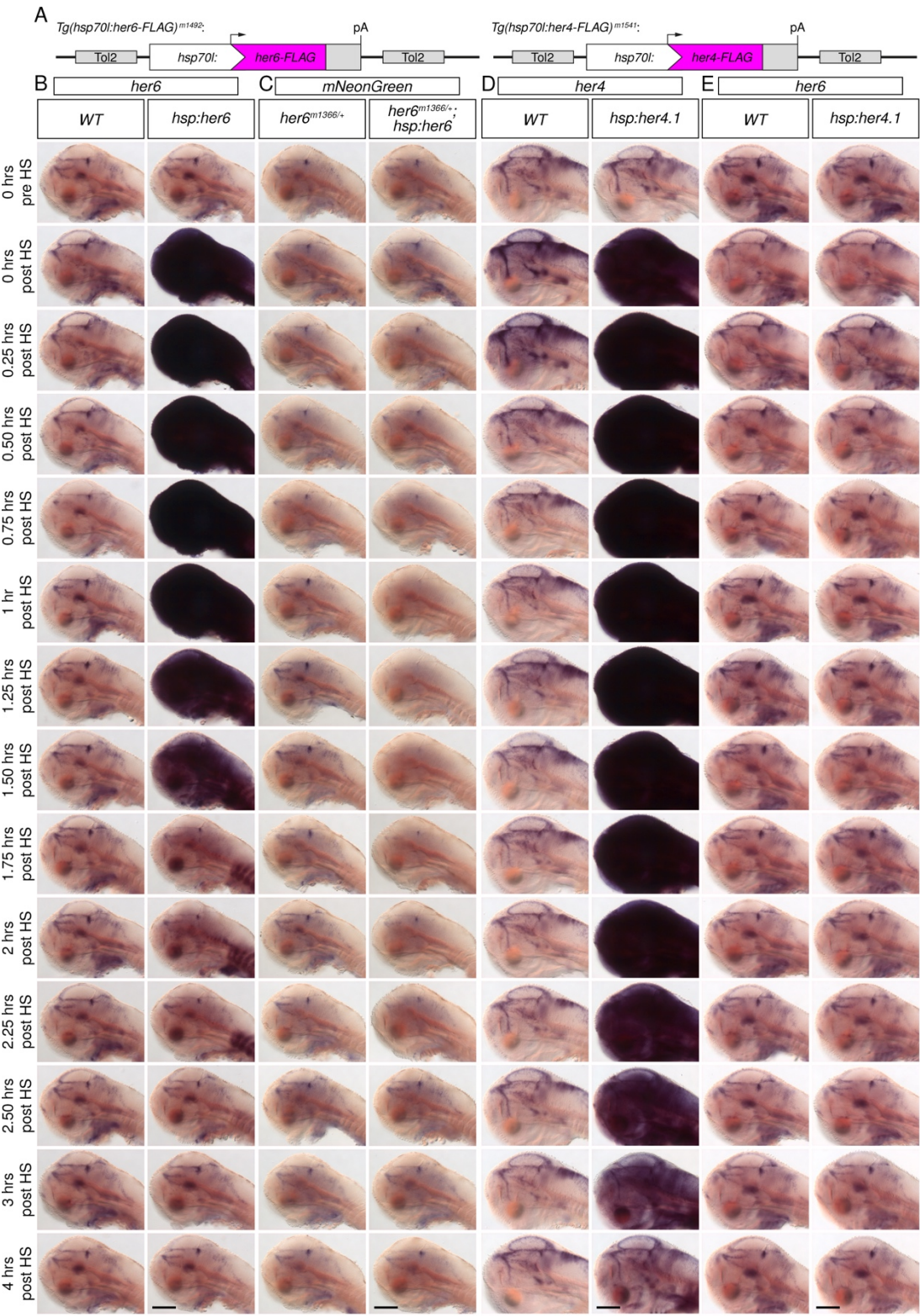

**Figure S11: Heat shock induced overexpression of Her6 downregulates *her6-mNeonGreen* expression, while overexpression of Her4 has no detectable effects on *her6* expression.**

(A) Schematic depiction of transgenes used for *her6* and *her4* overexpression respectively. (B - E) In situ hybridizations for *her6* (B, E), *mNeonGreen* (C) and *her4* (D). For (B) and (C), Her6 was overexpressed using the *Tg(hsp70l:her6-FLAG)<sup>m1492</sup>* line in heterozygous *Tg(her6:her6-mNeonGreen)<sup>m1366</sup>* embryos by a 15 min heat shock. For (D) and (E), Her4.1 was overexpressed using the *Tg(hsp70l:her4.1-FLAG)<sup>m1541</sup>* line by a 15 min heat shock. Time series analysis (B) – (E) was conducted at 15 min intervals, with time points of fixation indicated at left. WT indicates *her6*<sup>+/+</sup>, *her4*<sup>+/+</sup>. Scale bars: 200 μm.

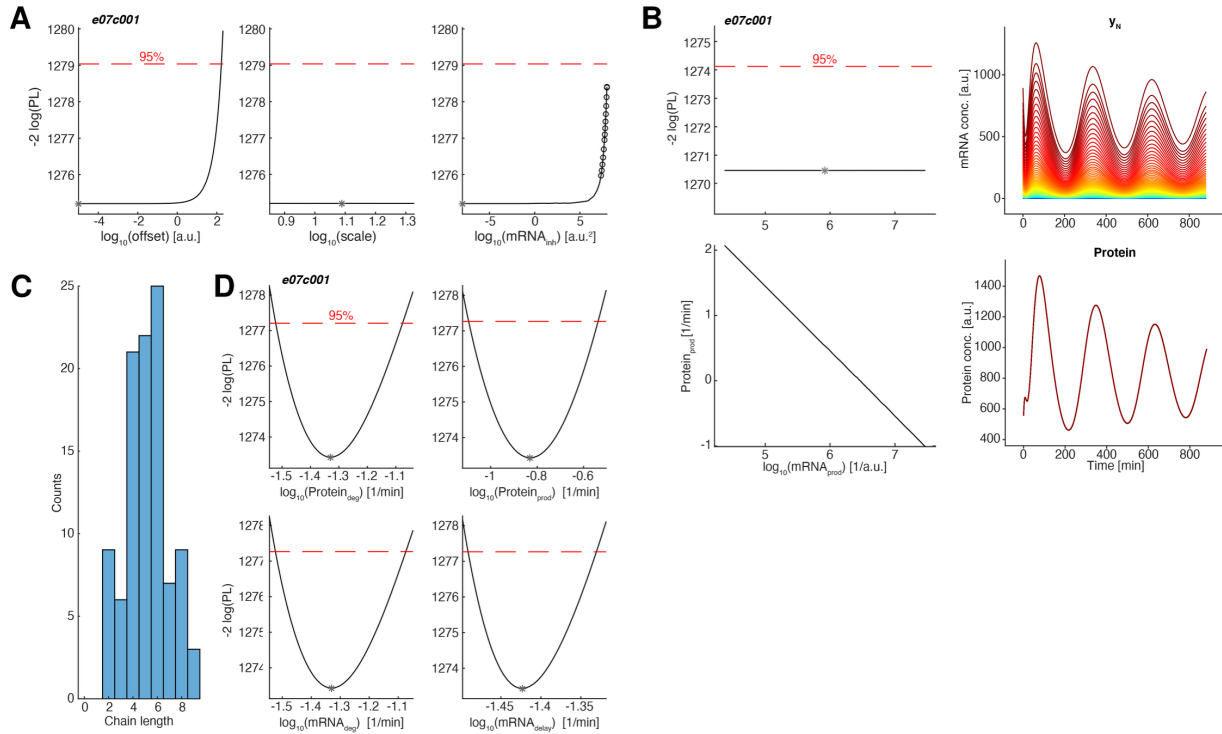

**Figure S12: Parameter estimation for the dynamic model of Her6 expression as well as comparison with $Protein_{deg}$ and $mRNA_{deg}$ values obtained directly from degradation measurement**

**A**, Profile likelihood of a subset of parameters of the non-reduced oscillation model (Equation 1, see Methods 18.2). Asterisks denote the best-fit value of the parameter. Circles denote points where a parameter hits the boundaries of parameter space after optimization. Confidence intervals of estimated parameters are given by values for which the profile likelihood PL (black) does not exceed the critical value corresponding to a significance level of 5% (dashed, red).

The parameters $offset$ and $mRNA_{inh}$ are practically non-identifiable; their confidence interval is unbounded towards negative infinity. The parameter $scale$ is structurally non-identifiable: Its confidence interval is unbounded in both directions, i.e. no matter what value is chosen for scale, the goodness-of-fit does not change. Therefore, we can reduce the model by fixing the parameters $offset$, $scale$ and $mRNA_{inh}$.

**B**, After performing the model reduction steps indicated in (A), the profile likelihood reveals a structural non-identifiability of $mRNA_{prod}$ (upper left panel). An increase in $mRNA_{prod}$ is compensated by a decrease of $Protein_{prod}$ (lower left panel) without changing the fit (lower right panel). However, unobserved model states such as $y_N$ (mRNA) are heavily affected by such changes (upper right panel): Each line corresponds to the model simulations for $y_N$ for a different value of $Protein_{prod}$, which is encoded by the color.

**C**, Histogram of optimal lengths of the linear chain in our model estimated from the individual time series of oscillating cells. For most time series, a chain length of four to six is optimal.

**D**, Profile likelihood of the relevant kinetic parameters contained in the reduced oscillation
model (Methods 18.2). Intersection points with the threshold value derived from a $\chi^2_I$-
328 distribution define the confidence intervals. All kinetic parameters are identifiable.

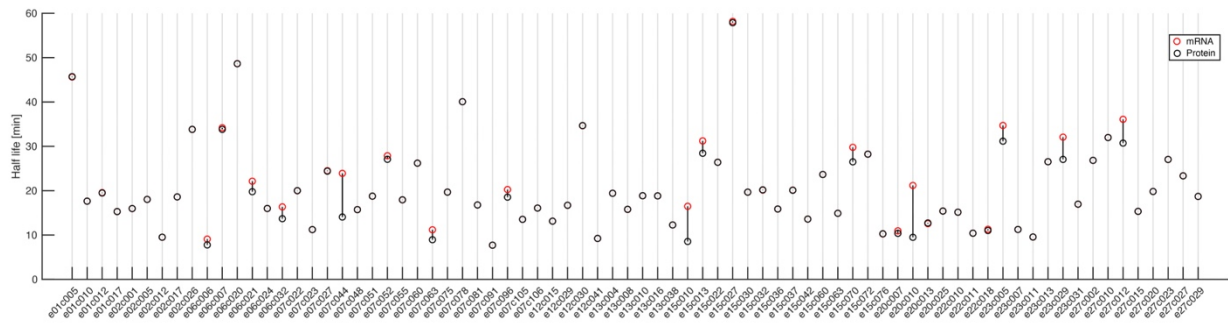

**Figure S13: Estimates of Her6 protein and *her6* mRNA stability based on mathematical model for each individual cell classified as oscillating.**

Graph shows half-life in min of *her6* mRNA (red) and Her6 protein (black) calculated for 79 stable oscillating cells based on the mathematical model (Figure 5A-G). Half-lives & degradation rate constants for *her6* mRNA and Her6 protein were derived from fitting the mathematical model (Figure 5A-G) to the Her6 protein time series data using maximum likelihood estimation, as illustrated in, e.g., Figure 5B for e07c001 and e15c042. This figure contains the resulting half-lives for time series data from 79 stable oscillating cells across 11 embryos. Most of the cells exhibit closely matching half-lives for mRNA and protein. This property facilitates stable oscillations, as shown in Figure 5F.

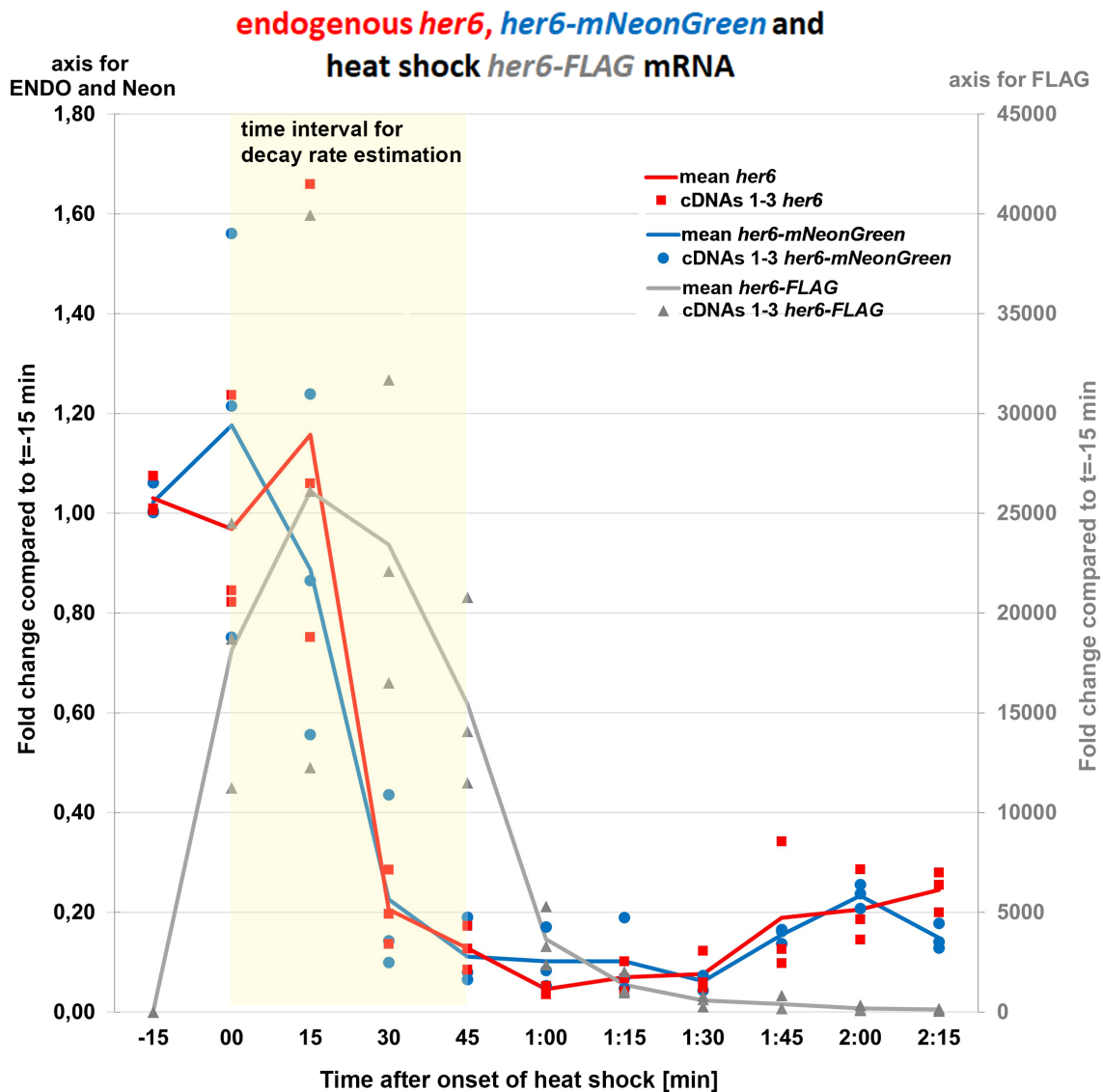

**Figure S14: Q-PCR quantification of *her6* levels after heat shock overexpression of *her6-FLAG*.**

Embryos from a $Tg(hsp70l:her6-FLAG)^{m1492}$ and $Tg(her6:her6-mNeonGreen)^{m1366}$ double transgenic cross were heat shocked for 15 min and samples collected in triplicates just before onset (-15 min) and at end of heat shock (00 min), and then every 15 min until 135 min post heat shock. cDNAs were synthesized (cDNAs 1-3) and expression levels of endogenous *her6* mRNA, *her6-mNeonGreen* mRNA, and heat shock overexpressed *her6-FLAG* quantified by Q-PCR. The plot shows fold-change in mRNA levels compared to the sample taking before start of heat shock (-15 min). Given the strong heat shock induction of *her6-FLAG*, a separate scale and y-axis is provided at right. The yellow area indicates the time interval for decay rate estimation.

357
358

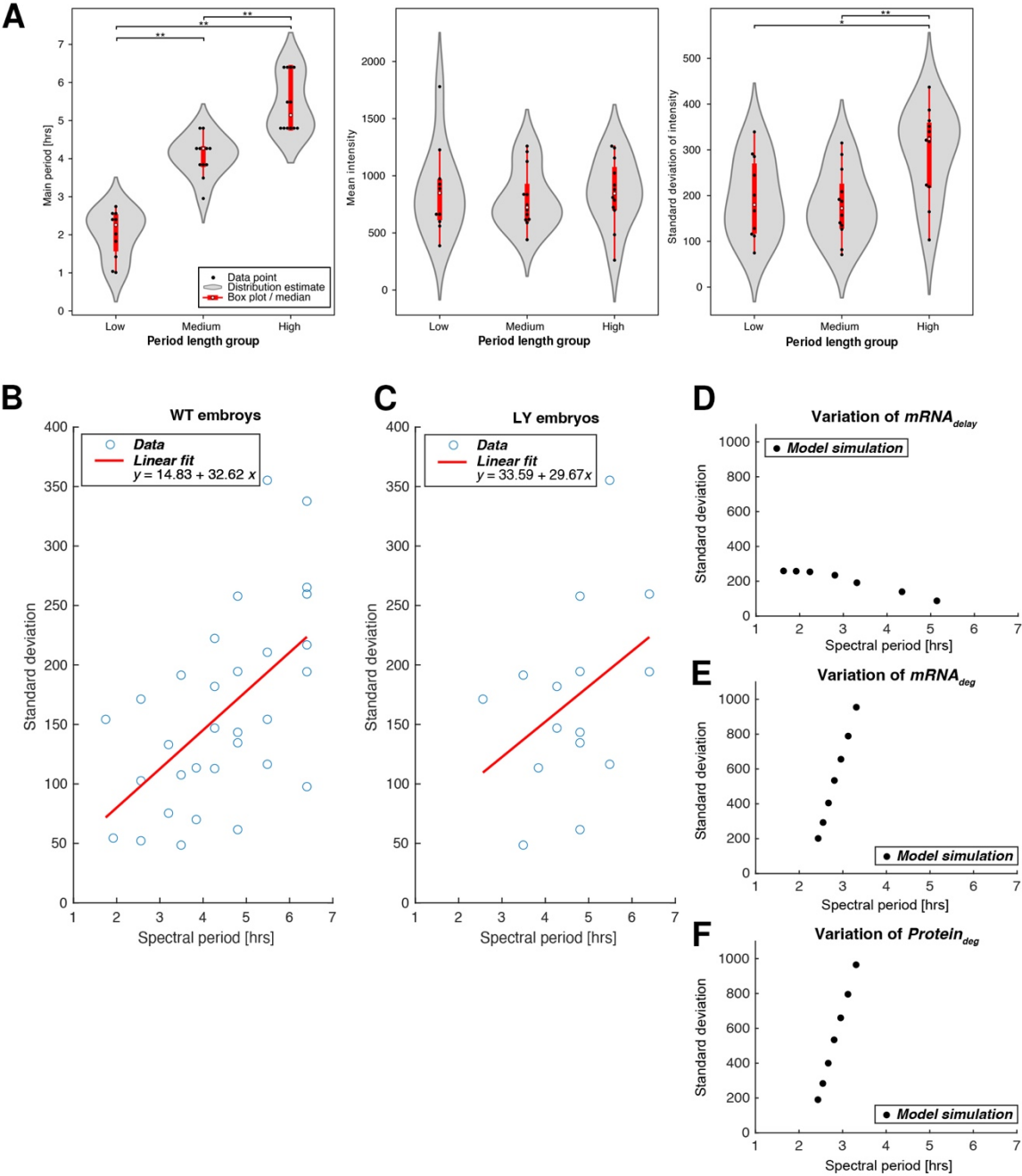

**Figure S15: Analysis of correlations between periods and amplitudes of oscillations.**

(A) Analysis of potential correlations of period length (left) with mean (middle) and standard deviation (right, as proxy for amplitude) of fluorescence intensities for cells of embryo 15. Analysis focussed on one embryo only as cells are recorded under comparable conditions.

Oscillating cells were equally distributed by period length into three equally sized groups (low, medium, high). \* $p \leq 0.05$, \*\* $p \leq 0.01$.

p-values for comparison Low versus Medium (L-M), Low versus High (L-H), Medium versus High (M-H)

Periods length

L-M $p=4.8e-05$; L-H $p=4.8e-05$; M-H $p=0.00013$

Mean intensity

L-M $p=0.71$; L-H $p=0.71$; M-H $p=0.45$

Standard deviation of intensity

L-M $p=0.80$; L-H $p=0.019$; M-H $p=0.0038$

**(B,C)** Standard deviations of time series show a correlation with the period calculated from the spectrum for wild-type embryos (B), as well as LY-treated embryos (C). We use the standard deviation as a measure for the amplitude of an oscillating mixing process. The red line depicts a linear fit of the data, where p-values for the slope coefficients are given by $p = 0.0008$ for WT, and $p = 0.14$ for LY treated embryos.

**(D-F)** With the best-fit parameters of e07c001, we simulate our mathematical model for the Her6 oscillator with different values for the kinetic rates $\text{mRNA}_{\text{delay}}$ (D), $\text{mRNA}_{\text{deg}}$ (E), and $\text{Protein}_{\text{deg}}$ (F), and calculate the standard deviation as well as the spectral periods of the model simulations.
